# Ubiquitin ligases and a processive proteasome facilitate protein clearance during the oocyte-to-embryo transition in *Caenorhabditis elegans*

**DOI:** 10.1101/2021.11.05.467479

**Authors:** Caroline A. Spike, Tatsuya Tsukamoto, David Greenstein

**Affiliations:** Department of Genetics, Cell Biology and Development, University of Minnesota, Minneapolis, Minnesota 55455, USA

**Author notes:** Corresponding author: David Greenstein, Department of Genetics, Cell Biology, and Development, University of Minnesota, 4-208 MCB, 420 Washington Avenue SE, Minneapolis, MN 55455.

**Keywords:** oocyte meiotic maturation, oocyte-to-embryo transition, translational regulation, RNA-binding proteins, ubiquitin-mediated protein degradation

## Abstract

The ubiquitin-mediated degradation of oocyte translational regulatory proteins is a conserved feature of the oocyte-to-embryo transition (OET). In the nematode *Caenorhabditis elegans*, multiple translational regulatory proteins, including the TRIM-NHL RNA-binding protein LIN-41/Trim71 and the Pumilio-family RNA-binding proteins PUF-3 and PUF-11, are degraded during the OET. Degradation of each protein requires activation of the M-phase cyclin-dependent kinase CDK-1, is largely complete by the end of the first meiotic division and does not require the anaphase promoting complex (APC). However, only LIN-41 degradation requires the F-box protein SEL-10/FBW7/Cdc4p, the substrate recognition subunit of an SCF-type E3 ubiquitin ligase. This finding suggests that PUF-3 and PUF-11, which localize to LIN-41-containing ribonucleoprotein particles (RNPs), are independently degraded through the action of other factors and that the oocyte RNPs are disassembled in a concerted fashion during the OET. We develop and test the hypothesis that PUF-3 and PUF-11 are targeted for degradation by the proteasome-associated HECT-type ubiquitin ligase ETC-1/UBE3C/Hu15, which is broadly expressed in *C. elegans*. We find that several GFP-tagged fusion proteins that are degraded during the OET, including fusions with PUF-3, PUF-11, LIN-41, IFY-1/Securin and CYB-1/Cyclin B, are incompletely degraded when ETC-1 function is compromised. However, it is the fused GFP moiety that appears to be the critical determinant of this proteolysis defect. These findings are consistent with a conserved role for ETC-1 in promoting proteasome processivity and suggest that proteasomal processivity is an important element of the OET during which many key oocyte regulatory proteins are rapidly targeted for degradation.

**Article Summary:** The ubiquitin-mediated degradation of translational regulatory RNA-binding proteins is a conserved feature of the oocyte-to-embryo transition (OET). *C. elegans* LIN-41 is a master regulator of oogenesis and is found in a large translational regulatory ribonucleoprotein (RNP) complex with more than 1000 maternal transcripts and the Pumilio-family RNA-binding proteins PUF-3 and PUF-11. We show that the concerted action of ubiquitin-mediated protein degradation and proteasome processivity rapidly disassemble LIN-41-containing RNPs during the OET thereby relieving repression of many maternal transcripts.

## INTRODUCTION

The oocyte-to-embryo transition (OET) initiates when oocytes exit meiotic prophase and enter the first meiotic metaphase, a cell cycle and developmental event also known as meiotic resumption or meiotic maturation. The OET is complete when zygotic gene transcription begins after fertilization in the early embryo. In many mammals and in the nematode *Caenorhabditis elegans*, zygotic gene transcription begins early at the 2- or ∼4-cell stage, respectively (Clegg and Pik6 1982; Latham *et al*. 1991; Edgar *et al*., 1994; Seydoux and Fire 1994; Hamatani *et al*. 2004). In some animals, such as Drosophila and Xenopus, zygotic gene activation predominantly begins later-at approximately the eighth nuclear division in Drosophila (Pritchard and Schubiger 1996; reviewed by Harrison and Eisen 2015) and at the midblastula transition in Xenopus (Bachvarova *et al*. 1966; Newport and Kirschner 1982a,b; reviewed by Blitz and Cho 2021). Because the late-stage oocytes of most animals are transcriptionally quiescent, the regulation of protein translation by RNA-binding proteins represents a dominant mode of developmental control during oocyte meiotic maturation and the OET (reviewed by Kotani *et al*. 2017; Huelgas-Morales and Greenstein 2018; Teixeira and Lehmann 2019). In fact, genomic analyses suggest that mammalian oocytes contain more than 200 RNA-binding proteins (Luong and Conti 2019). The presumption is that many of these RNA-binding proteins regulate protein translation in oocytes and preserve the dowry of maternal mRNA contributed to the early embryo.

In *C. elegans* oocytes, the TRIM-NHL RNA-binding protein LIN-41 is a key component of a large translational regulatory RNP complex that associates with more than 1000 mRNAs (Spike *et al*. 2014b; Tsukamoto *et al*. 2017). Many of these mRNAs encode protein components of LIN-41-containing RNPs and function as translational regulators during oogenesis or early embryonic development. LIN-41 also mediates the 3’-untranslated region (UTR)-mediated translational repression of some of the transcripts that associate with LIN-41 in oocytes and has an essential function to promote the extended meiotic prophase of oocytes (Spike *et al*. 2014a,b; Tsukamoto *et al*. 2017). *lin-41* null mutants are sterile because they produce pachytene-stage oocytes that exit meiotic prophase prematurely; these oocytes inappropriately disassemble the synaptonemal complex, cellularize, activate the CDK-1/cyclin B cyclin-dependent kinase, and enter M phase (Spike *et al*. 2014a). Hypomorphic *lin-41* mutant alleles reduce oocyte quality and cause nondisjunction during female meiosis (Spike *et al*. 2014a). LIN-41 prevents the premature activation of the CDK-1/cyclin B during early oogenesis, in part, through the 3’UTR-mediated translational repression of the CDC-25.3 phosphatase, a CDK-1 activator (Spike *et al*. 2014a,b). Premature CDK-1 activation in *lin-41* mutant oocytes drives precocious M-phase entry and causes oocytes to abnormally transcribe and express genes that are normally expressed zygotically after the OET (Allen *et al*. 2014; Spike *et al*. 2014a; Tocchini *et al*. 2014; Matsuura *et al*. 2016). Thus, a major function of LIN-41 appears to be to preserve mRNAs that are maternally inherited and translationally activated late in oogenesis or during early embryonic development. Many LIN-41-associated transcripts are translated only after LIN-41 is inactivated and degraded via ubiquitin-mediated degradation upon the onset of meiotic maturation (Spike *et al*. 2018). For example, LIN-41 associates with and represses the translation of the ORC-1 subunit of the origin recognition complex during meiotic prophase, and its degradation permits ORC-1 expression for DNA replication in the embryo (Tsukamoto *et al*. 2017).

LIN-41 is highly conserved and its mammalian ortholog, LIN-41/TRIM71, is required for embryonic viability and neural tube closure in mice (Maller Schulman *et al*. 2008; Chen *et al*. 2012; Cuevas *et al*. 2015; Mitschka *et al*. 2015). This defect appears to be related to a requirement for LIN-41/TRIM71 for maintaining pluripotency in neural progenitor cells in the embryo. LIN-41/TRIM71 has also been found to promote the pluripotency of embryonic stem cells in the mouse (Chang *et al*. 2012; Worringer *et al*. 2014; Mitschka *et al*. 2015; Liu *et al*. 2021). The defects in pluripotency, which underlie the neural tube defects observed in LIN-41/TRIM71 deficient mice, may be highly relevant to the finding of *de novo* heterozygous LIN-41/TRIM71 missense mutations in congenital hydrocephalus cases in humans (Furey *et al*. 2018; Jin *et al*. 2020). Germ cell-specific depletion of LIN-41/TRIM71 in mice causes germ cell loss and sterility in both sexes (Torres-Fernandez *et al*. 2021a). Interestingly, male mice with LIN-41/TRIM71-deficient germ cells exhibit a Sertoli cell-only phenotype, which resembles the infertility phenotype of human males with heterozygous LIN-41/TRIM71 mutations (Torres-Fernandez *et al*. 2021a). The essential role of LIN-41/TRIM71 in female germ cells remains to be more fully explored. Thus, in mammals, LIN-41/TRIM71 mutations produce distinct phenotypes in germ cells and somatic cells, which highlights the flexibility of LIN-41/TRIM71 proteins for deployment in developmental cell fate decisions. Indeed, LIN-41 was first discovered as a target of the *let-7* microRNA in the genetic pathway controlling the developmental timing of somatic cells in *C. elegans* (Reinhart *et al*. 2000; Slack *et al*. 2000). The regulation of LIN-41/TRIM71 orthologs by *let-7* microRNAs appears to be highly conserved in animals (O’Farrell *et al*. 2008; Kloosterman *et al*. 2004; Kanamoto *et al*. 2006; Lin *et al*. 2007; Rybak *et al*. 2009; Worringer *et al*. 2014). This mode of regulation is not observed in the *C. elegans* germline, as the *let-7* microRNA appears to be only expressed in somatic cells (Lau *et al*. 2001). A new twist is that recent studies suggest that mammalian LIN-41/TRIM71 can inhibit the accumulation of *let-7* microRNAs (Liu *et al*. 2021; Torres-Fernandez *et al*. 2021b), suggesting its involvement as a component of a bistable developmental switch. Although it is not known whether LIN-41 can inhibit the *let-7* pathway in *C. elegans*, it is clear that LIN-41 is a component of a bistable switch regulating oocyte meiotic maturation in this organism-LIN-41 inhibits CDK-1 activation, and in turn, active CDK-1 promotes the ubiquitin-mediated degradation of LIN-41 upon the onset of meiotic maturation (Spike *et al*. 2018).

The rapid elimination of LIN-41 upon the onset of meiotic maturation is emblematic of the highly choreographed and exquisitely timed patterns of degradation of RNA-binding proteins during late oogenesis and the OET in many organisms (Nishi *et al*. 2005; Stitzel *et al*. 2006; Shirayama *et al*. 2006; Sha *et al*. 2017; Kisielnicka *et al*. 2018; Cao *et al*. 2020, 2021; Zavortink *et al*. 2020; reviewed by Robertson and Lin 2015). Previous results suggested that an SCF-type E3 ubiquitin ligase containing the FBW7/Cdc4 ortholog SEL-10 promotes the degradation of LIN-41 using a bi-partite degron located near the N-terminus of LIN-41 (Spike *et al*. 2018). The ubiquitin proteasome pathway utilizes an E1-E2-E3 enzymatic relay to transfer ubiquitin to mark proteins for degradation by the 26S proteasome. E3 ubiquitin ligases, comprising multi-subunit enzymes or single chain proteins, function as substrate specificity factors to conjugate multiple ubiquitin moieties to target proteins. SEL-10 is a highly conserved and well-studied substrate-recognition subunit (Hubbard *et al*. 1997, de la Cova and Greenwald 2012). After substrate proteins are recognized and ubiquitinated, they are received by the 19S regulatory particles of the proteasome, which bind the ubiquitin chains, unfold the substrate, and processively translocate it into the 20S particle for degradation (reviewed by Finley 2009; Collins and Goldberg 2017).

In this work, we strengthen our understanding of LIN-41 degradation and begin to investigate how additional RNA-binding proteins in LIN-41 RNPs are degraded upon the onset of meiotic maturation. First, we show that SEL-10 is expressed in oocytes and use photoconversion of DENDRA::LIN-41 in maturing oocytes to demonstrate unequivocally that SEL-10 is required to degrade LIN-41 during the OET. Next, we identify the Pumilio/FBF-family RNA-binding proteins PUF-3 and PUF-11 as proteins that are degraded soon after meiotic maturation. PUF-3 and PUF-11 are components of LIN-41-containing RNPs in oocytes (Tsukamoto *et al*. 2017) that are 89 percent identical at the amino acid level and play important roles in germline stem cell and embryonic development (Hubstenberger *et al* 2012, Haupt *et al* 2020). Although both PUF-3 and PUF-11 are degraded soon after meiotic maturation in a CDK-1-dependent fashion, the degradation of both proteins occurs independently of SEL-10. Collectively, these results indicate that PUF-3 and PUF-11 are not degraded as a consequence of LIN-41 degradation and that LIN-41 RNPs are dismantled in a concerted fashion during the OET. Finally, we find that several different GFP-tagged fusion proteins that are degraded during the OET, including fusions with PUF-3, PUF-11 and LIN-41, are incompletely degraded when the function of the conserved ETC-1/UBE3C/Hu15 HECT-domain ubiquitin ligase is reduced, indicating that ETC-1 is important for proteasome processivity. We provide evidence that the GFP tag causes this proteolysis defect, re-evaluate key aspects of a previous report that assigned a different role to ETC-1 (Wang *et al*. 2013), and propose that proteasomal processivity is critical for the rapid and efficient targeting of a plethora of oocyte regulatory proteins for degradation during the OET.

## MATERIALS AND METHODS

### Strains and genetic analysis

The genotypes of strains used in this study are reported in Table S1. All analyses were conducted at 20 °C, unless specified otherwise. *mat-l(ax16l*ts) animals are susceptible to even brief observations and manipulations at room temperature. Thus, strains containing the *mat-l(ax16l*ts) mutation were handled as little as possible (< 15 min at room temperature) and prechilled (15°C) media was used to establish cultures. The following mutations were used: LGI-*rnp-8(tn1860[rnp-8::gfp::tev::3xflag])*, *mat-l(ax16l*ts), *lin-41(tn154l[gfp::tev::s::lin-41]))*, *lin-41(tn1892[mscarlet::tev::3xflag:lin-41])*, *lin-41(tn1894[dendra::tev::3xflag::lin-41])*, *lin-41(tn2054[gfp::tev::3xflag::lin-41])*, *lin-41(tn2055[gfp::tev::3xflag::lin-41])*, and *fog-3(q443)*. *lin-41(tn2054)* and *lin-41(tn2055)* were generated independently but are identical at the DNA sequence level. LGII-*dpy-10(tn2076)*, *etc-1(gk5182)*, *etc-1(tn1919[gfp::tev::3xflag::etc-1]), etc-1(tn1920[gfp::tev::3xflag::etc-1]), etc-1(tn2077),* and *unc-4(e120)*. LGIII-*plk-l(or683*ts*), emb-30(tn377*ts), and *cul-2(or209*ts). LGIV-*puf-ll(tn1824[puf-ll::gfp::tev::3xflag]), puf-ll(q97l)*, *cyb-l(tn1806[cyb-l::gfp::tev::3xflag]), unc-22(e66), puf-3(tn1820[puf-3::gfp::tev::3xflag]),* and *puf-3(q966)*. LGV-*lon-3(e2175)*, *sel-10(ar4l)*, *sel-10(ok1632)*, *sel-10(tn1816[sel-10::gfp::tev::3xflag])*, *sel-10(tn1817[sel-10::gfp::tev::3xflag])*, *sel-10(tn1875[gfp::tev::3xflag::sel-10])* and *fog-2(oz40)*. *sel-10(tn1816)* and *sel-10(tn1817)* were generated independently but are identical at the DNA sequence level. The following balancers were used: *tmC18[dpy-5(tmls1236)]* I (Dejima *et al*. 2018), *mnCl[dpy-10(e128) unc-52(e444) umnls32]* II, *nTl[qls5l]* (IV; V). The following transgene insertions were used: *itls37[pie-lp::mcherry::histoneH2B::pie-l 3’UTR, unc-119(+)]* IV (McNally *et al*. 2006) and *ddIs128[ify-1::2xty1::egfp::3xflag(92C12) + unc-119(+)]* (Sarov *et al*. 2012).

### mScarlet and Dendra repair templates

The repair templates used to create *lin-41(tn1892[mscarlet::tev::3xflag::lin-41])* and *lin-41(tn1894[dendra::tev::3xflag::lin-41])* were generated by Gibson assembly using the NEBuilder HiFi DNA Assembly master mix (New England Biolabs, Ipswich, MA) and five different PCR products: (1) a *lin-41* 5’ homology arm (∼0.6 kb), (2) a codon-optimized mScarlet-I or a Dendra coding sequence with introns (∼0.9 kb), (3) self-excising cassette (SEC)-containing sequences (∼5.7 kb), (4) a *lin-41* 3’ homology arm (∼0.8 kb), and (5) vector backbone sequences (∼2.6 kb). *lin-41* homology arms (1 and 4) were amplified from genomic DNA prepared from the wild type. SEC and vector sequences (3 and 5) were amplified from *Cla*I and *Spe*I-digested pDD268 plasmid DNA (Dickinson *et al*. 2015). mScarlet-I and Dendra were amplified from the plasmids pMS050 (Addgene plasmid #91826), a gift from Bob Goldstein, and pEG545 (Griffin *et al*. 2011), a gift from Geraldine Seydoux. Primer sequences are presented in File S1. All PCR products were initially column purified (Qiagen, Valencia, CA) and most of the plasmids recovered after assembly lacked the SEC fragment; only 10% and 5% of the mScarlet-I and Dendra repair constructs were complete, respectively. While making different *lin-41* repair constructs using the same method and many of the same PCR fragments, we found that gel purifying the SEC PCR product increased the inclusion of the SEC fragment to 80% of the assembled plasmids.

Constructs created by Griffin *et al*. (2011) that contain DENDRA-expressing sequences optimized for use in *C. elegans* (Gallo *et al*. 2010) are annotated in Addgene as containing DENDRA2 (e.g., pEG545, Addgene plasmid #40116 and pEG345, Addgene plasmid #40077). DENDRA2 is a modified form of DENDRA that has an important single amino acid change near the C-terminus (A224V) as well as a 5-amino acid insertion near the N-terminus (Gurskaya *et al*. 2006; Chudakov *et al*. 2007). The relationship between DENDRA and DENDRA2 is most easily visualized using the fluorescent protein database www.fpbase.org (Lambert 2019). We sequenced the pEG545 plasmid and confirmed that it encodes DENDRA; the DENDRA-coding sequence lacks both the A224V amino acid change and the 5-amino acid insertion found in DENDRA2. Since we obtained pEG545 directly from the Seydoux lab, we also analyzed the Addgene-deposited sequences of pEG545 and pEG345. Although those sequences are incomplete, they confirm that each plasmid encodes DENDRA[A224]. As both plasmids have been used to generate many different fusion proteins labeled as containing DENDRA2 [e.g., Herrera *et al*. (2016) and Rosu and Orna Cohen-Fix (2017)], this annotation error may be relatively common in the *C. elegans* literature.

### Genome editing

Plasmids that express guide RNAs (gRNAs) under the control of the U6 promoter were generated as described (Arribere *et al*. 2014). Repair templates used to tag genes with *gfp* were also generated as described (Dickinson *et al*. 2015). Repair templates used to tag genes with mSCARLET-1 and DENDRA were generated as described in the previous section. Genome editing was performed by injecting wild-type adult gonads with a DNA mix containing a repair template (10 ng/μl), one or more gRNA plasmids (25 ng/μl each), Cas9-expressing plasmid (pDD162, 50 ng/μl), and injection marker (pMyo2::Tdtomato, 4 ng/μl) and selecting for repairs and selection cassette excisions using standard methods (Dickinson *et al*. 2015). Null mutations in *lin-41* are both Dumpy (Dpy) and sterile (Slack *et al*. 2000; Spike *et al*. 2014a). None of the animals homozygous for the *mScarletASECAlin-41*, *DendraASECAlin-41* and *GFPASECAlin-41* repairs were Dpy, but some were sterile, consistent with the interpretation that the insertion of each SEC (self-excising cassette) at the 5’ end of the *lin-41* gene created reduction-of-function rather than null alleles of *lin-41*. Each *lin-41* repair was balanced with *tmC18* and the SEC was removed from the *lin-41* locus in balanced heterozygotes. Fluorescently tagged *lin-41* alleles were homozygous fertile after SEC excision. All edited loci were validated by sequencing the repair junctions using PCR products as templates. At least two alleles derived from independently injected parents were isolated and assigned allele names. A 4240 bp deletion in the *etc-1* locus, which removes the entire open reading frame was constructed using the *dpy-1*0 co-conversion method (Arribere *et al*. 2014). The injection mix contained pJA58 (7.5 ng/µl), AF-ZF-827 (500 nM), etc-1 sgRNA2 (pCS629) (25 ng/µl), etc-1 sgRNA4 (pCS633) (25 ng/µl), the repair template etc-1_RT1 (500 nM), and pDD162 (50 ng/µl). Correct targeting was verified by conducting PCR with primer pairs eif-2Bepsilon_F2 and hgap-2_F2 followed by Sanger sequencing. The DG5379 *dpy-10(tn2076) etc-1(tn2077)* strain was isolated after screening the progeny of 267 F1 Roller animals. The *etc-1(tn2077)* null mutation was separated from the *dpy-10(tn2076)* mutation by recombination by seeking non-Dpy non-Rol non-Unc animals from *dpy-10(tn2076) etc-1(tn2077)/unc-4(e120)* heterozygotes.

### Antibody staining

Dissected gonads stained with the rabbit anti-GLD-1 primary antibody (1:200 or 1:300 dilution; Jan *et al*. 1999) were fixed in 1% paraformaldehyde for 10 min, with a 5-min post fix step in ice-cold methanol as previously described (Rose *et al*. 1997; Spike *et al*. 2018). Primary antibodies were detected using Cy3-conjugated donkey anti-rabbit secondary antibodies (1:500 dilution; Jackson ImmunoResearch, West Grove, PA).

### Microscopy and image analysis

Microscope images in Figures 4 and S2 were acquired on a Carl Zeiss motorized Axioplan 2 microscope with a 63x Plan-Apochromat (numerical aperture 1.4) objective lens using an AxioCam MRm camera and AxioVision software (Carl Zeiss, Thornwood, NY). Microscope images in Figures S4, 5I-L, S6, SS and S9 were acquired on a Nikon Ni-E microscope with either a Plan Apo λ 60x (numerical aperture 1.4) objective or a Plan Fluor 40x Oil (numerical aperture 1.3) objective using NIS elements (Nikon Inc., Melville, NY). All the remaining microscope images were acquired on a Nikon Ti2 inverted confocal microscope with a Plan ApoIR 60x objective (numerical aperture 1.27), motorized stage and Galvano scanner using NIS elements (Nikon Inc., Melville, NY). When using the Nikon Ti2 microscope, the large image function of NIS elements was used to acquire images that show the entire germline. We quantified the average fluorescence intensity in selected areas of images using the measure function of ImageJ (Schneider *et al*. 2012). Each measurement was background corrected using the average fluorescence intensity of a nearby area outside the body of the worm. Multiple images were analyzed per genotype, as specified in the text.

### Photoconverting ^Green^DENDRA::LIN-41 to ^Red^DENDRA::LIN-41

Worms were mounted without anesthetic on 10% agarose pads in a 1:1 solution of M9 buffer and 0.1 μM polystyrene beads (Polysciences, Inc., Warrington, PA) essentially as described (Rosu and Cohen-Fix 2020). Photoactivation and imaging was performed using the Nikon Ti2 inverted confocal microscope equipped as described in the previous section. A region of interest encompassing the −1 oocyte was selected for photoactivation using the 405 nm laser at 10% power for 31.25 seconds (125 loops at 4 frames per second) and a large pinhole (173.69 microns) to promote DENDRA photoconversion throughout the oocyte. After photoactivation, animals were allowed to recover on NGM plates seeded with OP50-1 bacteria for either 60 min or 100 min and then mounted a second time. Confocal settings optimized to image oocytes after photoconversion were used to collect images of the ^Green^DENDRA::LIN-41 and ^Red^DENDRA::LIN-41 in oocytes and embryos at all stages of the experiment: before photoactivation, after photoactivation, and after recovery.

Although the conditions used for photoactivation were generally compatible with oocyte meiotic maturation and ovulation, our experimental manipulations sometimes interfered with these events. For example, one *lin-41(tn1894)* animal failed to move after recovery, suggesting that it might have been damaged while mounting on or recovering from the agarose pad. ^Red^DENDRA::LIN-41 was still detectable in the −1 and −2 oocytes of the photoconverted arm after a 100-min recovery period, indicating that the converted oocytes in this damaged animal had not matured or ovulated. The other *lin-41(tn1894)* animals (n=5) survived the experimental manipulations and exhibited exclusively green oocytes in the −1 and −2 positions in each photoconverted arm (n=6 arms, the −1 oocytes of both arms were converted in one animal), confirming that oocytes had matured and been ovulated into the uterus during the recovery period (60 min (n=2) or 100 min (n=4)). All *lin-41(tn1894); sel-10(ok1632)* animals survived our experimental manipulations (n=4 animals), and most of the photoconverted arms had exclusively green oocytes in the −1 and −2 positions (n=4/5 arms, the −1 oocytes of both arms were converted in one animal) and one or two red embryos in the uterus after the recovery period. The sole exception was in an animal that laid only 2 eggs during the 100-min recovery period and appeared to be sickly. This animal had a bright red 1-cell stage embryo in the +1 position and a fainter red oocyte in the −1 position. The −1 oocyte converted in this gonad arm must have experienced a substantial temporal delay prior to oocyte maturation and ovulation, as a shorter (60 min) recovery period was sufficient for two ovulation events in the healthier *lin-41(tn1894); sel-10(ok1632)* animals.

### itls37 puf-3(tn1820) and puf-ll(tn1824) puf-3(tn1820) double mutants

Crosses with *puf-ll(tn1824)* and *puf-3(tn1820)* established that the *it!s37[pie-lp::mcherry::histoneH2B::pie-l 3’UTR, unc-119(+)}* insertion on chromosome IV is tightly linked to *puf-ll* and more loosely linked to *puf-3*. 0 of 72 progeny from *it!s37 + I + puf-ll(tn1824)* parents carried a recombinant *it!s37 puf-ll(tn1824)* chromosome, while 5 of 64 progeny from *it!s37 + I + puf-3(tn1820)* parents carried a recombinant *it!s37 puf-3(tn1820)* chromosome. Together, these observations suggested that *it!s37* is ∼7.8 map units to the left of *puf-3*, in good agreement with the relative genetic positions of *puf-ll* and *puf-3* described in WormBase (http://www.wormbase.org, release WS278; Davis *et al*. 2022). We utilized *unc-22(e66)*, which causes a homozygous Uncoordinated (Unc) phenotype and is located in-between *puf-ll* and *puf-3*, to construct the *puf-ll(tn1824) puf-3(tn1820)* double mutant. Animals carrying *it!s37 puf-3(tn1820)* and *puf-ll(tn1824) unc-*22*(e66)* recombinant chromosomes were mated with each other. Animals with one copy of each chromosome were selected and allowed to self-fertilize. In the next generation, non-Unc-22 animals were examined for *it!s37*-derived mCHERRY::HISTONE expression using fluorescence microscopy A single *it!s37*-negative animal was identified among the 168 non-Unc animals examined, and this animal segregated approximately one quarter Unc progeny in the next generation. Based on the mapping data, we anticipated that this rare animal carried either a *puf-ll(tn1824) puf-3(tn1820)* or a *puf-ll(+) puf- 3(tn1820)* recombinant chromosome in *trans* to the parental *puf-ll(tn1824) unc-22(e66)* chromosome. Animals homozygous for the recombinant chromosome were analyzed using primers that produce allele-specific PCR products (File S1) and found to carry both *puf-ll(tn1824)* and *puf-3(tn1820)*.

### RNA interference

Gene-specific RNA interference (RNAi) was performed by feeding *C. elegans* with double-stranded RNA (dsRNA)-expressing *E. coli* (Timmons and Fire 1998) at 22 °C using the RNAi culture media described by Govindan *et al*. (2006). RNAi clones were obtained from Source BioScience (Nottingham, UK), and the identities of important RNAi clones were verified by DNA sequencing (see File S1 for details). Exposure to dsRNA-expressing *E. coli* was initiated during the fourth larval stage and animals were examined and imaged after 2 days. Under these RNAi conditions and at the time-point analyzed, all *cdk-l(RNAi)* animals produced arrested 1-cell embryos and therefore had a strong *cdk-l(RNAi)* phenotype (see Boxem *et al*. 1999). Using the same RNAi conditions, the gonad arms of *wee-l.3(RNAi)* animals exhibited a range of oocyte phenotypes. Gonads with a mild *wee-l.3(RNAi)* phenotype have relatively large and normal-appearing oocytes that occupy approximately normal positions with respect to each other and to the spermatheca. Gonads with strong *wee-l.3(RNAi)* phenotype have small, sometimes indistinct oocytes or a mass of oocyte-like material near the spermatheca. Burrows *et al*. (2006) also noted that prolonged RNAi results in small oocytes, consistent with the idea that this phenotype is caused by a strong response to *wee-l.3(RNAi)*. GFP_S_::LIN-41 is prematurely degraded in animals with both weak and strong *wee-l.3(RNAi)* phenotypes (Spike *et al*. 2014a, 2018). In the screen for E3 ubiquitin ligases that degrade PUF-3::GFP, we did not exclusively use RNAi to test the SCF subunit CUL-2. Since our *cul-2(RNAi)* clone does not trigger a good RNAi response (Spike *et al*. 2018), we also examined *cul-2(or209*ts); *puf-3(tn1820[puf-3::gfp::tev::3xflag])* adults 1 day after they were upshifted to 25 °C as 14-stage larvae. All other ubiquitin ligase subunits were screened using RNA interference alone.

### Western blots

Each lane was loaded with a protein lysate made from adult worms picked into 10 microliters of phosphate-buffered saline (PBS) containing proteinase inhibitors (2x complete EDTA-free; Millipore-Sigma, St. Louis, MO). Whole-worm lysates were frozen at -80 °C and prepared for electrophoresis by adding LDS sample buffer supplemented with a reducing agent (Invitrogen, Carlsbad, CA). Immediately after adding the sample buffer, the samples were heated at 70 °C and vortexed multiple times prior to loading on a gel. Lysates were made from 40 Day 1 adult worms (24 hours past the 14 stage at 20 °C) with a few exceptions. Some lysates expressing CYB-1::GFP were made from 50 instead of 40 worms (hermaphrodites in Figures 7A and S7B; males in Figures 7B and S7D). In addition, the mated females used for lysates were not developmentally staged (Figure S7C). Instead, gravid females were selected from mixed-stage plates. Proteins were separated using 3-8% Tris-Acetate gels (Invitrogen, Carlsbad, CA) and visualized after western blotting. Blots were blocked with 5% nonfat dried milk. The primary antibodies used were mouse anti-FLAG M2 (Millipore-Sigma) at a 1:2,000-1:4,000 dilution and rabbit anti-GFP NB600-308 (Novus Biologicals, Littleton, CO) at a 1:4,000 dilution. The secondary antibodies used were peroxidase-conjugated goat anti-mouse (Thermo Scientific, Waltham, MA) at a 1:30,000 dilution or peroxidase-conjugated donkey anti-rabbit (Thermo Scientific) at a 1:5,000 dilution. Detection was performed using SuperSignal West Femto Maximum Sensitivity Substrate and CL-XPosure film (Thermo Scientific).

### eGFP, mSCARLET-1 and DENDRA fusion protein tag sequences

The predicted amino sequence and molecular mass of each protein tag is presented in File S2 for reference. All GFP tags express the GFP[S65C] enhanced GFP variant commonly used in *C elegans* (Green *et al* 2008). Most also contain the apparently neutral GFP[Q80R] change likely introduced into the original GFP cDNA sequence by PCR error (Tsien 1998).

### Data availability

All alleles and strains (Table S1), plasmids (File S1) and Sanger sequencing files are available upon request. Primer sequences are provided in File S1. File S2 contains the amino acid and nucleotide sequences of fluorophore tags, tagged alleles, and deletion alleles created for or used in this work. Supplemental materials are available at Figshare: https://doi.org/

## RESULTS

### SEL-10 is present in the cytoplasm and nucleoplasm of oogenic germ cells

SEL-10/Cdc4/FBW7 is a highly conserved F-box protein; it is the substrate-recognition subunit of a Skp, Cullin, F-box (SCF)-containing E3 ubiquitin ligase complex that promotes the ubiquitination and degradation of specific substrates (reviewed in Deshaies and Ferrell 2001; Welcker and Clurman 2008). In *C. elegans*, SEL-10 functions in the oogenic germline and during the OET to promote the timely elimination of the cytoplasmic RNA-binding proteins GLD-1 and LIN-41 at different stages of oogenesis (Kisielnicka *et al*. 2018; Spike *et al*. 2018). Consistent with this idea, *sel-10* mRNA is expressed throughout the oogenic germlines of adult hermaphrodites and the *sel-10* 3’ UTR is permissive for the translation of green fluorescent protein (GFP) in pre-meiotic and meiotic cells during oogenesis (Kisielnicka *et al*. 2018). However, the expression of SEL-10 has not been reported in oocytes or in any other region of the *C. elegans* germline, possibly because SEL-10 expression constructs were examined in the context of extrachromosomal arrays (Ding *et al*. 2007; Dorfman *et al*. 2009), which can silence the expression of germline genes in *C. elegans* (Kelly *et al*. 1997). To avoid this technical issue, we used genome editing to generate new alleles of *sel-10* that have a GFP-tag fused to the SEL-10 protein at both the amino-terminus (N-terminus, GFP_F_::SEL-10 in Figure 1) and carboxyl-terminus (C-terminus, SEL-10::GFP_F_ in Figure 1). In the figures, and where needed for textual clarity, we will indicate the presence and relative position of a 3xFLAG or an S epitope tag in a fusion protein with an F or S subscript, respectively. The alleles that express GFP::SEL-10, SEL-10::GFP, and many of the other fusion proteins used in this work, are shown in Figure S1.

**Figure 1.**
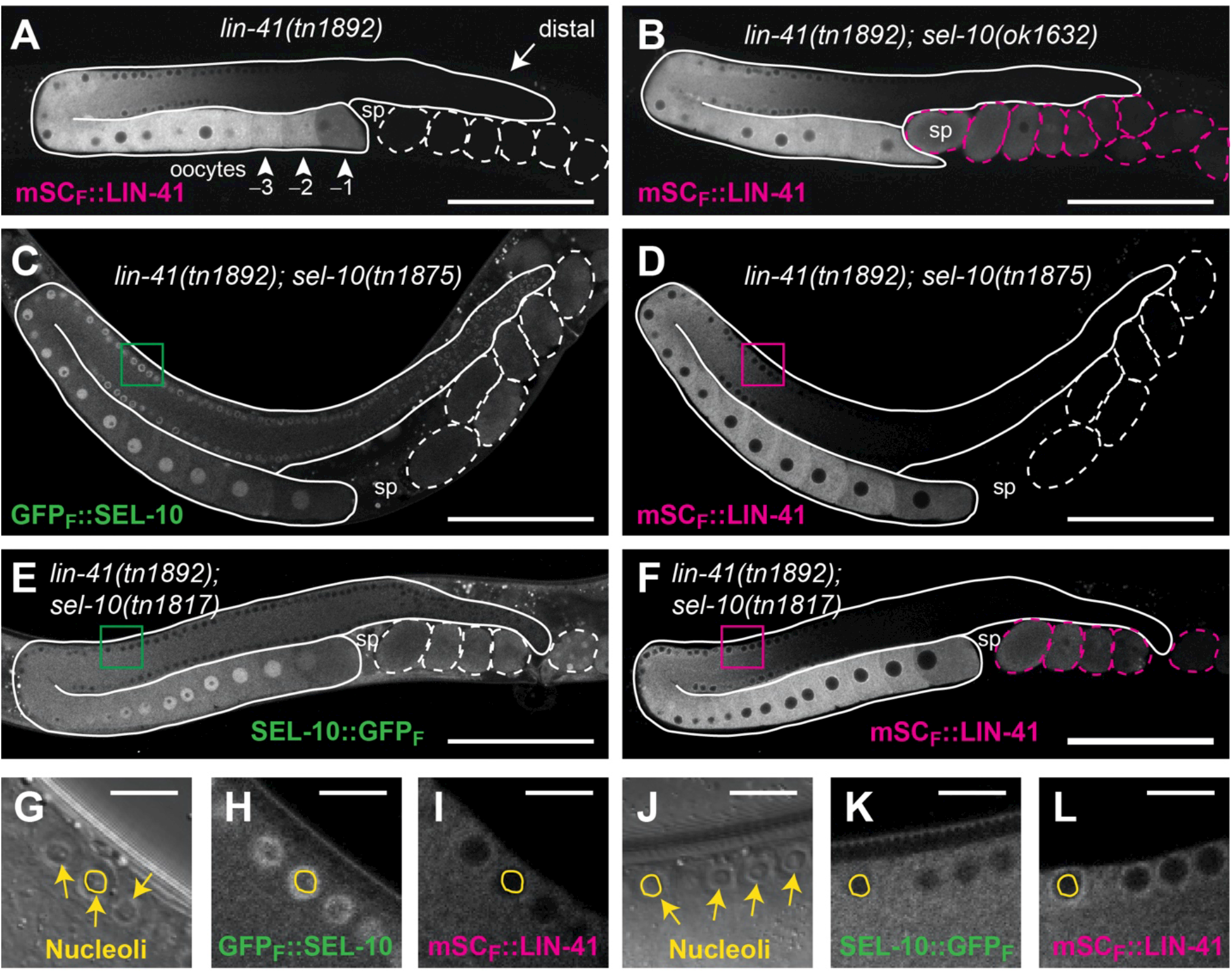
SEL-10 is expressed in oocytes and embryos and promotes the degradation of mSCARLET::LIN-41 during the OET. mSCARLET::LIN-41 (A, B, D, F, I, and L), GFP::SEL-10 (C and H) and SEL-10::GFP (E and K) fluorescence in *C. elegans* germlines (solid outlines) and early embryos (dashed outlines). mSCARLET::LIN-41 is rapidly degraded after oogenesis in the wild type (A) and *sel-1O(tn1875[gfp::3xflag::sel-1O])* (D) hermaphrodites but persists in early embryos in the *sel-1O(ok1632)* deletion (B) and *sel-1O(tn1817[sel-1O::gfp::3xflag])* (F) hermaphrodites. Embryos with an increased amount of mSCARLET::LIN-41 relative to *sel-1O(+)* embryos are outlined in magenta (B and F). Magnified image insets of the selected regions (C-F) are shown for GFP::SEL-10 (G-I) and for SEL-10::GFP (J-L), respectively. Differential interference contrast microscopic images are shown (G and J) to highlight germ cell nucleoli (yellow arrows). The circled nucleoli (G-L) lie in the middle focal plane of a germ cell nucleus, illustrating that functional GFP::SEL-10 is enriched in the nucleoplasm but largely excluded from nucleoli (G-I). Non-functional SEL-10::GFP is also excluded from nucleoli but is not enriched in the nucleoplasm relative to the cytoplasm at this stage of oocyte development (J-L). mSCARLET::LIN-41 is exclusively cytoplasmic and excluded from nuclei (I and L). Landmark structures such as the distal mitotic region of the germline (arrow in A), the proximal-most oocytes (arrowheads in A) and the spermatheca (sp), are indicated for reference. Scale bars, 100 μm (A-F) and 10 μm (G-L), respectively.

The pattern of GFP expression from each *sel-10* allele confirms that SEL-10 is expressed throughout the oogenic germline of adult hermaphrodites and in early embryos (Figure 1, C and E). GFP::SEL-10 and SEL-10::GFP are present in the nucleoplasm and cytoplasm of oocytes and distal germ cells, but show different patterns of localization, likely reflecting the fact that GFP::SEL-10 is functional, whereas SEL-10::GFP is not, as described in the subsequent section. GFP::SEL-10 is enriched in the nucleoplasm relative to the cytoplasm at all stages of germ cell development (Figure 1C). GFP::SEL-10 localization throughout the distal region of the gonad appears to be “ring-like” because it is enriched in the nucleoplasm relative to the cytoplasm and largely excluded from nucleoli (Figure 1, C and G-I). SEL-10::GFP is also excluded from nucleoli at all stages of germ cell development (Figure 1, E and J-L). However, SEL-10::GFP is present at similar levels in the nucleoplasm and cytoplasm of distal germ cells and only becomes enriched in the nucleoplasm of growing oocytes (Figure 1E). Using confocal images collected with identical settings, we measured 23% and 43% increases in background-corrected cytoplasmic fluorescence in the rachis and the −3 oocyte, respectively, in SEL-10::GFP animals relative to GFP::SEL-10 animals (*P*<0.01, n=7 images per genotype). However, nuclear fluorescence in the -3 oocyte of each strain was not significantly different (*P*>0.1). We conclude that, at least in the oogenic germline, SEL-10::GFP appears to be more abundant than GFP::SEL-10 and more likely to reside in the cytoplasm.

### GFP::SEL-10 is functional but SEL-10::GFP is not

The different patterns of GFP::SEL-10 and SEL-10::GFP localization might reflect the disruption or alteration of SEL-10 function by the inserted protein tags. We examined the accumulation of LIN-41 and GLD-1 in animals homozygous for each GFP-tagged allele of *sel-1O* to address this possibility. LIN-41 was monitored using *lin-41(tn1892[mscarlet::3xflag::lin-41})*, a new allele of *lin-41* that produces a fusion protein with an mSCARLET-tag fused to the N-terminus of LIN-41 (Figure S1B; mSC_F_::LIN-41). Endogenous GLD-1 levels were analyzed by immunostaining. mSCARLET::LIN-41 is strongly expressed in wild-type and *sel-1O(ok1632)* loss-of-function mutant oocytes (Figure 1, A and B). mSCARLET::LIN-41 is down-regulated in wild-type embryos (Figure 1A), but persists at elevated levels in the embryos produced by *sel-1O(ok1632)* mutant animals (Figure 1B). The downregulation of mSCARLET::LIN-41 protein expression during the OET is therefore SEL-10-dependent, as described for endogenous LIN-41 and a GFP-tagged LIN-41 fusion protein (Figure S1B; GFP_S_::LIN-41; Spike *et al*. 2018). mSCARLET::LIN-41 is downregulated in the embryos of GFP::SEL-10-expressing animals (*sel-1O(tn1875)*, Figure 1D), but persists at elevated levels in the embryos of SEL-10::GFP-expressing animals (*sel-1O(tn1817)*, Figure 1F). Similar results were observed with GLD-1 (Figure S2). GLD-1 is downregulated as developing oocytes exit pachytene near the loop region (Figure S2, A and C), and this process is impaired in *sel-1O(lf)* mutants, resulting in elevated levels of GLD-1 in proximal oocytes (Kisielnicka *et al*. 2018; Spike *et al*. 2018). GLD-1 is downregulated in the oocytes of GFP::SEL-10-expressing animals (*sel-1O(tn1875)*, Figure S2B), but persists at elevated levels in the oocytes of SEL-10::GFP-expressing animals (*sel-1O(tn1816)*, Figure S2D). We conclude that at least two of the normal functions of SEL-10 are compatible with the insertion of a GFP tag at the N-terminus of SEL-10 but are disrupted by the insertion of a GFP tag at the SEL-10 C-terminus. Furthermore, these observations suggest that the nucleoplasm-enriched localization pattern of GFP::SEL-10 shown in Figure 1C likely represents the correct localization pattern of SEL-10 in the oogenic germline. Although known targets of SEL-10 in the *C. elegans* germline are predominantly cytoplasmic (e.g., LIN-41, GLD-1 and CPB-3 (Kisielnicka *et al*. 2018; Spike *et al*. 2018)), orthologs of SEL-10 are present in the nucleoplasm in *S. cerevisiae* (Cdc4p) (Choi *et al*. 1990) and in both the nucleoplasm and cytoplasm in humans (FBW7α and FBW7β isoforms, respectively) (Welcker *et al*. 2004).

### SEL-10 is required to degrade LIN-41 during the OET

LIN-41 disappears rapidly upon the onset of meiotic maturation. We previously hypothesized that LIN-41 is phosphorylated at this stage and recognized by a SEL-10-containing E3 ubiquitin ligase that targets LIN-41 for proteolysis (Spike *et al*. 2018). To confirm that SEL-10 promotes the degradation of LIN-41 during the OET, we generated *lin-41(tn1894[dendra::3xflag::lin-41})*, a new allele of *lin-41* that produces a protein with a DENDRA-tag fused to the LIN-41 N-terminus (Figure S1B; DND_F_::LIN-41) and crossed it into a *sel-10(ok1632)* mutant background. Photoactivation irreversibly converts the fluorescence emitted by DENDRA from green to red (^Green^DENDRA and ^Red^DENDRA, respectively) (Gurskaya *et al*. 2006). The ^Green^DENDRA::LIN-41 present in the −1 oocytes of *sel-10(ok1632)* and *sel-10(+)* animals was targeted for conversion to ^Red^DENDRA::LIN-41 using a 405 nm laser. Robust photoconversion was observed in the −1 oocyte, with a small amount of red fluorescence sometimes visible in the adjacent cytoplasm of the −2 oocyte (Figure 2, A, B, F, and G). After a suitable recovery period to permit oocyte maturation, ovulation, and some embryonic development, bright ^Red^DENDRA::LIN-41 was evident in a single embryo in the uterus of *sel-10(ok1632)* animals (Figure 2, C-E; n=5/5). In general, we noted that the next-youngest embryo (or −1 oocyte in one animal, see Materials and Methods) also contained ^Red^DENDRA::LIN-41 (n=4/5), but at a much lower level. We did not observe bright ^Red^DENDRA::LIN-41 in any of the embryos in the uteri of *sel-10(+)* animals (n=6), although we sometimes saw an embryo with a slight haze of red fluorescence that we hypothesized might be derived from the converted ^Red^DENDRA::LIN-41 −1 oocyte (Figure 2, H and I). Relative to these *sel-10(+)* embryos, we measured a 6- to 10-fold higher level of background-corrected ^Red^DENDRA::LIN-41 fluorescence in the brightest *sel-10(ok1632)* embryos (n=4 of each genotype). Together, these results indicate that the fluorescently tagged ^Red^DENDRA::LIN-41 observed in *sel-10(ok1632)* embryos originates in oocytes and confirm that LIN-41 is degraded during the OET in wild-type animals.

**Figure 2.**
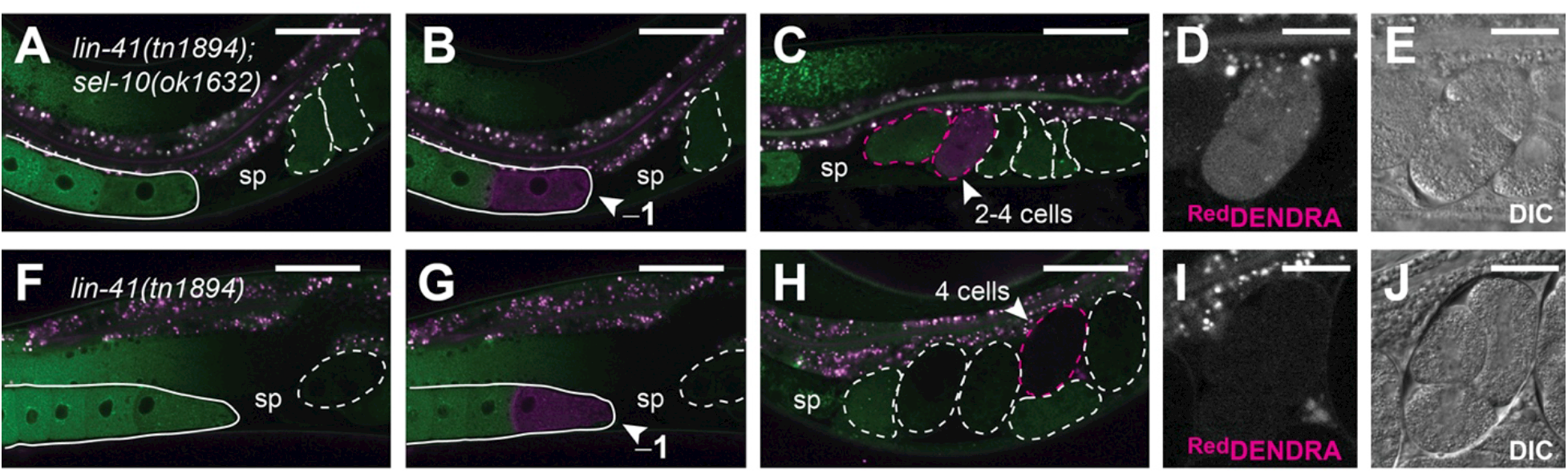
SEL-10 degrades oocyte-expressed DENDRA::LIN-41. Red and green channel fluorescence (A-C and F-H) in DENDRA::LIN-41-expressing *sel-1O(ok1632)* (A-E) and *sel- 10(+)* (F-J) hermaphrodites. The positions of oocytes (solid outlines), embryos (dashed outlines) and spermathecae (sp) are indicated for reference. Images were collected either before photoactivation (A and F), immediately after photoactivation of the −1 oocyte (arrowheads, B and G) or after a 100-min recovery period (C-E and H-J). Embryos with above-background levels of red channel fluorescence are outlined in magenta; the embryo that most likely developed from the photoactivated −1 oocyte is indicated with an arrowhead (C and H). Magnified red channel fluorescence (D and I) and DIC images (E and J) of this embryo are shown. Scale bars, 50 μm (A-C and F-H) and 20 μm (D, E, I, and J).

### PUF-3 and PUF-11 are degraded during the OET

PUF-3 and PUF-11 are nearly identical Pumilio-family RNA-binding proteins that function redundantly with respect to normal embryonic development in *C elegans* (Hubstenberger *et al* 2012; Haupt *et al* 2020). PUF-3 and PUF-11 proteins and mRNAs co-purify with LIN-41-containing ribonucleoprotein (RNP) complexes (Tsukamoto *et al* 2017). Furthermore, LIN-41 and PUF-3/11 were each identified as factors important for the 3’UTR-mediated translational repression of the Rbfox-related RNA-binding protein SPN-4 in oocytes (Hubstenberger *et al* 2012; Tsukamoto *et al*. 2017). These observations suggest that LIN-41, PUF-3, and PUF-11 function together in developing oocytes. We confirmed that PUF-3 and PUF-11 are expressed in oocytes, as described (Haupt *et al* 2020), and also examined the expression patterns of PUF-3 and PUF-11 during the oocyte-to-embryo transition. We created new alleles of *puf-3* and *puf-11* that fuse a GFP-tag to the C-terminus of each PUF protein (Figure S1C; PUF-3::GFP_F_ and PUF-11::GFP_F_) and found that PUF-3 and PUF-11 are eliminated during the OET in a manner that resembles LIN-41 (Figure 3). PUF-3::GFP and PUF-11::GFP are abundant in oocytes, but the expression of each protein is strongly diminished soon after oocyte meiotic maturation and ovulation (Figure 3, A and D; Figure S3, A and B). Consequently, PUF-3::GFP and PUF-11::GFP appear to be largely absent from embryos (Figure 3A) or limited to a single young embryo (Figure 3D and Figure S3, A and B). A similar pattern is observed with GFP::LIN-41 and mSCARLET::LIN-41 (Spike *et al*. 2018; Figure 1A). Moreover, PUF-3::GFP or PUF-11::GFP and mSCARLET::LIN-41 identify the same young embryos (if present, as in Figure 3, D and E; n2:7 animals of each genotype imaged as shown). To determine when PUF-3 is eliminated from embryos, we examined PUF-3::GFP in oocytes and embryos that express an mCHERRY::HISTONE fusion protein. Relative to the adjacent −1 oocyte, PUF-3::GFP levels are substantially reduced in meiotic embryos that are in metaphase I (Figure 3, G-J) and most PUF-3::GFP appears to be eliminated by the end of the first meiotic division (Figure 3K). This roughly corresponds to the time window during which LIN-41 is degraded; time-lapse imaging previously demonstrated that GFP::LIN-41 is dramatically reduced by the end of the first meiotic division (Spike *et al*. 2018).

**Figure 3.**
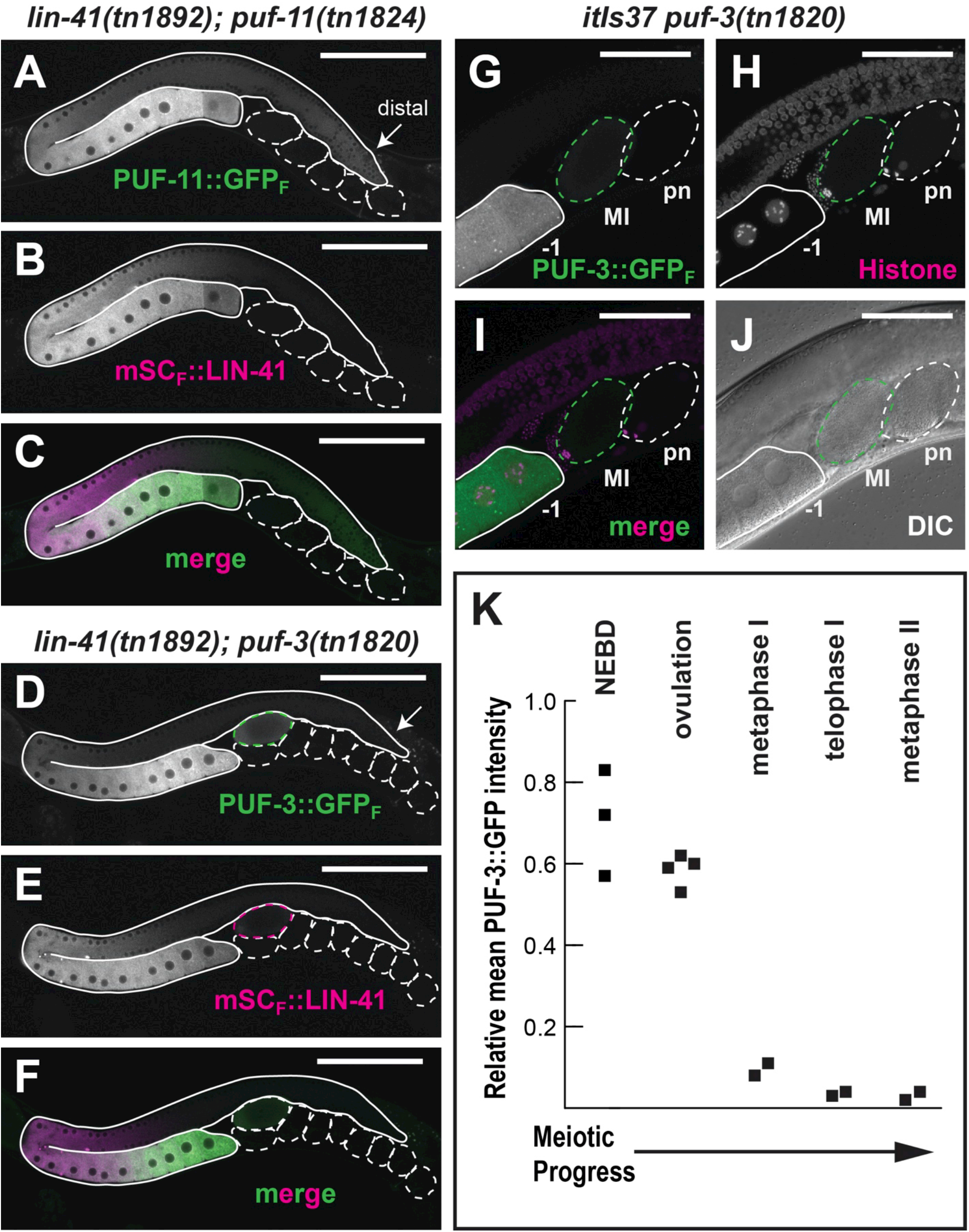
PUF-11::GFP and PUF-3::GFP are degraded during the OET. (A-F) PUF-11::GFP (A), PUF-3::GFP (D and G) and mSCARLET::LIN-41 (B and E) fluorescence in *C elegans* germlines (solid outlines) and early embryos (dashed outlines). The arrow in (A and D) indicates the distal mitotic region of each germline. Merged images of (A and B) and (D and E) are shown in (C) and (F), respectively. (G-J) Images of *it!s37[pie-1p::mcherry::histoneH2B::pie-1 3’UTR, unc-119(+)] puf-3(tn1820[puf-3::gfp::3xflag])* oocytes and embryos reveal that PUF-3::GFP (G) is strongly reduced in a meiosis I stage embryo (MI) relative to the −1 oocyte. Chromosomes are marked with mCHERRY:HISTONE (H), and the oocyte-derived chromosomes in this MI embryo appear to be aligning on the metaphase plate. PUF-3::GFP is at background levels in the adjacent post-meiotic 1-cell embryo (pn); this embryo has two polar bodies and maternal and paternal pronuclei at opposite ends of the embryo. Merged and DIC images of the same oocytes and embryos are shown in (I) and (J), respectively. The fluorescent images in this series (G-I) are 3D projections of a Z-stack. The DIC image (J) shows a single focal plane. Embryos with slightly higher levels of GFP and mSCARLET relative to older embryos are outlined in green and magenta, respectively (D, E, and G-J). Scale bars, 100 μm (A-F) and 50 μm (G-J). (K) PUF-3::GFP levels decrease after meiotic resumption. Mean GFP fluorescence was measured in images of *itls37[pie-lp::mcherry::histoneH2B::pie-l 3’UTR} puf-3(tn182O)* oocytes and embryos and background-corrected. Fluorescence intensity is expressed relative to the amount of GFP fluorescence in the oldest oocyte with an intact nucleus; this is the −2 oocyte if the −1 oocyte has initiated nuclear envelope breakdown (NEBD) or is starting to move into the spermatheca (early ovulation stage). Embryonic meiotic stage (e.g., metaphase I, telophase I and metaphase II) was classified based on the arrangement and organization of maternal chromosomes.

While LIN-41, PUF-3 and PUF-11 are strongly reduced soon after meiotic maturation and ovulation, each protein has a distinct pattern of expression in the adult germline (Figure 3, A-F). For example, the accumulation of PUF-3::GFP and PUF-11::GFP appears to peak at a later stage of oogenesis than mSCARLET::LIN-41 (Figure 3, C and F). In addition, PUF-11::GFP is distinctly more abundant than PUF-3::GFP in the distal germline (compare Figure 3, A and D; Figure S3D). Haupt *et al*. (2020) also observed higher levels of PUF-11 in the distal germline using alleles that express V5 epitope-tagged PUF-3 and PUF-11 proteins. Because *puf-3* and *puf-II* are redundant, to assess the function of the fusion proteins, we screened for and identified a recombinant that expresses both PUF-3::GFP and PUF-11::GFP using standard genetic methods (see Materials and Methods). As expected from the individual expression patterns, GFP expression was approximately two-fold higher in the oocytes of these animals [*puf-ll(tn1824) puf-3(tn1820)* genotype] than in the oocytes of either parent [*puf-ll(tn1824)* or *puf-3(tn1820)* alone], and the level of GFP expression in the distal germline was similar to the PUF-11::GFP-expressing strain (Figure S3, A-E). *puf-ll(tn1824) puf-3(tn1820)* animals are overtly wild-type; they are homozygous viable and fertile, with a brood size similar to the wild type (333 ± 40 larvae, n=15 animals at 20 °C). In addition, most of the embryos produced by *puf-ll(tn1824) puf-3(tn1820)* animals hatch (∼1% embryonic lethality). Since animals that are doubly homozygous for deletion alleles of *puf-ll* and *puf-3* [*puf-ll(q97l) puf-3(q966)* animals] exhibit maternal-effect embryonic lethality (Haupt *et al*. 2020), the GFP-tagged alleles of *puf-ll* and *puf-3* must retain some wild-type function. As a more stringent test, we crossed homozygous *puf-ll(tn1824) puf-3(tn1820)* males to balanced *puf-ll(q97l) puf-3(q966) I nTl[qIs51]* hermaphrodites to generate *puf-ll(tn1824) puf-3(tn1820) I puf-ll(q97l) puf-3(q966)* animals. Animals with this genotype produced many embryos that were viable (n=25 animals). We conclude that the GFP tag inserted at the C-terminus of PUF-3 and PUF-11 is largely compatible with the function of each protein.

### PUF-3 and PUF-11 are degraded in response to active CDK-1

LIN-41 is degraded in response to meiotic resumption and the activation of the cyclin-dependent kinase CDK-1 (Spike *et al*. 2014a, 2018). Likewise, we find that using RNA interference (RNAi) to strongly reduce CDK-1 activity causes PUF-3::GFP and PUF-11::GFP to persist in embryos (Figure 4, A and E; n>65 for each PUF::GFP with 100% penetrance). Furthermore, a strong RNAi-mediated reduction in the activity of the WEE-1.3 kinase, a negative regulator of CDK-1 (Burrows *et al*. 2006), causes oocytes to downregulate PUF-3::GFP and PUF-11::GFP prematurely (Figure 4, B and F). The latter phenotype was partially penetrant; it was observed in 32/46 of the PUF-3::GFP and 25/44 of the PUF-11::GFP-expressing *wee-l.3(RNAi)* gonads and mainly seen in *wee-l.3(RNAi)* animals with highly abnormal oocytes. GFP::LIN-41 is also degraded prematurely after *wee-l.3(RNAi)*, but this can be seen in gonads with relatively normal oocytes (Spike *et al*. 2014a), suggesting that LIN-41 might be more sensitive to CDK-1 activity. We tested this idea by treating animals that co-express mSCARLET::LIN-41 and PUF-11::GFP with *wee-l.3(RNAi)* and selectively examining gonads with large, well-organized oocytes. In these animals, mSCARLET::LIN-41 was down-regulated in younger (more distal) oocytes (Figure 4I) and, as a consequence, prematurely eliminated from oocytes more frequently than PUF-11::GFP (in 11/12 instead of 6/12 gonads, respectively). Collectively, these results indicate that active CDK-1 promotes the elimination of PUF-3 and PUF-11 and suggest that LIN-41 might be more sensitive to the dysregulatory effects of *wee-l.3(RNAi)*.

**Figure 4.**
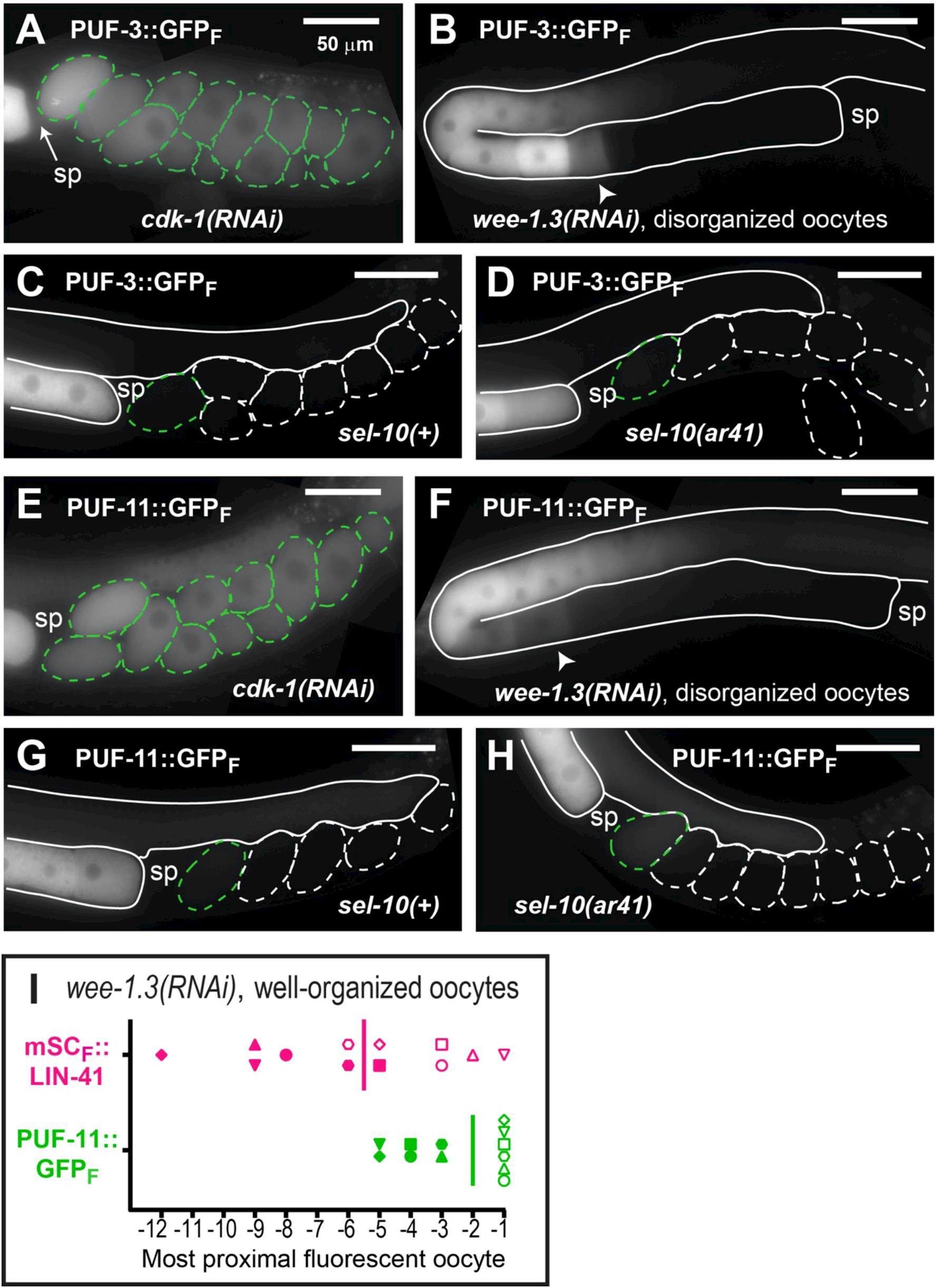
The elimination of PUF-3::GFP and PUF-11::GFP requires active CDK-1 but does not require SEL-10. PUF-3::GFP (A-D) and PUF-11::GFP (E-H) fluorescence in *C. elegans* germlines (solid outlines) and early embryos (dashed outlines) in *cdk-l(RNAi)* (A and E), *wee-l.3(RNAi)* (B and F), *lon-3(e2175)* (C and G) and *lon-3(e2175) sel-1O(ar4l)* (D and H) hermaphrodites. For each GFP fusion protein, the images of the *lon-3(e2175)* and *lon-3(e2175) sel-1O(ar4l)* animals were collected using the same settings (e.g., C and D). Embryos that appear GFP-positive are outlined in green. The slight reduction of PUF-3/11::GFP levels in more proximally located (i.e., older) arrested embryos in the uterus (A and E) might reflect CDK-1-independent turnover or it could result from residual CDK-1 activity following the RNAi treatment. The position of the spermatheca (sp) is indicated for reference. Arrowheads indicate the most proximal oocyte in each *wee-l.3(RNAi)* gonad that is strongly GFP-positive. (I) This graph illustrates the relative position of the most proximal mSCARLET or GFP-positive oocyte in 12 different *lin-41(tn1892[mScarlet::3xflag::lin-41]); puf-ll(tn1824[puf-ll::gfp::3xflag]); wee-l.3(RNAi)* animals. *wee-l.3(RNAi)* animals with well-formed and organized oocytes were scored. Oocyte position was measured relative to other oocytes and the spermatheca (position 0). Each animal is represented by a different symbol (e.g., filled diamond or open circle). The selection of a filled or open symbol is arbitrary and made to depict the 12 animals scored with six symbol shapes. The color of the symbol represents the fusion protein that was scored: mSCARLET::LIN-41 in magenta and PUF-3::GFP in green. The vertical bar indicates the median. Scale bars, 50 μm (A-H).

### SCF_SEL-10_ and APC ubiquitin ligases do not target PUF-3 or PUF-11 for degradation

PUF-3 and PUF-11 are components of LIN-41 RNPs and appear to be eliminated in response to the same signal and at a similar time as LIN-41. We therefore asked whether they are targeted for elimination by the same SCF E3 ubiquitin ligase. LIN-41 persists in early embryos after the RNAi-mediated knockdown of the highly similar Skp-related genes *skr-l* and *skr-2,* the Cullin *cul-l*, the F-box substrate recognition subunit *sel-1O* and in the *sel-1O(lf)* mutants *sel-1O(ok1632)* and *sel-1O(ar41)* (Spike *et al*. 2018). However PUF-3::GFP and PUF-11::GFP are eliminated normally from the embryos of *lon-3(e2175) sel-1O(ar41)* mutant animals (Figure 4, C, D, G, and H) and after *skr-1/2(RNAi)* (Figure S4, C and I, n>40 animals for each PUF::GFP fusion protein) and *cul-1(RNAi)* (Figure S4, B and H, n2:20 animals for each PUF::GFP fusion protein). A distinct, SEL-10-independent mechanism must therefore promote the elimination of PUF-3 and PUF-11 from early embryos. Furthermore, this indicates that the degradation of PUF-3 and PUF-11 during the OET is not a consequence of the SEL-10-dependent degradation of LIN-41, consistent with our data suggesting that LIN-41 is more sensitive to *wee-1.3(RNAi)* and the idea that multiple mechanisms remodel and take apart oocyte RNPs during the OET.

An interesting feature of LIN-41 degradation is that it occurs even when the metaphase- to-anaphase transition of the first meiotic division is blocked by disrupting the function of the anaphase-promoting complex (APC) (Spike *et al*. 2014a). This feature distinguishes LIN-41 from proteins such as IFY-1/Securin (Kitagawa *et al*. 2002; Wang *et al*. 2013) and CYB-1/Cyclin B1 (Liu *et al*. 2004), which are degraded slightly later than LIN-41 in an APC-dependent fashion. Therefore, we tested whether the metaphase-to-anaphase transition regulates the elimination of PUF-3::GFP by impairing the function of two key members of the APC, *mat-1*/Apc3 and *emb-3O*/Apc4 (Furuta *et al*. 2000; Shakes *et al*. 2003). *mat-1(RNAi)* caused fully penetrant 1-cell embryonic arrest, but did not delay or prevent the elimination of PUF-3::GFP from embryos (Figure S4D; n=34 animals). Likewise, PUF-3::GFP disappears normally from the embryos of *emb-3O(tn377*ts) mutants shifted to 25 °C at the 14 stage (Figure S4F; n > 35); these embryos arrest in metaphase during the first meiotic division (Furuta *et al*. 2000). PUF-11::GFP degradation was also unaffected by *mat-1(RNAi)* (Figure S4J; n=9). We conclude that, as for LIN-41, the elimination of either PUF-3 or PUF-11 from early embryos does not require the activity of the APC. However, unlike LIN-41, the elimination of PUF-3::GFP and PUF-11::GFP from embryos is *sel-10* independent.

### ETC-11Hu151UBE3C promotes efficient protein clearance during the OET

Since PUF-3/11 are degraded in a SEL-10-independent manner, and because regulated protein degradation often requires ubiquitination (reviewed by Dikic *et al*. 2017), we tested whether a different E3 ubiquitin ligase might promote the elimination of PUF-3::GFP from embryos. Many potential ubiquitin ligases have been identified in *C. elegans*, including (1) multi-subunit E3s (SCF-type, APC), (2) HECT-domain E3s, (3) U-box E3s, and (4) more than 100 RING-finger proteins that might function as monomeric E3s (Kipreos *et al*. 2005). We performed a small-scale RNAi screen that targeted the first three classes of ubiquitin ligases and identified the HECT-domain ubiquitin ligase ETC-1/Hu15/UBE3C as a promising candidate. When animals expressing both mSCARLET::LIN-41 and PUF-3::GFP [*lin-41(tn1892); puf-3(tn1820)* genotype] were exposed to *etc-1(RNAi)*, there was no discernable change in mSCARLET::LIN-41 degradation, yet elevated levels of GFP were observed in multiple young embryos (n=24/24 animals). This suggested that *etc-1* knock down specifically interferes with the elimination of oocyte-expressed PUF-3::GFP but not mSCARLET::LIN-41. We confirmed these RNAi-based observations with *etc-1(gk5182)*, a predicted loss-of-function allele, which introduces a premature stop codon, generated by the *C. elegans* Reverse Genetics Core Facility at the University of British Columbia (The *C. elegans* Deletion Mutant Consortium 2012). *etc-1(gk5182); puf-3(tn1820)* and *etc-1(gk5182); puf-ll(tn1824)* animals exhibit GFP fluorescence in multiple embryos compared to *etc-1(+)* controls (Figure 5, A-D), while expanded mSCARLET is not observed in *lin-41(tn1892); etc-1(gk5182)* animals (Figure 5, E and F). However, detailed analysis presented in subsequent sections suggests that ETC-1 unlikely functions as an E3 ubiquitin ligase that targets specific proteins for degradation during the OET but instead functions as a factor that increases the processivity of the proteasome.

**Figure 5.**
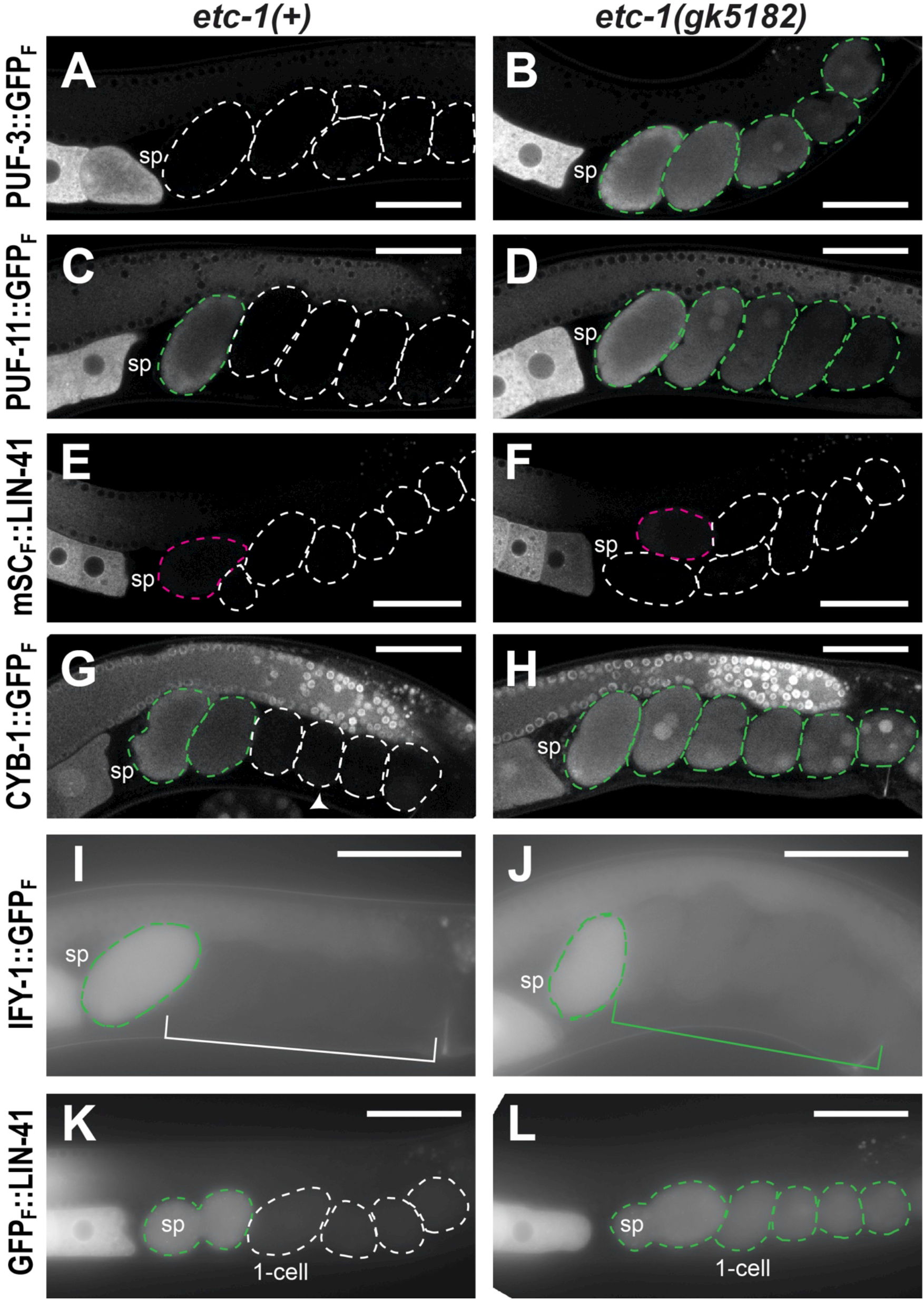
*etc-1(gk5182)* perturbs the degradation of GFP-fusion proteins during the OET. PUF-3::GFP (A and B), PUF-11::GFP (C and D), mSCARLET::LIN-41 (E and F), CYB-1::GFP (G and H), IFY-1::GFP (I and J) and GFP_F_::LIN-41 (K and L) fluorescence in young *etc-1(+)* (A, C, E, G, I, and K) and *etc-1(gk5182)* (B, D, F, H, J, and L) embryos. In each image, the position of the spermatheca (sp) is indicated for reference. Young embryos in the uterus are indicated with dashed outlines that highlight individual embryos (A-H, K, and L) or a bracket that indicates a region containing multiple post-meiotic embryos (I and J). GFP-positive embryos are indicated in green (A-D and G-L) while mSCARLET-positive embryos are indicated in magenta (E and F). For each fusion protein, these embryos contain detectably more fluorescence than the *etc-1(+)* embryos indicated in white. CYB-1::GFP re-accumulates to relatively high levels in older *etc-1(+)* embryos (G, arrowhead; this embryo is no longer in the uterus) and therefore has a distinctly different expression pattern than the other OET-degraded fusion proteins. Scale bars, 50 μm.

ETC-1 was proposed to function as an E3 ubiquitin ligase that regulates the level of cytoplasmic IFY-1/Securin and CYB-1/cyclin B1 in post-meiosis I embryos in *C. elegans* (Wang *et al*. 2013). In that study, RNA interference was used to reduce *etc-1* function [*etc-1(RNAi)*] and CYB-1 degradation was monitored using a transgene that uses heterologous regulatory elements to drive the expression of a GFP::CYB-1 fusion protein in the *C. elegans* germline (*ekIs2[pie-lp::gfp::cyb-l]*). We used a new viable and fertile allele of *cyb-l* that expresses a GFP-tagged CYB-1 fusion protein (Figure S1C; CYB-1::GFP_F_) to examine CYB-1::GFP expression in wild-type and in *etc-1(gk5182)* embryos (Figure 5, G and H). Since CYB-1 is essential for viability (van der Voet *et al*. 2009), the CYB-1::GFP fusion protein made by *cyb-l(tn1806[cyb-l::gfp::3xflag])* homozygotes must be substantially functional *in vivo*. In otherwise wild-type animals, CYB-1::GFP is reduced in young mitotic embryos relative to meiotic embryos and oocytes (Figure 5G). However, in *etc-1(gk5182); cyb-l(tn1806)* embryos GFP levels continue to remain high after meiosis (Figure 5H). We also confirmed that *etc-1(gk5182)* affects the degradation of IFY-1::GFP (IFY-1::GFP_F_) using the fosmid-based transgene *ddIs128[ify-l::gfp::3xflag]* (Sarov *et al*. 2012). *etc-1(gk5182)* appears to have a relatively weak effect on IFY-1::GFP (Figure 5, I and J) compared to PUF-3::GFP, PUF-11::GFP and CYB-1::GFP (Figure 5, A-D and G-H). The amount of GFP fluorescence in post-meiotic *etc-1(gk5182); ddIs128* embryos is only slightly elevated relative to *etc-1(+)* control embryos (Figure 5, I and J). We quantified this modest increase by comparing the IFY-1::GFP fluorescence levels in +2 embryos from *etc-I(gk5I82)* animals and *etc-I(+)* controls and found it to be increased 1.4-fold in *etc-I* mutants (P<1×10^-4^, *t*-test; n=12 for each genotype).

### ETC-1 is broadly expressed and present in oocytes and early embryos

ETC-1 and its orthologs, Hu15 (yeast) and UBE3C (humans) have two well-defined structural domains: an IQ domain close to the N-terminus and a HECT domain at the extreme C-terminus (Figure 6A; Wang *et al*. 2013; Marin 2018). The HECT domain of UBE3C and other HECT-domain E3 ubiquitin ligases has a well-defined function; it is essential for mediating the acceptance and transfer of ubiquitin from an E2-conjugating enzyme to a degradation substrate via a cysteine residue at its catalytic site (You and Pickart 2001; Singh and Sivaraman 2020; reviewed in Lorenz 2018). The IQ domain-containing N-terminus of UBE3C mediates an interaction with the proteasome that is also important for function, but this portion of UBE3C is less well-conserved than the HECT domain and its function is less well-defined (You and Pickart 2001; You *et al*. 2003; Wang *et al*. 2013). Given this structural and functional information, we decided to tag the extreme N-terminus with GFP to examine where ETC-1 is expressed and localized in *C. elegans*.

**Figure 6.**
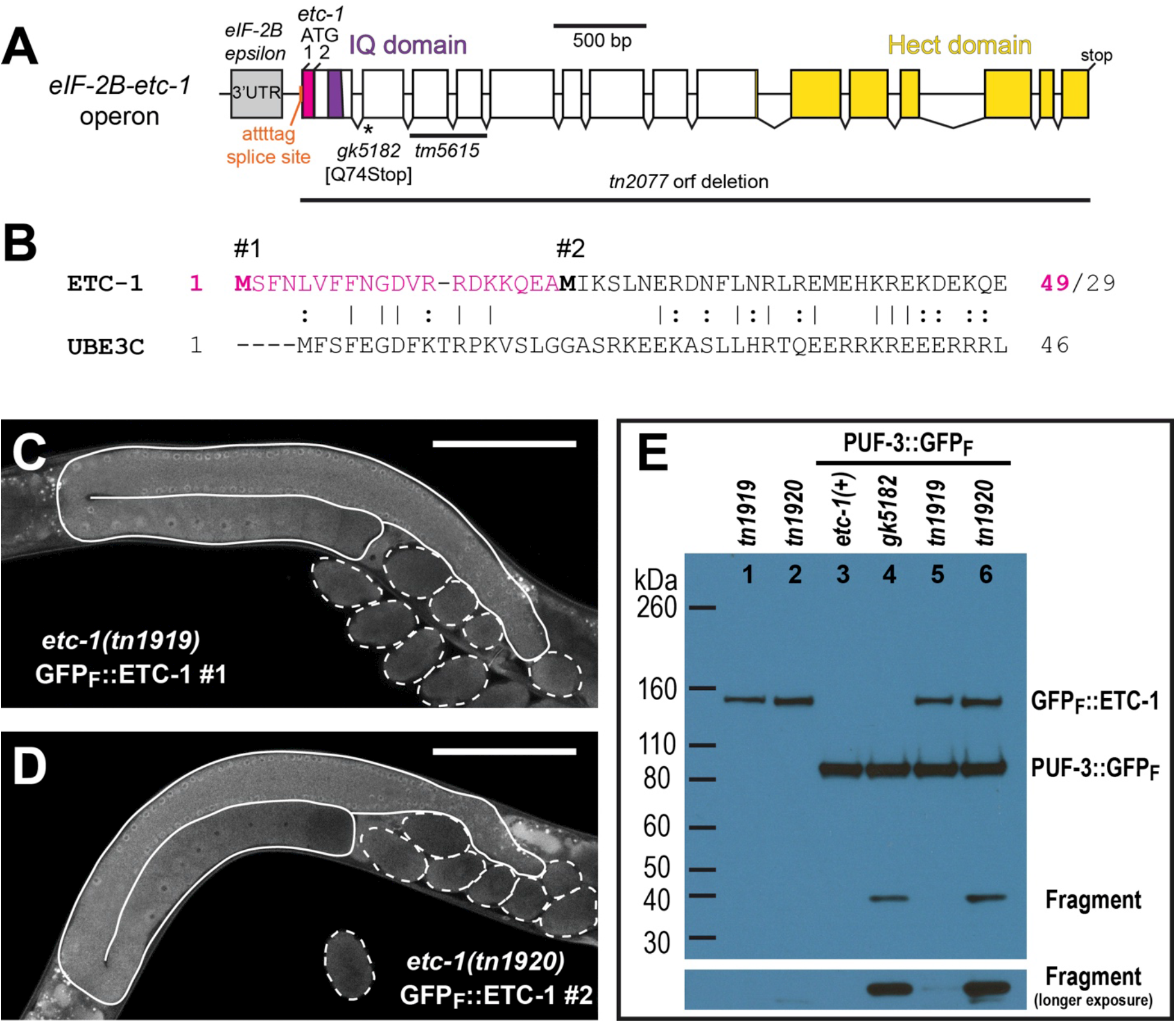
ETC-1 is expressed in the *C. elegans* germline. (A) *etc-1* alleles and the exon-intron structure of *etc-1*, the downstream gene in operon CEOP2380. Exons are shown as open boxes; the region of exon 1 colored in pink would encode 20 additional amino acids if translation is initiated at ATG #1, the start codon immediately adjacent to the annotated S11/S12 trans-splice site (orange), instead of ATG #2. Other colors indicate the 3’ UTR of the upstream gene eIF-2B (gray) and the regions that encode the conserved IQ (purple) and HECT (yellow) domains of ETC-1. (B) The N-terminus of ETC-1 aligned with the N-terminus of the human ortholog UBE3C; the amino acids predicted to be added to ETC-1 are in pink. This is the N-terminal portion of a full-length protein alignment generated by EMBOSS Needle with default alignment parameters. ETC-1 and UBE3C were 26% identical or 46% similar overall with 16% of the alignment gapped. (C and D) GFP::ETC-1 fluorescence in *C. elegans* germlines (solid outlines) and early embryos (dashed outlines). GFP is N-terminal to either the first (C, *etc-1(tn1919[gfp::3xflag::etc-1 #1})*) or second (D, *etc-1(tn1920[gfp::3xflag::etc-1#2})*) methionine of ETC-1. Scale bars, 100 μm. (E) A western blot of GFP::ETC-1 (∼150 kDa, see main text) and PUF-3::GFP-expressing strains with an anti-FLAG antibody that recognizes both fusion proteins. GFP::FLAG fragments derived from PUF-3::GFP strongly accumulate in *etc-1(gk5182)* and *etc-1(tn1920[gfp::3xflag::etc-1#2})* hermaphrodites.

When we examined the sequences at the 5’ end of the *etc-I* gene (Figure 6A) it became apparent that ETC-1 translation might begin 60 bp upstream of the annotated start codon (ATG #2 in Figure 6A) (http://www.wormbase.org, release WS278). *etc-I* is the downstream gene in a two gene operon and contains an annotated and sequence-verified trans-splice acceptor site immediately adjacent to a possible start codon (ATG #1 in Figure 6A). In *C. elegans*, trans-splicing is used to physically separate the co-transcribed messenger RNAs of an operon (Spieth *et al*. 1993). We predicted that ETC-1 translation should be initiated at ATG #1 for the following reasons: (1) start codons are commonly found within 10 bp of a trans-splice acceptor site (Allen *et al*. 2011), and (2) after trans-splicing, ATG #1 would be the first start codon in the *etc-1* mRNA. If correct, 20 additional amino acids would be appended to the N-terminus of the predicted ETC-1 protein. Seven of these amino acids are similar or identical to equivalent residues at the N-terminus of UBE3C (Figure 6B). We created two different GFP-tagged alleles of *etc-1*: (1) *etc-1(tn1919[gfp::3xflag::etc-1#l])*, which places GFP (and another start codon) immediately upstream of the start codon adjacent to the trans-splice acceptor site (ATG #1), and (2) *etc-1(tn1920[gfp::3xflag::etc-1#2])*, which places an identical tag immediately upstream of the annotated start codon (ATG #2) (Figure S5A). Both *etc-1(tn1919)* and *etc-1(tn1920)* express GFP (Figure 6, C and D, and Figure S5, B and C) and make large fusion proteins that approximate the predicted sizes of the GFP::ETC-1 fusion proteins (∼150 kDa, see below) (Figure 6E, lanes 1 and 2). We conclude that a start codon immediately adjacent to the *etc-1* 5’ trans-splice site is capable of initiating translation *in vivo*.

*etc-1(tn1919)* and *etc-1(tn1920)* express GFP::ETC-1 (labeled GFP_F_::ETC-1) in the oogenic germline of hermaphrodites, where it is both cytoplasmic and nucleoplasmic (Figure 6, C and D). We observed no significant difference in the amount of cytoplasmic GFP::ETC-1 in the germlines of *etc-1(tn1919)* (n=11) and *etc-1(tn1920)* (n=8) adult hermaphrodites (*P*>0.1 using a Student’s *t*-test). In addition, both alleles express GFP::ETC-1 in embryos (Figure 6, C and D) and in a variety of somatic tissues, including the intestine, the spermatheca, and unidentified cells near the pharynx (arrow in Figure S5, B and C). Interestingly, *etc-1(tn1920)* adult hermaphrodites (n=3) express 2.4-fold more GFP::ETC-1 in the unidentified cells near the pharynx than *etc-1(tn1919)* adult hermaphrodites (n=7) (*P*<1×10^-6^ using a Student’s *t*-test). This result suggests there could be differential start codon usage in some cell types. In *etc-1(tn1920)* animals, the start codon at the beginning of the GFP coding sequence is 60 bp downstream of ATG #1 and in the same relative position as the annotated start codon (ATG #2) in the endogenous *etc-1* gene (Figure S5A). We hypothesized that *etc-1(tn1920)* animals might be able to initiate GFP::ETC-1 translation both at the start codon used by *etc-1(tn1919)* animals (150.5 kDa GFP::ETC-1) and at the start codon located at the beginning of the GFP coding sequence (148 kDa GFP::ETC-1). Consistent with this idea, some of the GFP::ETC-1 made by *etc-1(tn1920)* animals appears to migrate slightly faster on western blots (Figure 6E, lane 2). These observations suggest that ETC-1 translation can initiate at ATG #1 in most tissues, including the germline, while leaving the possibility that ETC-1 translation also initiates at ATG #2 in some somatic cells. However, it is also possible that the slightly different GFP expression patterns seen in *etc-1(tn1919)* and *etc-1(tn1920)* are caused by a difference in gene structure that we have not explicitly considered or the fact that these alleles are differently functional. As we will describe, *etc-1(tn1919)* (ATG #1) appears to be largely functional during the OET, while *etc-1(tn1920)* (ATG #2) resembles the strong loss-of-function *etc-1(gk5182)* allele.

### ETC-1 is non-essential

*etc-1(RNAi)* and *etc-1(tm5615)* deletion animals are viable and fertile; they do not have an overt phenotype in an otherwise wild-type background (Wang *et al*. 2013). Whereas *etc-1(tm5615)* is predicted to be an in-frame deletion (Figure 6A), *etc-1(gk5182)* is a premature stop codon in the second exon of the *etc-1* gene (Figure 6A). This suggested to us that *etc-1(gk5182)* should be a strong loss-of-function allele. We therefore examined these animals for a variety of phenotypes, including fertility defects, but failed to find any obvious problems. For example, the brood size of *etc-1(gk5182)* hermaphrodites was no different from that of the wild type at 20 °C [285.0 ± 25.2 (n=23) for *etc-1(gk5182)* vs. 292.8 ± 20.7 (n=20) in the wild type; *P*= 0.2784, *t* test]. *etc-1(gk5182)* hermaphrodites also exhibited only low levels of embryonic lethality [0.3 ± 0.3 % embryonic lethality (n=6575) for *etc-1(gk5182)* vs. 0.2 ± 0.3 (n=5853) in the wild type]. We tested whether the strong loss-of-function mutation *sel-1O(ar41)* (Figure S1A) would provide a sensitized background to reveal defects because SEL-10 and ETC-1 both help degrade proteins during the first meiotic division. However, *etc-1(gk5182)*; *lon-3(e2175)* and *etc-1(gk5182)*; *lon-3(e2175) sel-1O(ar41)* hermaphrodites are essentially indistinguishable and produce large numbers of progeny [286 ± 26 larvae (n=14) and 283 ± 29 larvae, (n=15), respectively, at 20 °C]. These brood sizes are similar to the brood sizes of *lon-3(e2175)* and *lon-3(e2175) sel-1O(ar41)* hermaphrodites at 20 °C (Spike *et al* 2018). We examined whether *etc-1(gk5182)* animals are unusually sensitive to elevated temperatures because Hu15, the ETC-1 ortholog of budding yeast, facilitates the degradation of protein aggregates that form during heat stress (Fang *et al*. 2011). However, *etc-1(gk5182)* animals are homozygous viable and fertile at 25 °C and can be maintained at this temperature for multiple generations. When wild-type animals are heat-stressed for 3 hours at 31 °C, partially penetrant embryonic lethality occurs in the 24-hour period immediately after the stress is applied (Huelgas-Morales *et al* 2016). Using this heat stress protocol, similar percentages of embryos produced within the relevant 24-hour period died when we compared animals with the *etc-1(gk5182)* genotype (21% lethality, n=565 embryos, n=12 animals) to the wild type (26% lethality, n=463 embryos, n=11 animals). Wild-type 14-stage larvae shifted from 20 °C to 37 °C for 1.5 hours exhibit partially penetrant larval arrest and lethality (Zevian and Yanowitz 2014). Using this protocol, similar percentages of wild-type (54%, n=268) and *etc-I(gk5I82)* mutant animals (48%, n=300) died after heat shock.

To unambiguously determine whether ETC-1 is important for fertility or viability, we used CRISPR-Cas9 genome editing to generate the *etc-I(tn2077)* null allele, which deletes the entire *etc-I* open reading frame (Figure 6A). *etc-I(tn2077)* mutants are viable and fertile at all temperatures tested (15, 20, and 25°C), confirming that *etc-I* is a non-essential gene in *C. elegan*s under standard laboratory conditions. Like *etc-I(gk5I82)* hermaphrodites, *etc-I(tn2077)* hermaphrodites exhibit negligible embryonic lethality [0.3 ± 0.6 % embryonic lethality (n=4304)]. However, *etc-I(tn2077)* hermaphrodites produce significantly lower brood sizes than *etc-I(gk5I82)* hermaphrodites [178.8 ± 55.6 (n=24) for *etc-I(tn2077)* vs. 285.0 ± 25.2 (n=23) for *etc-I(gk5I82)*; P < 0.0001, *t* test]. We also observed that *etc-I(tn2077)* hermaphrodites were sluggish and appeared to exhibit a reduced life span (D.G., unpublished results). The reduced brood size of *etc-I(tn2077)* hermaphrodites might be due to the production of less sperm, which is ordinarily limiting for fertility. Whether this speculation proves correct and reflects the role of ubiquitin-mediated protein degradation in germline sex determination (Starostina *et al*. 2007), will require further study. Although the differences between *etc-I(gk5I82)* and the large *etc-I(tn2077)* deletion could in principle be due to impacts on neighboring genes, these phenotypic differences are consistent with the idea that *etc-I(gk5I82)* possibly retains residual ETC-1 function.

### etc-I alleles exhibit a genetic interaction with the anaphase promoting complex

Wang *et al*. (2013) reported that *etc-I(RNAi)* enhances the phenotype of *mat-I(axI6I*ts*); him-8(eI489)* animals at the permissive temperature, and causes them to produce one-cell arrested embryos. This enhancement supported the idea that ETC-1 and the anaphase promoting complex (APC) share redundant functions during the OET, including the degradation of the key cell cycle regulators IFY-1/securin and CYB-1/Cyclin B. We initially revisited this genetic interaction using *etc-I(gk5I82)* and two different temperature-sensitive alleles that attenuate APC function in *C. elegans*: *emb-30(tn377*ts) and *mat-I(axI6I*ts) (Furuta *et al*. 2000, Shakes *et al*. 2003). Contrary to expectation, we found that *etc-I(gk5I82); emb-30(tn377*ts) and *mat-I(axI6I*ts*); etc-I(gk5I82)* double mutants are each homozygous viable and fertile at 15 °C. To extend this result, we examined the number of larvae produced by *mat-I(axI6I*ts*); etc-I(gk5I82)* and *mat-I(axI6I*ts) parents. These animals were initially derived from *mat-I*-balanced strains (Table S1). For each strain, we counted the number of larvae produced by parents that were third-generation *mat-I(axI6I*ts) homozygotes raised at 15 °C. *mat-I(axI6I*ts*); etc-I(gk5I82)* animals produced fewer progeny (100 ± 35 larvae, n=15 animals) than *mat-I(axI6I*ts) animals (165 ± 46 larvae, n=17 animals) and this difference is statistically significant (*P*=0.0001). However, we did not observe that the *mat-I(axI6I*ts*); etc-I(gk5I82)* double mutants produced one-cell arrested embryos. Thus, the double mutant phenotype is not comparable to the embryonic arrest phenotype of *mat-I(axI6I*ts) (Figure S6B) or *mat-I(axI6I*ts); *him-8(eI489)* animals at 25 °C (Shakes *et al*. 2003).

Since our phenotypic analyses suggested that *etc-I(gk5I82)* might retain some residual function, we analyzed whether *etc-I(tn2077)* exhibits a stronger genetic interaction with *mat-I(axI6I*ts). We observed that *mat-I(axI6I*ts*); etc-I(tn2077)* hermaphrodites are viable and fertile at 15°C and produce embryos that can undergo multiple rounds of cell division (Figure S6C). Some of these embryos hatch and grow into larvae. However, *mat-l(ax16l*ts*); etc-1(tn2077)* hermaphrodites produce many fewer larvae than *mat-l(ax16l*ts*); etc-1(gk5182)* hermaphrodites at 15 °C. First-generation homozygotes derived from *mat-l(ax16l*ts)/*tmC18*; *etc-1(tn2077)* parents produced, on average, 28 ± 19 larvae (n=18). Second-generation homozygotes were also fertile and produced a similarly small number of larvae [24 ± 14 larvae (n=36)]. However, dead embryos were also observed among the progeny of *mat-l(ax16l*ts*); etc-1(tn2077)* hermaphrodites, and dead embryos were typically more common than hatched larvae when both of these were counted (n=12 of 18 second-generation egg lays). Although we did not observe the one-cell arrest phenotype or the oocyte defects that were previously described (Figure S6), these observations nonetheless confirm that *mat-l(ax16l*ts) embryonic lethality is enhanced by *etc-1(tn2077)*, albeit at a later stage of embryogenesis that may result from a defect in mitotic cell division.

### GFP-fusion proteins are partially proteolyzed in etc-I(gk5I82) animals

The PUF-3::GFP, PUF-11::GFP, CYB-1::GFP and IFY-1::GFP fusion proteins each contain three FLAG epitope tags in addition to the fluorescent protein tag (Figure S1C, Sarov *et al*. 2012). We used an antibody that recognizes the FLAG tag to examine each fusion protein by western blot in wild-type and *etc-1(gk5182)* hermaphrodites. Full-length PUF-3::GFP, PUF-11::GFP and CYB-1::GFP fusion proteins migrate as expected in lysates made from both wild-type and *etc-1(gk5182)* hermaphrodites (Figure 7A, lanes 1-6), while the IFY-1::GFP fusion protein migrates slightly slower than expected (Figure 7C, lanes 1 and 2). However, in each *etc-1(gk5182)* lysate, a second band is also recognized by the anti-FLAG antibody (Figure 7A, lanes 2, 4, and 6, and Figure 7C, lane 2). This *etc-1(gk5182)*-specific band only accumulates when a GFP fusion protein is present (Figure S7A, compare lanes 2-7 with lanes 1 and 8) and can be recognized by a GFP-specific antibody (Figure S7B, lanes 2, 4, and 6). Depending on the GFP fusion protein, it migrates slightly faster (Figure 7A, lanes 2, 4, and 6) or slower (Figure 7C, lane 2) than a 40 kDa molecular weight marker. But in each case, the *etc-1(gk5182)*-specific band is predicted, based on size, to encompass the entire 31-32 kDa GFP::FLAG tag (File S2). We estimate that these GFP::FLAG fragments account for 10-20% of the total signal in each *etc-1(gk5182); puf-3(tn1820[puf-3::gfp::3xflag])* lane, 4-11% of the total signal in each *etc-1(gk5182); puf-11(tn1824[puf-11::gfp::3xflag)* lane and 4-10% of the total signal in each *etc-1(gk5182); cyb-1(tn1806[cyb-1::gfp::3xflag])* lane (Figure 7A, lanes 2, 4, and 6). GFP::FLAG fragments appear to be less abundant in *etc-1(gk5182); ddls128[ify-1::gfp::3xflag]* animals relative to the full length IFY-1::GFP fusion protein (Figure 7C), but we could not make an accurate assessment of this difference because the full-length fusion protein and fragment were not in the linear range of detection at the same time. These observations suggest that the GFP fluorescence observed in *etc-1(gk5182)* embryos (Figure 5, A-D and G-J) may derive from GFP::FLAG fragments rather than full-length PUF-3::GFP, PUF-11::GFP, CYB-1::GFP, or IFY-1::GFP fusion proteins.

**Figure 7.**
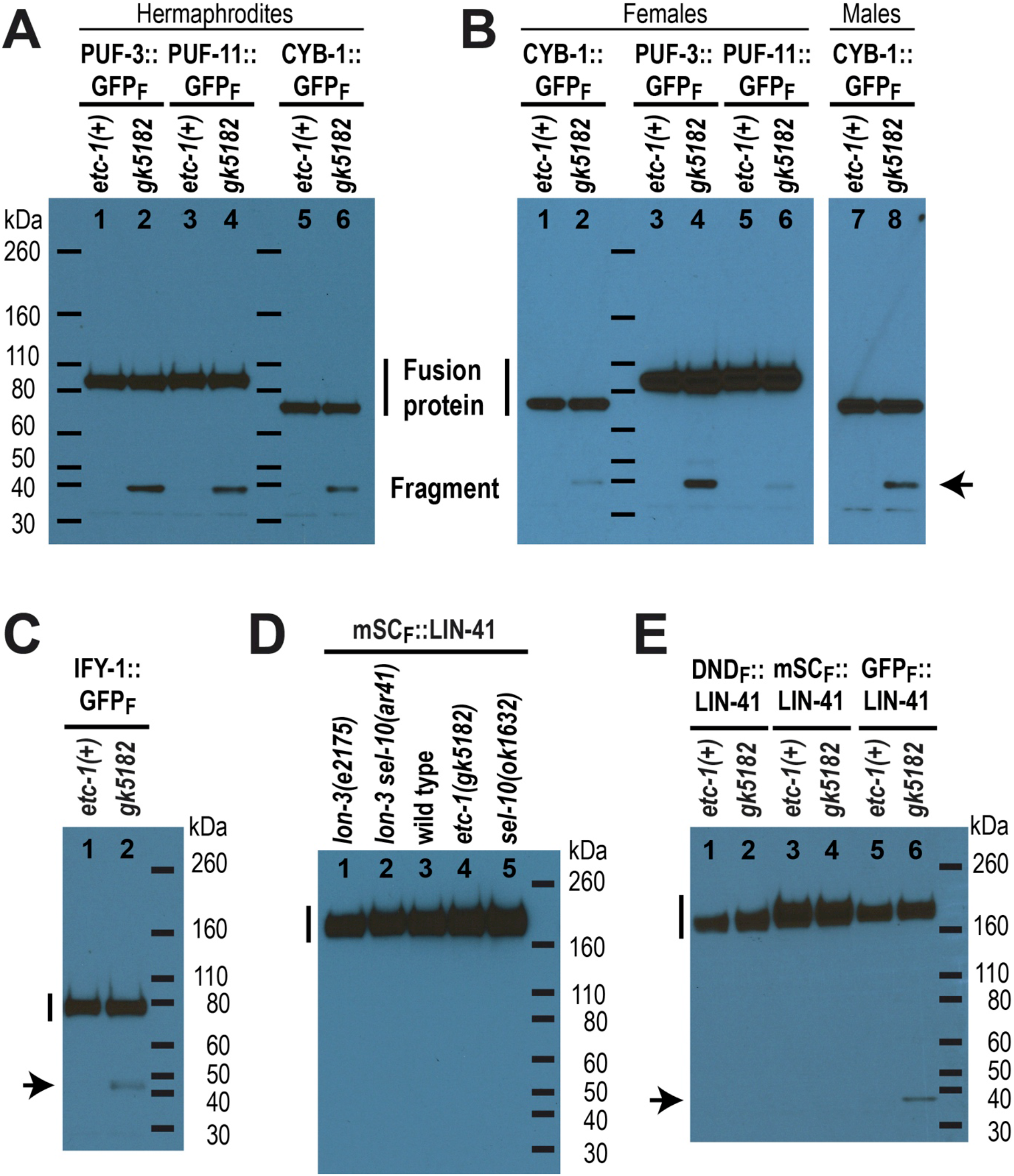
GFP-tagged fusion proteins are partially proteolyzed in *etc-1(gk5182)* animals. (A-E) Western blots with a FLAG-specific primary antibody. Three FLAG tags are located at the extreme C-terminus of PUF-3::GFP, PUF-11::GFP, CYB-1::GFP and IFY-1::GFP and in-between LIN-41 and each fluorescent protein (mSCARLET, DENDRA, or GFP). The predicted molecular mass of each full-length fusion protein (vertical line) is ∼89 kDa for each PUF::GFP fusion protein (A and B), 73 kDa for CYB-1::GFP (A and B), 61 kDa for IFY-1::GFP (C), and 154-156 kDa for each LIN-41 fusion protein (D and E), and. The same set of molecular weight markers (horizontal lines) were used in (A) and (B). A partially proteolyzed GFP::FLAG-containing fragment >32 kDa (arrow) accumulates in the *etc-1(gk5182)* animals that express a GFP fusion protein (A, B, C, and E). The protein lysates were made from adult hermaphrodites (A, C, D, and E), adult females (B) or adult males (B), as indicated.

Our observations of GFP fluorescence in living *etc-1(lf)* animals suggest that *etc-1(gk5182)* perturbs protein degradation during the OET (Figure 5). However, if the GFP::FLAG fragments generated from each fusion protein accumulate prior to this time (during oogenesis, for example), they might simply become easier to visualize when the full-length fusion protein is degraded during the OET. We examined whether GFP::FLAG fragments are generated from the PUF-3::GFP, PUF-11::GFP and CYB-1::GFP fusion proteins in unmated *etc-1(gk5182); fog-2(oz40)* females to address this possibility. Development of the oogenic germline is normal in *fog-2* mutant females, but due to the absence of the major sperm protein signal from sperm (McCarter *et al*. 1999; Miller *et al*. 2001), the rate of oocyte meiotic maturation is substantially decreased, and fertilization and embryo formation are prevented. We found that GFP::FLAG fragments are generated from each fusion protein in *etc-1(gk5182); fog-2* females (Figure 7B, lanes 2, 4, and 6; see Figure S7D for a longer exposure). Relative to the amount of each full-length fusion protein, however, GFP::FLAG fragments appear to be considerably less abundant in unmated *etc-1(gk5182); fog-2* females than in *etc-1(gk5182)* hermaphrodites (compare Figure 7, A and B) or mated *etc-1(gk5182); fog-2* females (compare Figure 7B and Figure S7, C and D). This was most noticeable for the GFP::FLAG fragments derived from CYB-1::GFP and PUF-11::GFP. These observations suggest that many of the GFP::FLAG fragments produced by *etc-1(gk5182)* mutant hermaphrodites and mated females are generated when each GFP fusion protein is degraded during the OET.

GFP::FLAG fragments also accumulate in *etc-1(gk5182); cyb-l(tn1806); fog-2(oz40)* adult males (Figure 7B, lane 8). Endogenous CYB-1 is abundant in the male germline and eliminated toward the end of meiosis (Yoon *et al*. 2017). CYB-1::GFP exhibits a similar pattern of accumulation in *fog-2* males (Figure S8, A and B). In *etc-1(gk5182); fog-2* males, however, a GFP signal perdures past the stage where CYB-1::GFP is normally eliminated (Figure S8, C and D). Much of this signal localizes to residual bodies (Figure S8, E and F). Residual bodies form at the end of the second meiotic division and receive cellular materials that are not required during subsequent stages of spermatogenesis (Ward *et al*. 1981; reviewed in Chu and Shakes 2013). It seems likely that, as in *etc-1(gk5182)* embryos, the GFP fluorescence seen in *etc-1(gk5182)* residual bodies derives from GFP::FLAG fragments that are formed when CYB-1::GFP is degraded during male meiosis.

### ETC-I may promote proteasome processivity

Hu15 and UBE3C, the yeast and human orthologs of ETC-1, are proteasome-associated ubiquitin ligases that help the proteasome degrade difficult substrates (Crosas *et al*. 2006; Fang *et al*. 2011; Kuo and Goldberg 2017; Gottlieb *et al*. 2019). In this context, Hu15 and UBE3C are thought to be important for the processive degradation of proteins that are stalled on the proteasome. This hypothesis was based, in part, on experiments with GFP fusion proteins. In the absence of Hu15, or after UBE3C knockdown, the proteasomal degradation of substrates consisting of an unstable domain fused with the much more stable GFP domain is incomplete. This results in the accumulation of stable GFP-containing fragments of the original fusion protein (Aviram and Kornitzer 2010; Martinez-Noel *et al*. 2012; Chu *et al*. 2013). The experiments described in the previous section strongly suggest that ETC-1 plays a similar role in *C. elegans*. We added to these observations by examining additional GFP and non-GFP-tagged fusion proteins in *etc-1(lf)* hermaphrodites, as described below.

#### GFP::ETC-I fusion proteins

Neither GFP::ETC-1 fusion protein is partially processed to a substantial extent (Figure 6E, lanes 1 and 2). To determine whether either GFP-tagged allele is functional *in vivo*, we examined whether PUF-3::GFP is partially processed in *etc-1(tn1919)* and *etc-1(tn1920)* hermaphrodites. GFP::FLAG fragments accumulate in PUF-3::GFP-expressing *etc-1(tn1920)* animals (Figure 6E, lane 6) and are similar in abundance to the GFP::FLAG fragments seen in *etc-1(gk5182)* animals (Figure 6E, lane 4). *etc-1(tn1920)* therefore appears to be a strong loss-of-function mutation. Only a very small amount of the PUF-3::GFP-derived GFP::FLAG fragment accumulates in *etc-1(tn1919)* animals (Figure 6E, see the longer exposure of lane 5), suggesting that *etc-1(tn1919)* retains a significant amount of function in the *C. elegans* germline. It is not clear why GFP::ETC-1-derived GFP::FLAG fragments do not substantially accumulate in *etc-1(tn1920)* animals. Possibly ETC-1 itself is not a substrate for proteasomal degradation. Alternatively, it is possible that these fragments accumulate to a level that is below our detection threshold. In this regard, it is worth noting that GFP::ETC-1 is expressed at much lower levels than PUF-3::GFP (Figure 6E).

#### RNP-8::GFP fusion proteins

We created a GFP-tagged allele of *rnp-8* (Figure S1C; RNP-8::GFP_F_ in all figures) and found that RNP-8::GFP is abundantly expressed in oocytes and early embryos (Figure S9A). We do not know whether this GFP-tagged allele of *rnp-8* (*rnp-8(tn1860[rnp-8::gfp:3xflag])*) is functional *in vivo*. Although RNP-8 is thought to be degraded in response to oocyte maturation (Kim *et al*. 2010), RNP-8::GFP is not strongly down-regulated during the oocyte-to-embryo transition (Figure S9A). GFP fluorescence is not obviously increased in *rnp-8(tn1860*); *etc-1(gk5182)* animals compared to *etc-1(+)* control animals (Figure S9, A and B). Nevertheless, a small proportion of the RNP-8::GFP fusion protein made by *rnp-8(tn1860*) is partially processed into GFP::FLAG fragments in *etc-1(gk5182)* hermaphrodites (Figure S7C, lanes 7 and 8); however, this might represent a low level of turnover at other stages of germline development distinct from the OET.

#### LIN-41 fusion proteins

Only full-length mSCARLET::LIN-41 fusion proteins accumulate to a substantial level in wild-type, *etc-1(gk5182)* and *sel-10(lf)* hermaphrodites (Figure 7D). We did not detect mSCARLET::FLAG or 0EN0RA::FLAG fragments at elevated levels in *etc-1(gk5182)*, *sel-10(ar41)* or *sel-10(ok1632)* animals (Figure 7D; Figure S7E). This is consistent with the observation that the levels of fluorescence appear to be the same in embryos made by wild-type and *etc-1(gk5182)* animals that express mSCARLET::LIN-41 (Figure 5, E and F) and the conclusion that the *etc-1(gk5182)* mutation does not perturb the SEL-10-mediated degradation of mSCARLET::LIN-41 or DENDRA::LIN-41 that we documented using fluorescence microscopy (Figure 1, A and B; Figure 2). We tested whether GFP::LIN-41 would behave in the same way by crossing our original GFP-tagged *lin-41(tn1541)* allele (Figure S1B; GFP_S_::LIN-41 below and in all figures) into an *etc-1(gk5182)* genetic background. However, we found that GFP is not appropriately down-regulated in the embryos of *lin-41(tn1541)*; *etc-1(gk5182)* hermaphrodites compared to *lin-41(tn1541)*; *etc-1(+)* controls (Figure S9, C and D). These observations suggest that GFP_S_::LIN-41 is partially processed in *etc-1(gk5182)* hermaphrodites and imply that GFP-tagged fusion proteins might be more recalcitrant than other fusion proteins to processivity-challenged proteasomes.

To rigorously investigate these possibilities, we created new *lin-41* alleles that express a GFP::FLAG-tagged LIN-41 fusion protein (GFP_F_::LIN-41). We isolated two independent insertions that were given different allele names [*lin-41(tn2054[gfp::3xflag::lin-41])* and *lin-41(tn2055[gfp::3xflag::lin-41])*, see Figure S1B]; these two alleles are molecularly identical and were used interchangeably in our experiments. Interestingly, GFP fluorescence in GFP_F_::LIN-41-expressing worms is much brighter than in GFP_S_::LIN-41-expressing worms (compare Figures S8, E and G), possibly because the mRNA sequence that encodes the GFP::3xFLAG tag has a much higher codon adaptation index (CAI=0.92) than the sequence that encodes the GFP::S tag (CAI=0.28) (Redemann *et al*. 2011). More than half of the GFP_F_::LIN-41-expressing animals also exhibit a defect in egg laying (Egl phenotype) as adult hermaphrodites at 20°C (Figure S9H). A similar phenotype is observed when LIN-41 is over-expressed in somatic cells during the fourth larval (14) stage by interfering with *let-7* microRNA regulation (e.g., using the *lin-41(xe11*gf) allele described by Ecsedi *et al*. 2015). With respect to our tagged alleles of *lin-41*, however, this defect uniquely characterizes the GFP::FLAG tagged alleles, as most GFP_S_::LIN-41 (Figure S9F), mSCARLET::LIN-41, and DENDRA::LIN-41-expressing adults do not have an Egl phenotype. Despite these intriguing differences, the temporal expression pattern and regulation of GFP_F_::LIN-41 strongly resembles GFP_S_::LIN-41. Most importantly, GFP_F_::LIN-41-expressing animals exhibit higher levels of GFP fluorescence in the early embryos made by *sel-10(ok1632)* and *etc-1(gk5182)* mutants relative to otherwise wild-type hermaphrodites (Figure 5, K and L; Figure S9, I-N). Interestingly, the subcellular localization pattern of GFP_F_::LIN-41-derived GFP fluorescence is distinctly different in the early embryos made by *sel-10(ok1632)* and *etc-1(gk5182)* mutant hermaphrodites (Figure S9, K and M). In *sel-10(ok1632)* embryos, GFP fluorescence is primarily cytoplasmic and appears to be enriched in ribonucleoprotein particles called P granules (Figure S9M). Localization to embryonic P granules is also observed when LIN-41 degron sequences are disrupted in GFP_S_::LIN-41 (Spike *et al*. 2018). In *etc-1(gk5182)* embryos, however, GFP fluorescence does not localize to P granules and is slightly enriched in the nucleus (Figure S9K). These observations are consistent with the idea that GFP_F_::LIN-41 is partially processed into a GFP-containing protein fragment that is largely responsible for the excess fluorescence observed in early *etc-1(gk5182)* embryos.

The only difference between GFP_F_:::LIN-41, mSCARLET::LIN-41 and DENDRA::LIN-41 is the fluorescent protein fused to LIN-41 (Figure S1B). We examined all of these proteins in *etc-1(gk5182)* and *etc-1(+)* hermaphrodites by western blot, and confirmed that GFP::FLAG fragments accumulate in GFP_F_:::LIN-41-expressing *etc-1(gk5182)* adults at elevated levels relative to *etc-1(+)* animals (Figure 7E). A longer exposure of the same western blot revealed low levels of GFP::FLAG fragments even in *etc-I(+)* hermaphrodites (Figure S7F, lane 5). However, *etc-I*-dependent fragments derived from the mSCARLET::LIN-41 and DENDRA::LIN-41 fusion proteins were not observed. We conclude that *etc-I(gk5I82)* specifically disrupts the proteolysis of the GFP-containing LIN-41 fusion proteins (GFP_F_::LIN-41 and GFP_S_::LIN-41) and speculate that tagging LIN-41 with GFP creates a LIN-41 fusion protein that is more difficult to degrade. Consistent with this idea, GFP is known to be a particularly stable fluorescent protein that can be difficult for the proteasome to degrade (Khmelinskii *et al*. 2016; Reichard *et al*. 2016; Bragan9a and Kraut 2020). Based on these observations, we propose that fusion with GFP is an important cause of the proteolysis defects observed in *etc-I(gk5I82)* animals (Figures 5 and 7). In this context, we note the study that identified and described ETC-1 as a target-specific E3 ubiquitin ligase in *C. elegans* (Wang *et al*. 2013) relied on GFP fusion proteins and monitored protein degradation by looking exclusively at fluorescence. Our observations suggest, instead, that the *C. elegans etc-I* gene has the same type of activity described for orthologous genes in yeast and humans (Aviram and Kornitzer 2010; Martinez-Noel *et al*. 2012; Chu *et al*. 2013) and that the ETC-1 protein promotes proteasome processivity, which may be especially important during the OET when many proteins are targeted for efficient proteasomal degradation.

## DISCUSSION

### Proteolysis and translational repression limit the expression of RNA-binding proteins during the OET

Many of the oocyte-expressed proteins degraded during the OET in *C. elegans* are RNA-binding proteins that regulate translation (reviewed by Robertson and Lin 2013, 2015). These RNA-binding proteins include LIN-41 and the Pumilio/FBF-family RNA-binding proteins PUF-3 and PUF-11. PUF-3 and PUF-11 function redundantly, repress the translation of specific targets in oocytes, and play important roles in germline stem cells and embryonic development in *C. elegans* (Hubstenberger *et al*. 2012; Haupt *et al*. 2020). Since PUF-3 and PUF-11 appear to be degraded soon after meiotic maturation (Figure 3), it seems likely that the maternal-effect embryonic lethality caused by an absence of both proteins is due to mRNA translation defects in developing oocytes. Interestingly, the timing and regulation of PUF-3 and PUF-11 degradation appear to be temporally and molecularly coordinated with meiotic maturation and strongly resemble what we have described for LIN-41 (Spike *et al*. 2018 and Figure 3). The resemblances include a requirement for active CDK-1 (Figure 4), which is essential for entry into meiotic M-phase (Boxem *et al*. 1999, Burrows *et al*. 2006), and independence from APC ubiquitin ligase subunits that promote the subsequent metaphase-to-anaphase transition (Furuta *et al*. 2000; Shakes *et al*. 2003). Additional connections between LIN-41, PUF-3, and PUF-11 include the identification of each protein and mRNA in purified LIN-41 RNPs, and the identification of LIN-41 and PUF-3/11 as factors important for the 3’UTR-mediated translational repression of a common target, the Rbfox-related RNA-binding protein SPN-4 (Hubstenberger *et al*. 2012; Tsukamoto *et al*. 2017). These disparate observations suggest that LIN-41, PUF-3, and PUF-11 might collaborate and function, at least in part, as components of a regulatory RNP to inhibit the translation of SPN-4 in developing oocytes. Previously, we proposed that LIN-41 is inactivated as a translational repressor prior to its degradation (Spike *et al*. 2018). Since *cdk-1(RNAi)* does not reduce or eliminate SPN-4::GFP expression in early embryos (Tsukamoto *et al*. 2017), we speculate that PUF-3 and PUF-11 are likewise inactivated prior to their degradation. Thus, we suggest that the function of a LIN-41/PUF-3/PUF-11 RNP is down-regulated through both CDK-1-dependent and -independent mechanisms near the end of oogenesis.

### ETC-I, the APC, and proteasome processivity during the OET

Wang *et al*. (2013) identified and described ETC-1 (UB*<u>E T</u>*hree *<u>C</u>*), the *C. elegans* UBE3C ortholog, as a HECT-type E3 ubiquitin ligase that promotes the degradation of the essential cell cycle proteins IFY-1/Securin and CYB-1/Cyclin B1 during the OET. Degradation was monitored using a GFP fusion protein (IFY-1::GFP or GFP::CYB-1), immunoprecipitation assays clearly documented a specific interaction between the GFP fusion protein and ETC-1, and *in vitro* assays confirmed that ETC-1 is able to interact with, and ubiquitylate, each target protein. These findings were consistent at the time with the interpretation that ETC-1 functions as an E3 ubiquitin ligase. However, they are also compatible with our results that suggest instead that ETC-1 functions as an E4 enzyme that mediates ubiquitin chain elongation for the processive degradation of proteins already targeted for ubiquitin conjugation by E3 enzymes-a function shown for Hu15 in budding yeast (Crosas *et al*. 2006). When combined with the observation that *etc-1(RNAi)* enhanced the one-cell arrest phenotype of mutants with a compromised APC (e.g., *mat-1(ax161*ts)), the biochemical and cell biological observations of Wang *et al*. (2013) appeared to support a simple and sensible model: that ETC-1 and the anaphase promoting complex target the same *C. elegans* proteins (e.g., IFY-1 and CYB-1) for ubiquitination and degradation during the OET. We revisited the genetic interaction between *etc-1* and the anaphase promoting complex, at least in part, because our new results suggest that ETC-1 is not a substrate-specific E3 ligase and make it seem unlikely that ETC-1 plays a prominent role in the degradation of endogenous IFY-1 or CYB-1. Rather, our findings suggest that *etc-1* mutants have difficulty degrading GFP-fusion proteins. The fact that *etc-1(tn2077)* null and *etc-1(gk5182)* loss-of-function mutants exhibit minimal embryonic lethality is consistent with this idea. Using *etc-1(tn2077)*, a molecular null mutation that deletes the entire *etc-1* open reading frame, we confirmed that the embryo lethal phenotype of *mat-l(ax16l*ts) is substantially enhanced at 15 °C in the absence of ETC-1. However, because the embryos produced by *mat-l(ax16lts); etc-1(tn2077)* hermaphrodites are primarily multicellular, this genetic interaction may reflect a defect in mitotic cell division rather than a defect in the OET.

### The in vivo dynamics of GFP-fusion protein degradation in the absence of ETC-1

Our analyses of several different GFP fusion proteins, including GFP fusions with IFY-1 and CYB-1, provide a clear explanation for the apparent perdurance of each protein in the absence of ETC-1. It appears that the addition of a GFP moiety to several proteins that are rapidly degraded during the OET, such as IFY-1, CYB-1, PUF-3, PUF-11, or LIN-41, creates fusion proteins that are incompletely degraded in the absence of *etc-1* function. Incomplete degradation results in the accumulation of a partial GFP-containing fragment in embryos that is easily confused for the full-length GFP fusion protein when only fluorescence is examined. Evidence that the GFP moiety is a determining factor is derived from the analysis of LIN-41 fusion proteins. LIN-41 fusion proteins containing one of three different fluorescent proteins, mSCARLET, DENDRA or GFP, were examined and the only fusion protein that was incompletely degraded in an ETC-1-dependent manner was GFP::LIN-41 (Figure 7E). Our experiments do not demonstrate, however, that the fused portion of the *C. elegans* protein is unimportant for determining whether substrate processivity is substantively reduced in the absence of ETC-1. For example, the GFP::ETC-1 fusion protein made by the loss-of-function *etc-I(tnI920)* allele may not be partially processed (Figure 6E). Although it is tempting to speculate that this is because GFP::ETC-1 is not unstable during the OET (Figure 6D), some of our other experiments indicate that fragments can be generated even when proteins appear to be relatively stable. For example, RNP-8::GFP is partially processed to a small extent in *etc-I* mutants even though it is not rapidly degraded during the OET (Figures S6C and S8B). Likewise, the behaviors of PUF-3::GFP, PUF-11::GFP and CYB-1::GFP in unmated females (Figure 7B) argue for a more nuanced view. It seems likely that the proteasome helps degrade or “turn over” each GFP fusion protein to some extent in the adult germlines of unmated females and hermaphrodites. However, because each GFP-tagged protein is relatively stable prior to the OET, many fewer fragments are formed in females in the absence of *etc-I* function. We suggest that the properties conferred by each fused *C. elegans* protein likely explain why some GFP fusion proteins appear to be less sensitive to *etc-I* function than others, even when both proteins are rapidly degraded during the OET (e.g.: IFY-1::GFP and PUF-3::GFP in Figures 5 and 7). Consistent with this idea, sequences outside the GFP moiety have been shown to be important for the Hu15-dependent proteolytic processing of a GFP-Pc15 fusion protein in *S. cerevisiae* (Aviram and Kornitzer 2010), and it is becoming increasingly] clear that the method and type of substrate ubiquitination influences proteasome processivity (Reichard *et al*. 2016; Bragan;;a and Kraut 2020).

## Conclusion

Our analysis of ETC-1 updates the literature to show that it plays a conserved role to promote proteasome processivity. Proteasome processivity may be critically important during the OET when many oocyte regulatory proteins are rapidly turned over. Our experiments also provide a cautionary note on the need to analyze endogenous proteins where possible, utilize multiple protein tags and, ideally, analyze proteins on western blots when studying protein degradation *in vivo*. Fortunately, the ease and rapidity of genome editing continues to revolutionize the ability to conduct such experiments in the *C. elegans* system (reviewed by Nance and Frnkjaer-Jensen 2019).

## ACKNOWLEDGMENTS

We thank the two anonymous reviewers for constructive suggestions, including the suggestion that we analyze an *etc-I* molecular null mutation. We are grateful to Bob Goldstein, Judith Kimble, Geraldine Seydoux, and Marcus Vargas for providing strains or reagents. Some strains were provided by the Caenorhabditis Genetics Center, which is funded by grant P4000010440 from the NIH 0ffice of Research Infrastructure Programs. We also thank WormBase for sequences and annotations. Gabriela Huelgas-Morales and Todd Starich provided helpful suggestions during the course of this work. This work was supported by NIH grant GM144029 (to 0.G.).

## LEGENDS TO SUPPLEMENTAL FIGURES

Figure S1. Alleles used in this study. (A) The exon-intron structures and allele names of the *sel-10* alleles used in this study. Exons are shown as open boxes. Colored regions of the *sel-10* exons encode the conserved F box (orange) and 7 WD repeat (gray) domains found in SEL-10. The GFP and FLAG-encoding regions added to the GFP-tagged alleles are colored in green and cyan, respectively. (B) The exon-intron structures and allele names of the *lin-41* alleles used in this study. Colored regions of the *lin-41* exons encode the conserved RING (yellow), B-box (orange), BBC (blue), Ig/filamin (purple) and NHL repeats (red) domains found in LIN-41. Fusion protein tags include GFP (green), S tag (gray), mSCARLET-I (brick red), FLAG tag (cyan) and DENDRA (green to red color gradient). (C) The exon-intron structures and allele names of *puf-3::gfp::3xflag*, *puf-11::gfp::3xflag*, *cyb-1::gfp::3xflag* and *rnp-8::gfp::3xflag*. The Pumilio-like repeats of PUF-3 and PUF-11 are in yellow. The Cyclin-N and Cyclin-C motifs of CYB-1 are in blue and orange, respectively. The RNA recognition motif (RRM) present in RNP-8 is in purple. As in prior panels, the GFP and FLAG protein tags are in green and cyan, respectively.

Figure S2. GLD-1 is degraded normally in *sel-10(tn1875[gfp::3xflag::sel-10])* animals. Dissected wild-type (A and C), *sel-10(tn1875[gfp::3xflag::sel-10])* (B) and *sel-10(tn1816[sel-10::gfp::3xflag])* (D) gonads (solid outlines) that have been fixed and stained with an anti-GLD-1 antibody. Each gonad is oriented so that the distal mitotic region is on the left and the proximal oocyte-containing region is on the right. Two different wild-type gonads are shown because slightly different staining and imaging conditions were used for each experiment; wild-type image A pairs with *sel-10(tn1875[gfp::3xflag::sel-10])* (A-B) and wild-type image C pairs with *sel-10(tn1816[sel-10::gfp::3xflag])* (C and D). Scale bars, 50 μm.

Figure S3. The elimination of PUF-3/11::GFP proteins during the OET. (A-C) GFP expression in *puf-ll(tn1824[puf-ll::gfp::3xflag])* (A), *puf-3(tn1820[puf-3::gfp::3xflag])* (B) and *puf-ll(tn1824); puf-3(tn1820)* (C) animals. GFP-positive embryos are outlined in green. The position of the spermatheca (sp) is indicated (A). All images were collected using the same settings. (D and E) Quantification of GFP fluorescence levels in arbitrary units across the distal and proximal gonad for the three genotypes analyzed: *puf-ll(tn1824[puf-ll::gfp::3xflag])* (magenta); *puf-3(tn1820[puf-3::gfp::3xflag])* (blue); and *puf-ll(tn1824); puf-3(tn1820)* (black). These quantitative data indicate that PUF-11::GFP accounts for the expression in the distal gonad, whereas both PUF-11::GFP and PUF-3::GFP are expressed at similar levels in oocytes. Scale bars, 100 μm.

Figure S4. Degradation of PUF-3::GFP and PUF-11::GFP during the OET do not require function of an Skp, Cullin, F-box (SCF)-containing E3 ubiquitin ligase or the anaphase-promoting complex. PUF-3::GFP (A-D) or PUF-11::GFP (G-J) were analyzed after empty vector control RNAi (A and G), *cul-l(RNAi)* (B and H), *skr-l/2(RNAi)* (C and I), or *mat-l(RNAi)* (D and J). Additionally, PUF-3::GFP was analyzed at 25°C in *emb-30(tn377*ts); *puf-3(tn1820[puf-3::gfp::3xflag])* (F), with *puf-3(tn1820[puf-3::gfp::3xflag])* as the control (E). The position of the spermatheca (sp) is indicated. Scale bars, 100 μm.

Figure S5. GFP-expressing alleles of *etc-1*. (A) The exon-intron structures and allele names of the *gfp::3xflag::etc-1* alleles used in this study. Colors and labels are as described for Figure 6, with the addition of exons (open boxes) that encode the GFP (green) and FLAG (cyan) tags. (C and D) GFP fluorescence in *etc-1(tn1919[gfp::3xflag::etc-1#l})* (C) and *etc-1(tn1920[gfp::3xflag::etc-1#2})* (D) hermaphrodites. These are larger versions of the images shown in Figure 7 that include the region of somatic GFP expression in the head (arrow) that is significantly brighter in *etc-1(tn1920)* hermaphrodites. GFP expression in the nucleus of an intestinal cell is also evident in the focal plane of the *etc-1(tn1919)* image (arrowhead). Additional somatic cells also express GFP::ETC-1, but most of these cells were not identified. Scale bars, 100 μm.

Figure S6. Genetic interaction between *etc-1(tn2077)* and *mat-l(ax16l*ts). (A-D and A’-D’) DIC images of day-1 (25°C) and day-2 (15°C) adult hermaphrodites of the indicated genotypes. Panels (A’-D’) show increased magnification (1.5X) of the worms shown in panels (A-D) to focus on embryos within the uterus. Arrowheads indicate multicellular embryos. The position of the spermatheca (sp) is indicated. Scale bars, 50 μm.

Figure S7. Additional western blots related to Figure 7. (A) A western blot that repeats the experiment shown in Figure 7A and includes wild type and *etc-1(gk5182)* lanes with no GFP::FLAG fusion protein. (B) PUF-3::GFP, PUF-11::GFP, and CYB-1:: GFP fusion proteins (lines) and *etc-1(gk5182)*-specific GFP::FLAG fragments (arrows) are detected by an affinity purified antibody that recognizes GFP. (C and D) Western blots with the anti-FLAG antibody on GFP fusion protein-expressing hermaphrodites (C), mated *fog-2(oz40)* females (C), unmated *fog-2(oz40)* females (D) and *fog-2(oz40)* males (D) with the indicated *etc-1* genotypes. For unknown reasons, the RNP-8::GFP fusion protein detected in (C) appears to be slightly larger than predicted. The predicted size of the large RNP-8a isoform (see http://www.wormbase.org, release WS278) fused to the GFP::3xFLAG tag is 97 kDa. The blots shown in (D) and Figure 7B are the same; Figure 7B shows a shorter exposure of the portion of the blot with the female lysates and a longer exposure of the blot with the male CYB-1::GFP lysates. (E) A longer exposure of the western blot shown in Figure 7D that also includes otherwise wild-type, *etc-1(gk5182)* and *sel-10(lf)* animals that express DENDRA::LIN-41. (F) A longer exposure of the western blot shown in Figure 7E. *etc-1(+)* animals that express GFP_F_::LIN-41 appear to accumulate a lesser amount of the GFP::FLAG fragment seen in *etc-1(gk5182)* animals. *etc-1(gk5182)*-specific fragments containing GFP::FLAG are indicated by arrows.

Figure S8. CYB-1::GFP expression in males. (A-F) CYB-1::GFP expression in *etc-1(+)* males (A) and *etc-1(gk5182)* males (C and F). CYB-1::GFP disappears abruptly in *cyb-l(tn1806[cyb-l::gfp::3xflag]); fog-2(oz40)* males (A and B) near the end of spermatogenesis (red boxes). At the same stage of spermatogenesis, *etc-1(gk5182)*; *cyb-l(tn1806[cyb-l::gfp::3xflag]); fog-2(oz40)* males (C-F) have higher levels of GFP fluorescence (C) and GFP-positive residual bodies (e.g., arrows in E and F). Scale bars, 50 μm (A-D) and 10 μm (E and F).

Figure S9. Additional images of GFP-tagged fusion proteins. (A and B) GFP is expressed at similar levels in the germlines (solid outlines) and early embryos (dashed outlines) of *rnp-8(tn1860[rpn-8::gfp::3xflag])* (A) and *etc-1(gk5182); rnp-8(tn1860[rpn-8::gfp::3xflag])* (B) hermaphrodites. (C and D) GFP fluorescence from *lin-41(tn1541[gfp::s::lin-41])* is distinctly brighter in the early embryos (dashed outlines) of *etc-1(gk5182)* hermaphrodites (D) relative to *etc-1(+)* hermaphrodites (C). (E-H) The GFP_S_-tagged allele *lin-41(tn1541[gfp::s::lin-41])* (E) and GFP_F_-tagged allele *lin-41(tn2054[gfp::3xflag::lin-41])* (G) have different levels of GFP expression. The average GFP fluorescence intensity in the cytoplasm of the −1 and −2 oocytes of *lin-41(tn2054)* and *lin-41(tn1541)* Day 1 adult hermaphrodites was measured (n=7 images for each allele, collected as shown). At each oocyte position (−1 or −2), the average background-corrected level of fluorescence was 2.4-fold higher in *lin-41(tn2054)* oocytes relative to *lin-41(tn1541)* oocytes (*P* < 1 x 10^-6^). Many *lin-41(tn2054[gfp::3xflag::lin-41])* Day 1 adult hermaphrodites also accumulate older embryos in their uterus (H) and appear to be sluggish. These phenotypes are shared by the independently-generated *lin-41(tn2055[gfp::3xflag::lin-41]* allele. At 20 °C, 43% and 62% of DG5263 *lin-41(tn2055)* Day 1 and Day 2 adults (n=29), respectively, had an egg laying (Egl) defect associated with premature mortality (dead on Day 2, 3, or 4 of adulthood) and a reduced number of progeny (52 ± 21 larvae, n=18 Egl adults; 256 ± 61 larvae, n=11 non-Egl adults). These somatic defects are not typically seen in *lin-41(tn1541[gfp::s::lin-41])* animals at the same stage of adulthood (F), and the average brood size of *lin-41(tn1541)* animals is greater than 300 (Spike *et al*. 2014a, 2018). (I-N) GFP fluorescence (I, K, and M) and DIC (J, L, and N) images of 2-cell *lin-41(tn2055[gfp::3xflag::lin-41])* embryos produced by otherwise wild-type (I and J), *etc-1(gk5182)* (K and L) and *sel-10(ok1632)* (M and N) parents. GFP is diffusely cytoplasmic in each embryo and is distinctly brighter in the *etc-1(gk5182)* and *sel-10(ok1632)* embryos relative to the wild-type embryo. Nuclear GFP (arrowheads) is evident in wild-type (I) and *etc-1(gk5182)* (K) embryos. In contrast, some GFP in the posterior blastomere of the *sel-10(ok1632)* embryo (M) appears to associate with P granules (arrow). The position of the spermatheca (sp) is indicated. Scale bars, 50 μm (A-H) and 10 μm (I-N).

Table S1. *C. elegans* strains used in this study.

File S1. Information on the repair templates, single guide RNA plasmids, PCR primers, and the RNAi clones used in this study.

File S2. The nucleotide and amino acid sequences of fluorophore tags, tagged alleles, and deletion alleles used in this study.

## Notes

### Competing Interest Statement

The authors have declared no competing interest.

### Summary of Updates

We have incorporated constructive suggestions of two anonymous reviewers.

