## Supplemental figures, table, and figure legends for "Ubiquitin ligases and a processive proteasome facilitate protein clearance during the oocyte-to-embryo transition in *Caenorhabditis elegans*"

Contents:

9 Figures

1 Table

2 Files

[1]Corresponding author: David Greenstein, Department of Genetics, Cell Biology, and Development, University of Minnesota, 4-208 MCB, 420 Washington Avenue SE, Minneapolis, MN 55455. Tel: 612-624-3955; FAX: 612-626-6140.

Table S1. *C. elegans* strains used in this study.

File S1. Information on the repair templates, single guide RNA plasmids, PCR primers, and the RNAi clones used in this study.

File S2. The nucleotide and amino acid sequences of fluorophore tags, tagged alleles, and deletion alleles used in this study.

### Figure S1

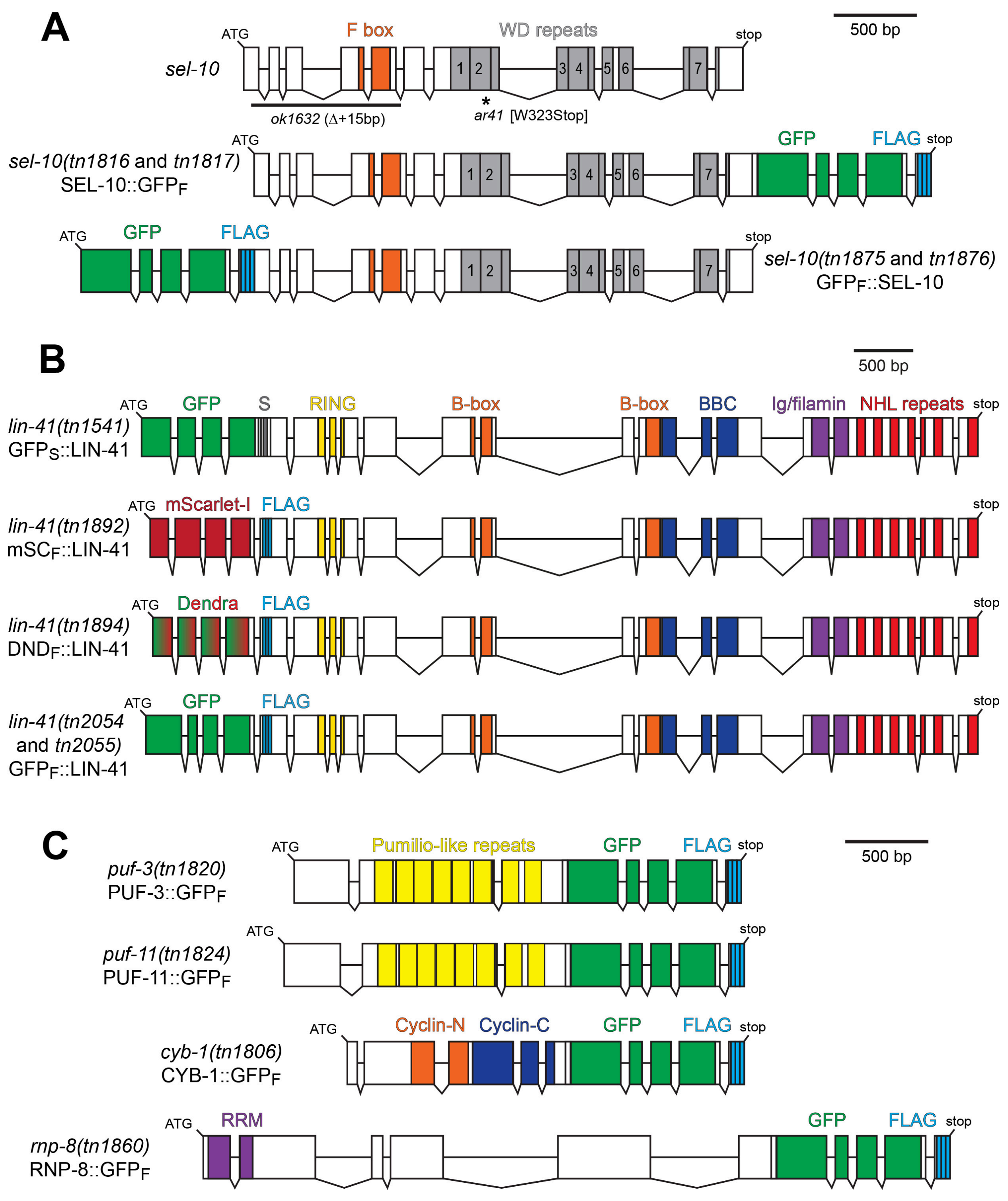


138

139

### Figure S2

140 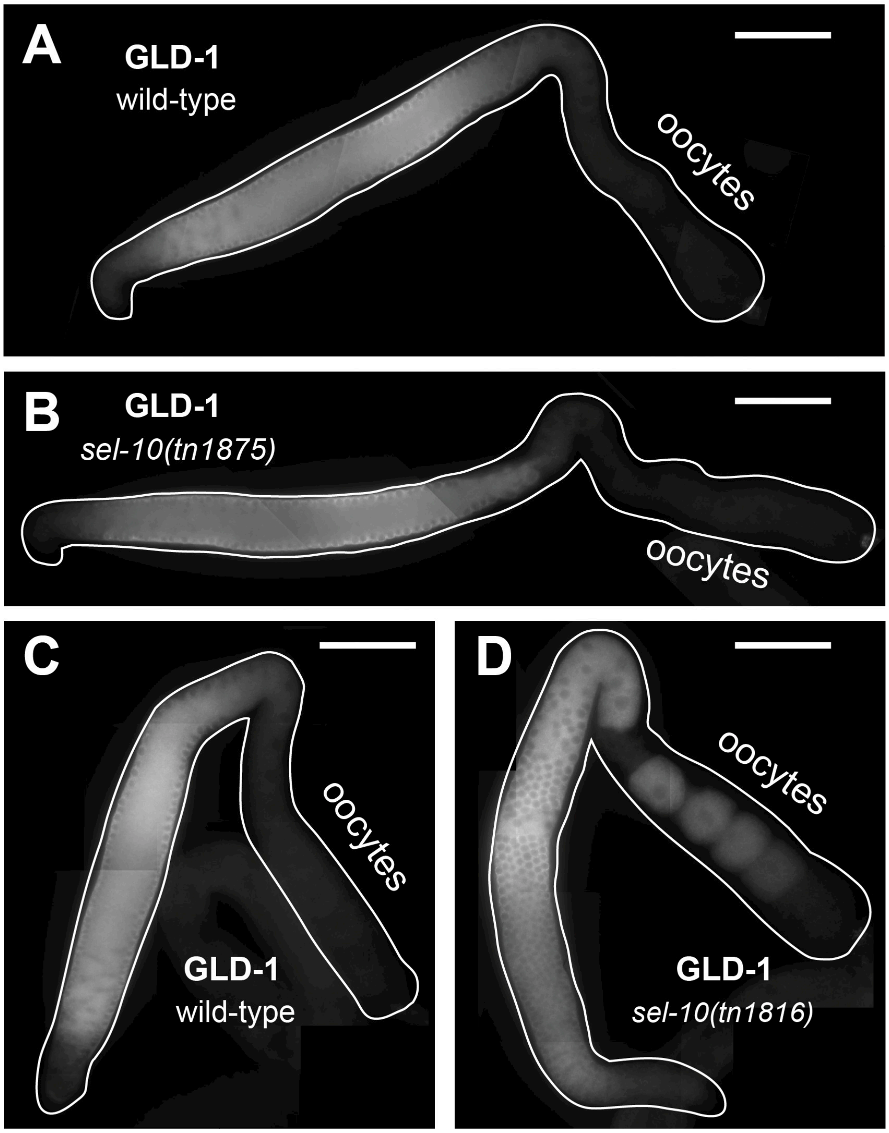


141

### Figure S3

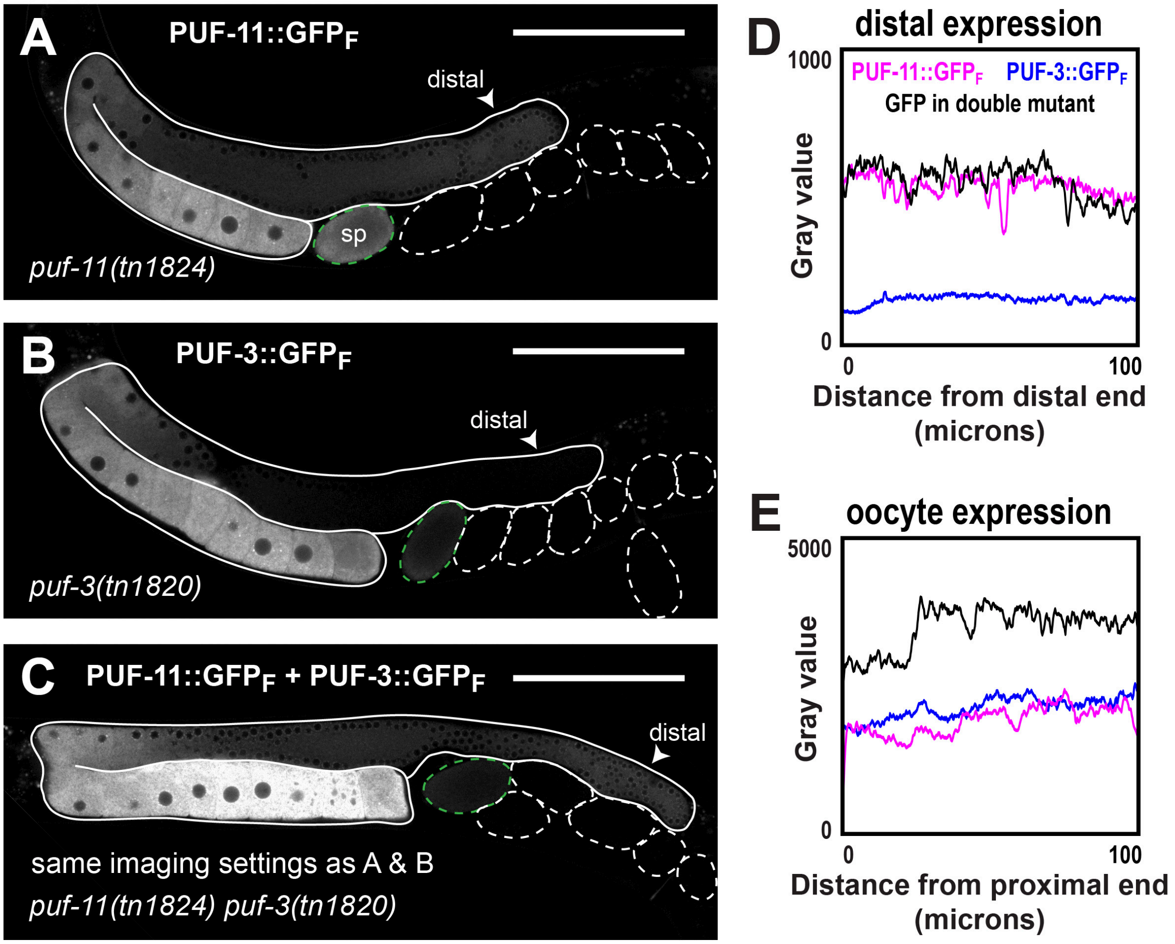


142

143

### Figure S4

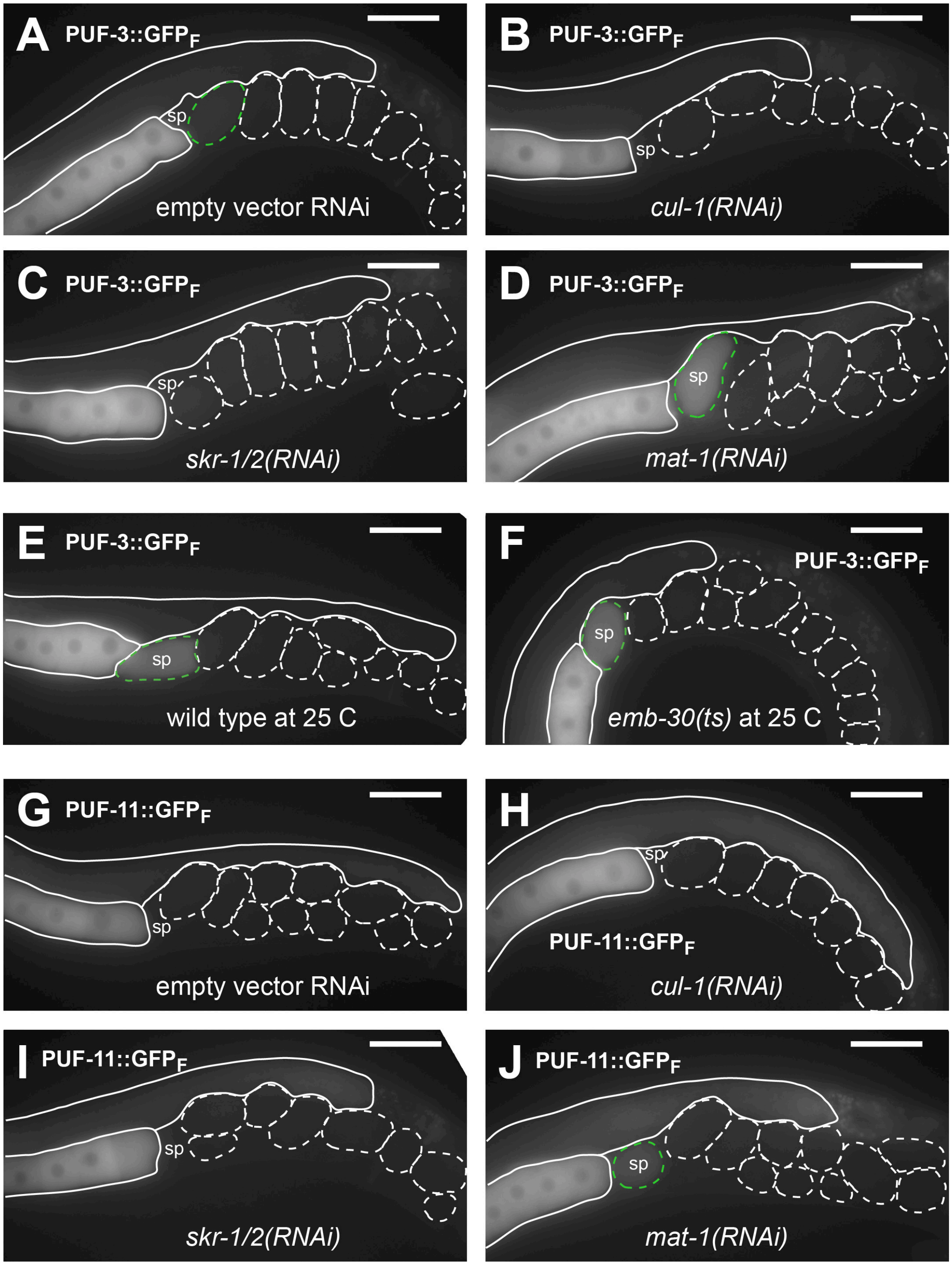
A
PUF-3::GFP$_F$
sp
empty vector RNAi
B
PUF-3::GFP$_F$
sp
*cul-1(RNAi)*
C
PUF-3::GFP$_F$
sp
*skr-1/2(RNAi)*
D
PUF-3::GFP$_F$
sp
*mat-1(RNAi)*
E
PUF-3::GFP$_F$
sp
wild type at 25 C
F
PUF-3::GFP$_F$
sp
*emb-30(ts)* at 25 C
G
PUF-11::GFP$_F$
sp
empty vector RNAi
H
sp
PUF-11::GFP$_F$
*cul-1(RNAi)*
I
PUF-11::GFP$_F$
sp
*skr-1/2(RNAi)*
J
PUF-11::GFP$_F$
sp
*mat-1(RNAi)*

### Figure S5

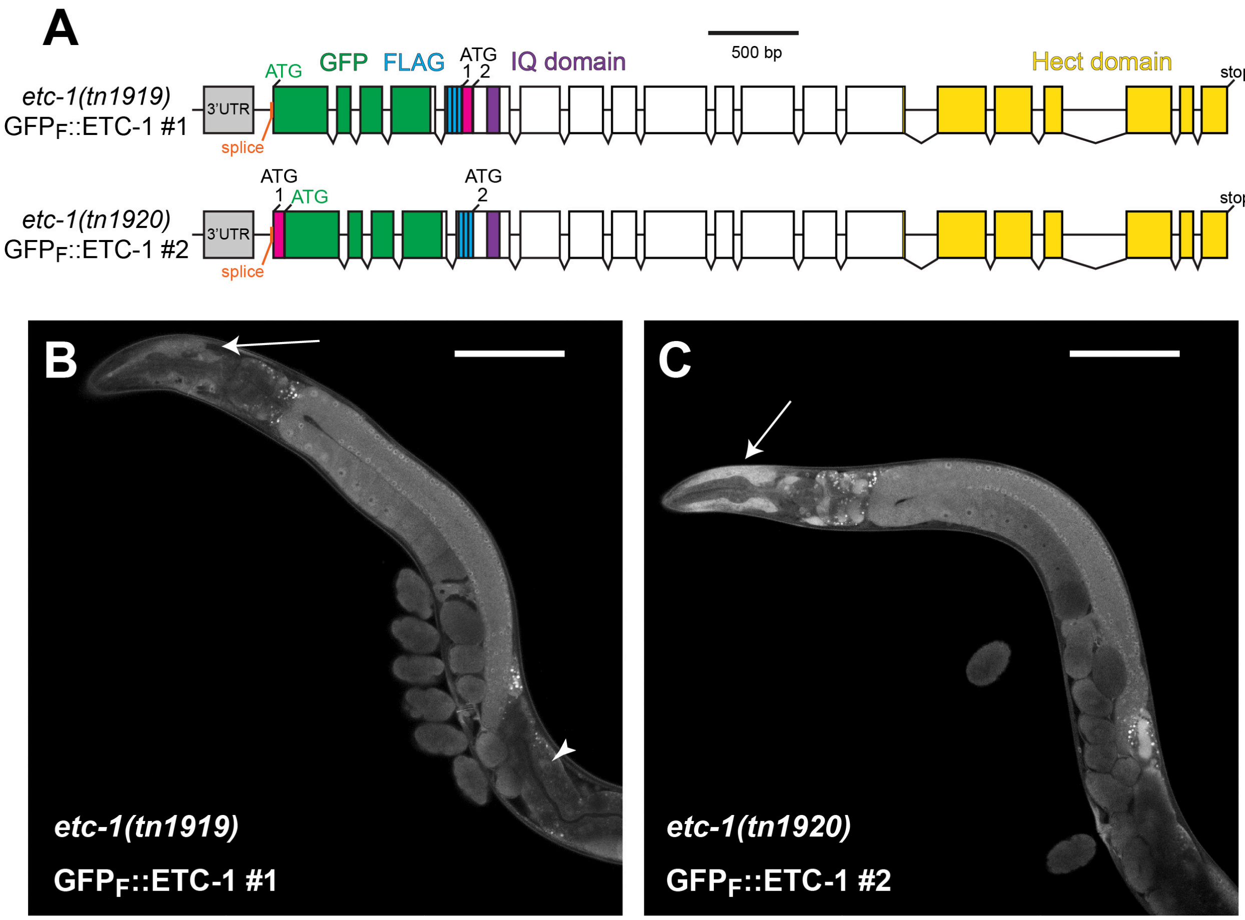
A
B
C
500 bp
ATG
GFP
FLAG
ATG
1
2
IQ domain
Hect domain
stop
etc-1(tn1919)
GFPF::ETC-1 #1
3'UTR
splice
ATG
1
ATG
ATG
2
stop
etc-1(tn1920)
GFPF::ETC-1 #2
3'UTR
splice
etc-1(tn1919)
GFPF::ETC-1 #1
etc-1(tn1920)
GFPF::ETC-1 #2

### Figure S6

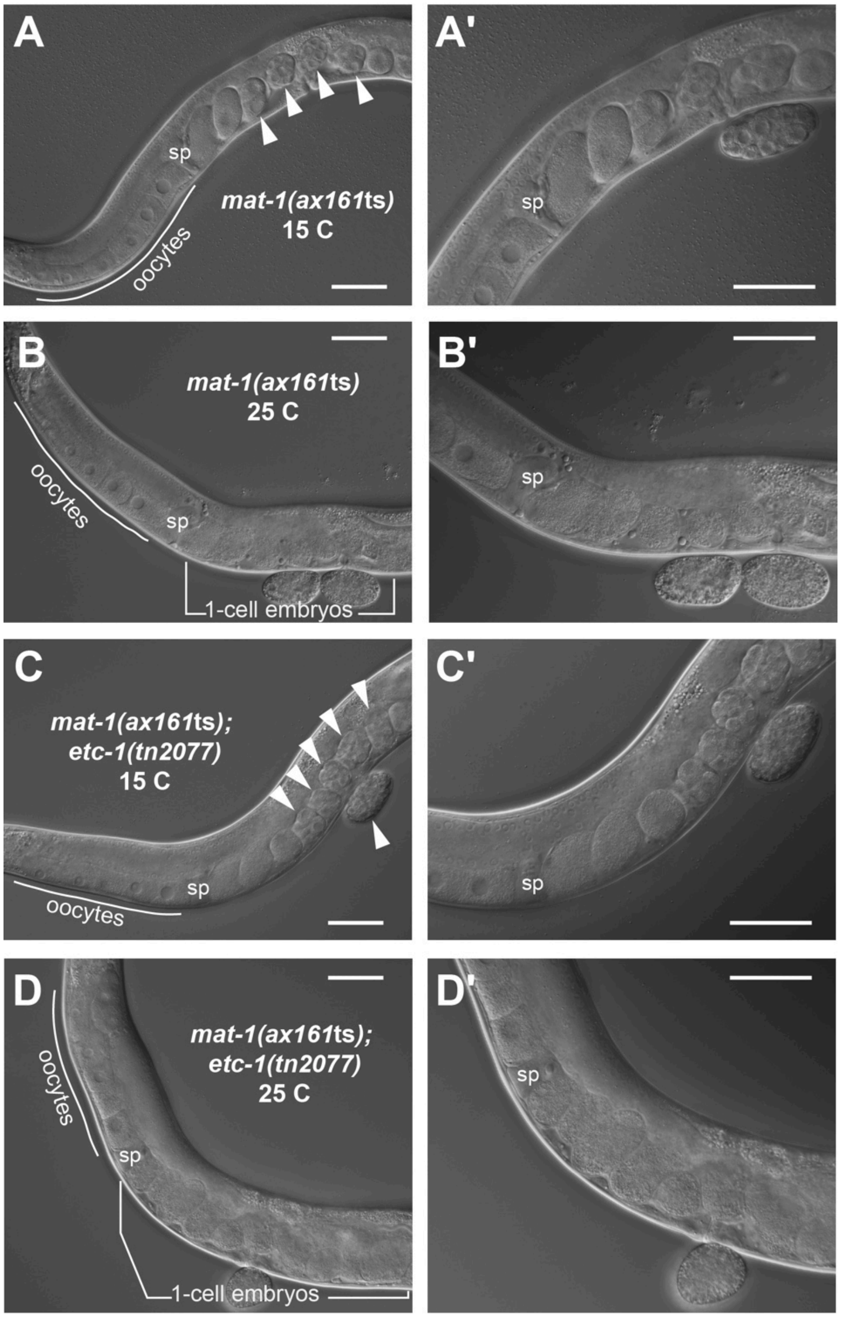
A
sp
mat-1(ax161ts)
15 C
oocytes
A'
sp
B
mat-1(ax161ts)
25 C
oocytes
sp
1-cell embryos
B'
sp
C
mat-1(ax161ts);
etc-1(tn2077)
15 C
sp
oocytes
C'
sp
D
oocytes
mat-1(ax161ts);
etc-1(tn2077)
25 C
sp
1-cell embryos
D'
sp

### Figure S7

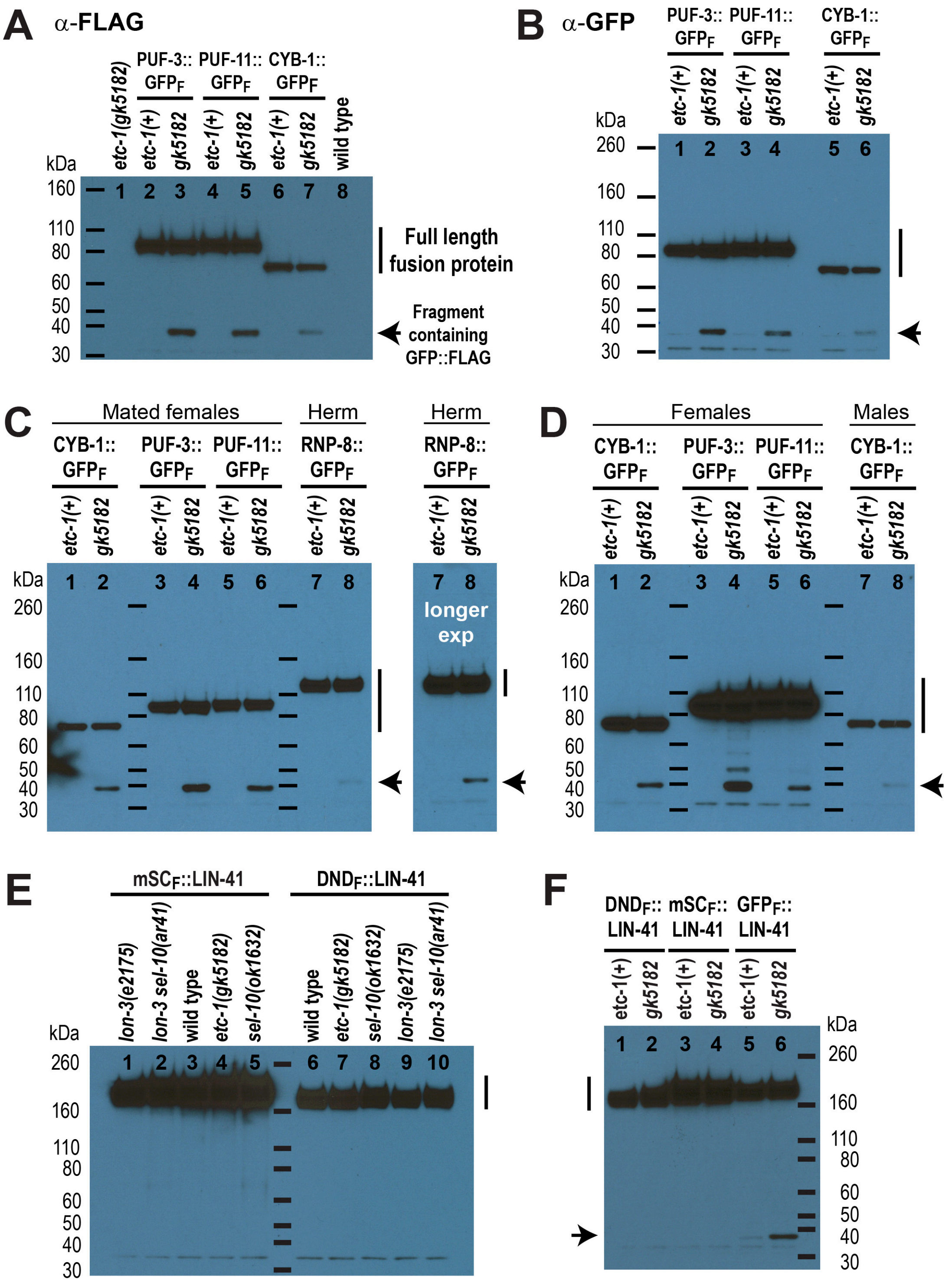
A
α-FLAG
PUF-3:: GFPF
PUF-11:: GFPF
CYB-1:: GFPF
etc-1(gk5182)
etc-1(+)
gk5182
wild type
kDa
160
110
80
60
50
40
30
Full length fusion protein
Fragment containing GFP::FLAG
B
α-GFP
PUF-3:: GFPF
PUF-11:: GFPF
CYB-1:: GFPF
etc-1(+)
gk5182
kDa
260
160
110
80
60
50
40
30
C
Mated females
Herm
CYB-1:: GFPF
PUF-3:: GFPF
PUF-11:: GFPF
RNP-8:: GFPF
etc-1(+)
gk5182
kDa
260
160
110
80
60
50
40
30
longer exp
D
Females
Males
CYB-1:: GFPF
PUF-3:: GFPF
PUF-11:: GFPF
etc-1(+)
gk5182
kDa
260
160
110
80
60
50
40
30
E
mSCF::LIN-41
DNDF::LIN-41
lon-3(e2175)
lon-3 sel-10(ar41)
wild type
etc-1(gk5182)
sel-10(ok1632)
kDa
260
160
110
80
60
50
40
30
F
DNDF:: LIN-41
mSCF:: LIN-41
GFPF:: LIN-41
etc-1(+)
gk5182
kDa
260
160
110
80
60
50
40
30

### Figure S8

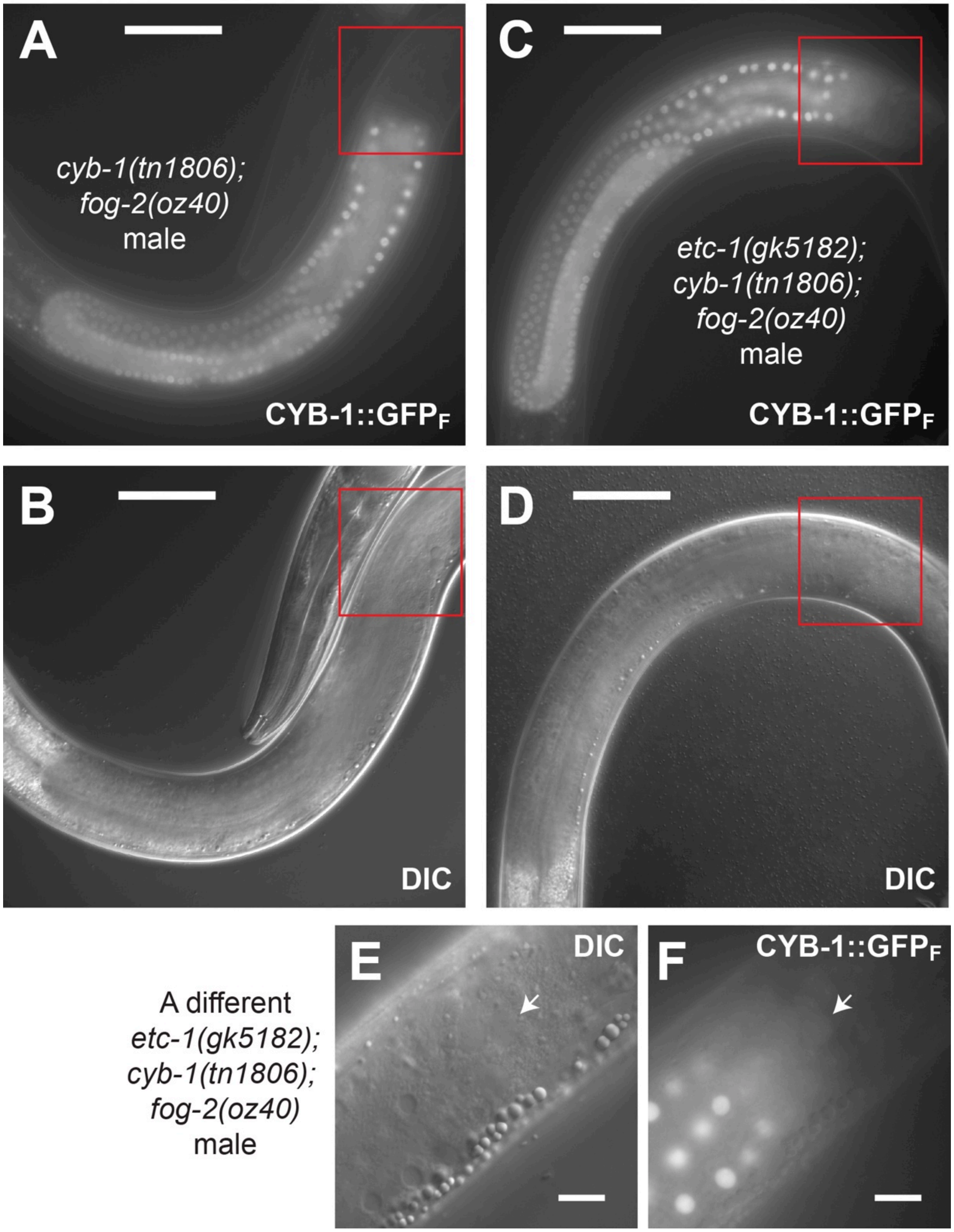

A
cyb-1(tn1806);
fog-2(oz40)
male
CYB-1::GFPF
C
etc-1(gk5182);
cyb-1(tn1806);
fog-2(oz40)
male
CYB-1::GFPF
B
DIC
D
DIC
A different
etc-1(gk5182);
cyb-1(tn1806);
fog-2(oz40)
male
E
DIC
F
CYB-1::GFPF

### Figure S9

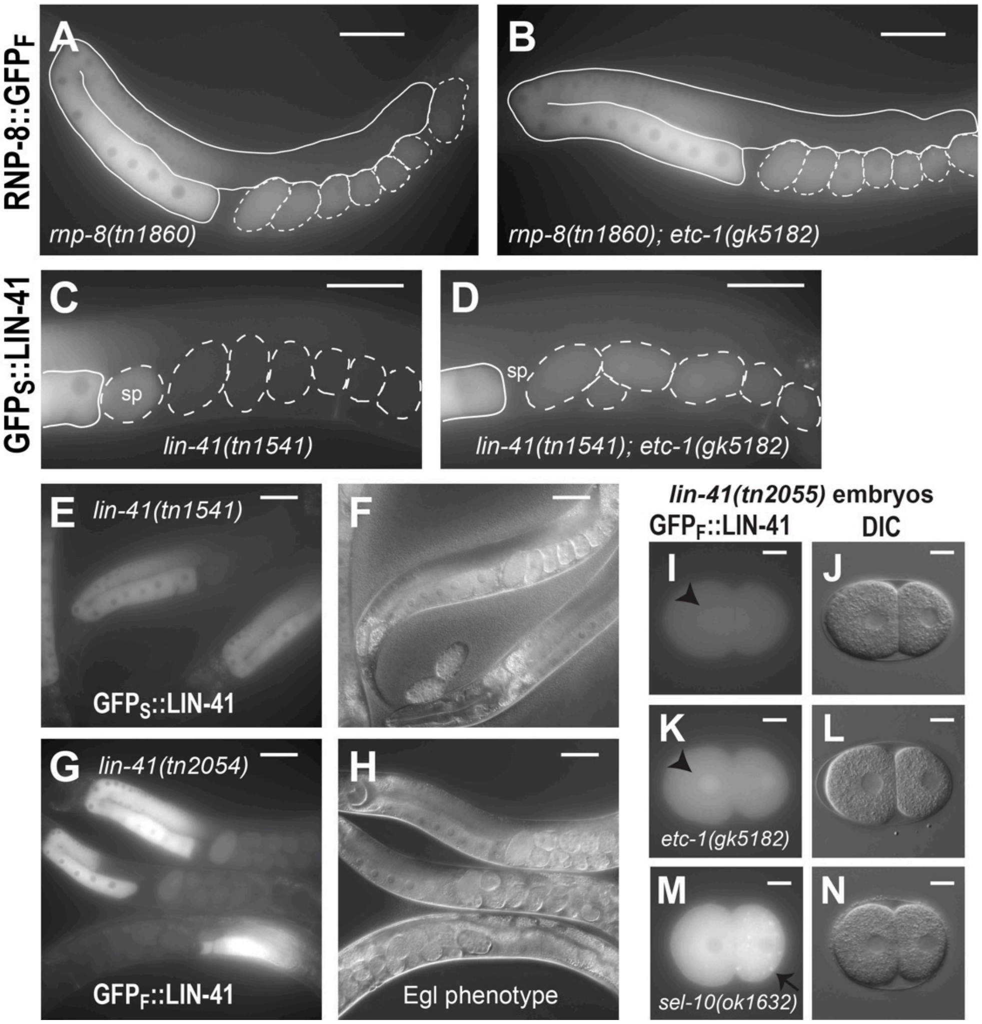
RNP-8::GFP$_F$
A
rnp-8(tn1860)
B
rnp-8(tn1860); etc-1(gk5182)
GFP$_S$::LIN-41
C
sp
lin-41(tn1541)
D
sp
lin-41(tn1541); etc-1(gk5182)
E
lin-41(tn1541)
GFP$_S$::LIN-41
F
G
lin-41(tn2054)
GFP$_F$::LIN-41
H
Egl phenotype
lin-41(tn2055) embryos
GFP$_F$::LIN-41
DIC
I
J
K
etc-1(gk5182)
L
M
sel-10(ok1632)
N

**Table S1 *C. elegans* strains used for this study**

| Strain | Genotype |
|---|---|
| N2 | Wild type, Bristol isolate |
| BS553 | *fog-2(oz40)* V |
| CB66 | *unc-22(e66)* IV |
| CB4123 | *lon-3(e2175)* V |
| CGC43 | *unc-4(e120) / mnC1[dpy-10(e128) unc-52(e444) umnIs32]* II |
| FX30168 | *tmC18[dpy-5(tmIs1236)]* I |
| GS922 | *lon-3(e2175) sel-10(ar41)* V |
| GS6156 | *sel-10(ok1632)* V |
| JK6321 | *puf-11(q971) puf-3(q966)* IV / *nT1[qIs51]* (IV;V) |
| TH214 | *unc-119(ed3) III; ddIs128[ify-1::ty1::egfp::3xflag(92C12) + unc-119(+)]* |
| VC4092 | *etc-1(gk5182)* II |
| DG627 | *emb-30(tn377*ts*)* III |
| DG3913 | *lin-41(tn1541[gfp::tev::s::lin-41])* I |
| DG4339 | *itIs37[pie-1p::mcherry::histoneH2B::pie-1 3'UTR, unc-119(+)]* IV |
| DG4569 | *cyb-1(tn1806[cyb-1::gfp::tev::3xflag])* IV |
| DG4600 | *cyb-1(tn1806[cyb-1::gfp::tev::3xflag])* IV; *fog-2(oz40)* V |
| DG4602 | *sel-10(tn1816[sel-10::gfp::tev::3xflag])* V |
| DG4603 | *sel-10(tn1817[sel-10::gfp::tev::3xflag])* V |
| DG4607 | *puf-3(tn1820[puf-3::gfp::tev::3xflag])* IV |
| DG4611 | *puf-11(tn1824[puf-11::gfp::tev::3xflag])* IV |
| DG4647 | *rnp-8(tn1860[rnp-8::gfp::tev::3xflag])* I |
| DG4669 | *puf-3(tn1820[puf-3::gfp::tev::3xflag])* IV; *lon-3(e2175)* V |
| DG4670 | *puf-3(tn1820[puf-3::gfp::tev::3xflag])* IV; *lon-3(e2175) sel-10(ar41)* V |
| DG4671 | *puf-11(tn1824[puf-11::gfp::tev::3xflag])* IV; *lon-3(e2175)* V |
| DG4672 | *puf-11(tn1824[puf-11::gfp::tev::3xflag])* IV; *lon-3(e2175) sel-10(ar41)* V |
| DG4675 | *sel-10(tn1875[gfp::tev::3xflag::sel-10])* V |
| DG4676 | *sel-10(tn1876[gfp::tev::3xflag::sel-10])* V |
| DG4689 | *puf-11(tn1824[puf-11::gfp::tev::3xflag])* IV; *fog-2(oz40)* V |
| DG4695 | *puf-3(tn1820[puf-3::gfp::tev::3xflag])* IV; *fog-2(oz40)* V |
| DG4698 | *itIs37[pie-1p::mcherry:: histoneH2B::pie-1 3'UTR, unc-119(+)] puf-3(tn1820[puf-3::gfp::tev::3xflag])* IV |
| DG4699 | *itIs37[pie-1p::mcherry:: histoneH2B::pie-1 3'UTR, unc-119(+)] puf-3(tn1820[puf-3::gfp::tev::3xflag])* IV |
| DG4718 | *emb-30(tn377*ts*)* III*; puf-3(tn1820[puf-3::gfp::tev::3xflag])* IV |
| DG4719 | *emb-30(tn377*ts*)* III*; puf-3(tn1820[puf-3::gfp::tev::3xflag])* IV |
| DG4722 | *plk-1(or683*ts*)* III; *puf-3(tn1820[puf-3::gfp::tev::3xflag])* IV |
| DG4723 | *cul-2(or209*ts*)* III; *puf-3(tn1820[puf-3::gfp::tev::3xflag])* IV |
| DG4727 | *lin-41(tn1892[mscarlet::tev::3xflag::lin-41])* I |
| DG4731 | *lin-41(tn1894[dendra::tev::3xflag::lin-41])* I |
| DG4737 | *lin-41(tn1892[mscarlet::tev::3xflag::lin-41])* I; *puf-11(tn1824[puf-11::gfp::tev::3xflag])* IV |
| DG4738 | *lin-41(tn1892[mscarlet::tev::3xflag::lin-41])* I; *puf-3(tn1820[puf-3::gfp::tev::3xflag])* IV |

| | |
|---|---|
| DG4740 | *lin-41(tn1892[mscarlet::tev::3xflag::lin-41])* I; *sel-10(tn1817[sel-10::gfp::tev::3xflag])* V |
| DG4741 | *lin-41(tn1892[mscarlet::tev::3xflag::lin-41])* I; *sel-10(tn1875[gfp::tev::3xflag::sel-10])* V |
| DG4742 | *lin-41(tn1892[mscarlet::tev::3xflag::lin-41])* I; *lon-3(e2175)* V |
| DG4743 | *lin-41(tn1892[mscarlet::tev::3xflag::lin-41])* I; *lon-3(e2175) sel-10(ar41)* V |
| DG4744 | *lin-41(tn1894[dendra::tev::3xflag::lin-41])* I; *lon-3(e2175)* V |
| DG4745 | *lin-41(tn1894[dendra::tev::3xflag::lin-41])* I; *lon-3(e2175) sel-10(ar41)* V |
| DG4751 | *lin-41(tn1892[mscarlet::tev::3xflag::lin-41])* I; *sel-10(ok1632)* V |
| DG4752 | *lin-41(tn1894[dendra::tev::3xflag::lin-41])* I; *sel-10(ok1632)* V |
| DG4790 | *etc-1(gk5182)* II (outcrossed) |
| DG4791 | *etc-1(gk5182)* II*; puf-3(tn1820[puf-3::gfp::tev::3xflag])* IV |
| DG4792 | *etc-1(gk5182)* II*; puf-11(tn1824[puf-11::gfp::tev::3xflag])* IV |
| DG4793 | *etc-1(gk5182)* II*; cyb-1(tn1806[cyb-1::gfp::tev::3xflag])* IV |
| DG4815 | *puf-11(tn1824[puf-11::gfp::tev::3xflag]) unc-22(e22)* IV |
| DG4821 | *etc-1(gk5182)* II; *lon-3(e2175)* V |
| DG4822 | *etc-1(gk5182)* II; *lon-3(e2175) sel-10(ar41)* V |
| DG4859 | *puf-11(tn1824[puf-11::gfp::tev::3xflag]) puf-3(tn1820[puf-3::gfp::tev::3xflag])* IV |
| DG4865 | *etc-1(tn1919[gfp::tev::3xflag:: etc-1(#1)])* II [tag inserted between the trans-splice site and the adjacent start codon] |
| DG4867 | *etc-1(tn1920[gfp::tev::3xflag::etc-1(#2)])* II [tag inserted before the annotated start codon] |
| DG4934 | *lin-41(tn1892[mscarlet::tev::3xflag::lin-41])* I; *etc-1(gk5182)* II |
| DG4935 | *lin-41(tn1894[dendra::tev::3xflag::lin-41])* I; *etc-1(gk5182)* II |
| DG4952 | *fog-3(q443)/tmC18[dpy-5(tmIs1236)]* I; *puf-11(q971) puf-3(q966)* IV */nT1[qIs51]* (IV;V) |
| DG4953 | *etc-1(gk5182)* II*; fog-2(oz40)* V |
| DG4965 | *etc-1(tn1919[gfp::tev::3xflag::etc-1(#1)])* II; *puf-3(tn1820[puf-3::gfp::tev::3xflag]*) IV |
| DG4966 | *etc-1(tn1920[gfp::tev::3xflag::etc-1(#2)])* II; *puf-3(tn1820[puf-3::gfp::tev::3xflag]*) IV |
| DG4975 | *rnp-8(tn1860[rnp-8::gfp::tev::3xflag])* I; *etc-1(gk5182)* II |
| DG4978 | *etc-1(gk5182)* II; *cyb-1(tn1806[cyb-1::gfp::tev::3xflag])* IV*; fog-2(oz40)* V |
| DG4980 | *etc-1(gk5182)* II; *puf-3(tn1820[puf-3::gfp::tev::3xflag])* IV*; fog-2(oz40)* V |
| DG4983 | *etc-1(gk5182)* II; *puf-11(tn1824[puf-11::gfp::tev::3xflag])* IV*; fog-2(oz40)* V |
| DG5060 | *lin-41(tn1541[gfp::tev::s::lin-41])* I; *etc-1(gk5182)* II |
| DG5092 | *etc-1(gk5182)* II; *ddIs128[ify-1::TY1::EGFP::3xFLAG(92C12) + unc-119(+)]* |
| DG5202 | *mat-1(ax161*ts)/*tmC18[dpy-5(tmIs1236)]* I |
| DG5204 | *mat-1(ax161*ts)/*tmC18[dpy-5(tmIs1236)]* I*; etc-1(gk5182)* II |
| DG5240 | *etc-1(gk5182)* II*; emb-30(tn377*ts*)* III |
| DG5246 | *lin-41(tn2054[gfp::tev::3xflag::lin-41])* I |
| DG5252 | *lin-41(tn2054[gfp::tev::3xflag::lin-41])* I; *etc-1(gk5182)* II |
| DG5258 | *lin-41(tn2055[gfp::tev::3xflag::lin-41])* I; *etc-1(gk5182)* II |
| DG5263 | *lin-41(tn2055[gfp::tev::3xflag::lin-41])* I |
| DG5273 | *lin-41(tn2055[gfp::tev::3xflag::lin-41])* I; *sel-10(ok1632)* V |

| | |
|---|---|
| DG5379 | *dpy-10(tn2076) etc-1(tn2077)* II |
| DG5381 | *etc-1(tn2077)* II |
| DG5400 | *mat-1(ax161*ts)/*tmC18[dpy-5(tmIs1236)]* I*; etc-1(tn2077)* II |
