## Supplementary material for "Ubiquitin ligases and a processive proteasome facilitate protein clearance during the oocyte-to-embryo transition in *Caenorhabditis elegans*": File S2

**(1) Amino acid sequences of all C-terminal tags**

(1a) GFP::3xFLAG, 32.4 kDa

Fused to the C-terminus of PUF-3/11, CYB-1, and RNP-8

GASGASGASMSKGEELFTGVVPILVELDGDVNGHKFSVSGEGEGDATYGKLTLKFICTTGKLPVPWPTLVTTFCYGVQCFSRYPDHMKRHDFFKSAMPEGYVQERTIFFKDDGNYKTRAEVKFEGDTLVNRIELKGIDFKEDGNILGHKLEYNYNSHNVYIMADKQKNGIKVNFKIRHNIEDGSVQLADHYQQNTPIGDGPVLLPDNHYLSTQSALSKDPNEKRDHMVLLEFVTAAGITHGMDELYKENLYFQSGKGAGSDYKDDDDKRDYKDDDDKRDYKDDDDKR*

(1b) 2xTY1::GFP::3xFLAG, 33.6 kDa

Fused to the C-terminus of IFY-1

EVHTNQDPLDEVHTNQDPLDTSMSKGEELFTGVVPILVELDGDVNGHKFSVSGEGEGDATYGKLTLKFICTTGKLPVPWPTLVTTFCYGVQCFSRYPDHMKQHDFFKSAMPEGYVQERTIFFKDDGNYKTRAEVKFEGDTLVNRIELKGIDFKEDGNILGHKLEYNYNSHNVYIMADKQKNGIKVNFKIRHNIEDGSVQLADHYQQNTPIGDGPVLLPDNHYLSTQSALSKDPNEKRDHMVLLEFVTAAGITHGMDELYKSGSSYSLESIGTSLEDYKDHDGDYKDHDIDYKDDDDK*

**(2) Amino acid sequences of all N-terminal tags**

(2a) GFP::S, 34.6 kDa

Fused to the N-terminus of LIN-41

MGSPGLQEFDIKLIDTVDLEGAVPVEKMSKGEELFTGVVPILVELDGDVNGHKFSVSGEGEGDATYGKLTLKFICTTGKLPVPWPTLVTTFCYGVQCFSRYPDHMKRHDFFKSAMPEGYVQERTIFFKDDGNYKTRAEVKFEGDTLVNRIELKGIDFKEDGNILGHKLEYNYNSHNVYIMADKQKNGIKVNFKIRHNIEDGSVQLADHYQQNTPIGDGPVLLPDNHYLSTQSALSKDPNEKRDHMVLLEFVTAAGITHGMDELYKCPGDRWSSTGGGRSRENLYFQGAAKFKETAAAKFERQHMDSGGGG

(2b) GFP::3xFLAG, 31.7 kDa

Fused to the N-terminus of ETC-1 and LIN-41

MSKGEELFTGVVPILVELDGDVNGHKFSVSGEGEGDATYGKLTLKFICTTGKLPVPWPTLVTTFCYGVQCFSRYPDHMKRHDFFKSAMPEGYVQERTIFFKDDGNYKTRAEVKFEGDTLVNRIELKGIDFKEDGNILGHKLEYNYNSHNVYIMADKQKNGIKVNFKIRHNIEDGSVQLADHYQQNTPIGDGPVLLPDNHYLSTQSALSKDPNEKRDHMVLLEFVTAAGITHGMDELYKENLYFQSGKGAGSDYKDDDDKRDYKDDDDKRDYKDDDDKR

(2c) mScarlet-I::3xFLAG, 31.1 kDa

Fused to the N-terminus of LIN-41

MVSKGEAVIKEFMRFKVHMEGSMNGHEFEIEGEGEGRPYEGTQTAKLKVTKGGPLPFSWDILSPQFMYGSRAFIKHPADIPDYYKQSFPEGFKWERVMNFEDGGAVTVTQDTSLEDGTLIYKVKLRGTNFPPDGPVMQKKTMGWEASTERLYPEDGVLKGDIKMALRLKDGGRYLADFKTTYKAKKPVQMPGAYNVDRKLDITSHNEDYTVVEQYERSEGRHSTGGMDELYKENLYFQSGKGAGSDYKDDDDKRDYKDDDDKRDYKDDDDKR

(2d) DENDRA::3xFLAG, 30.4 kDa

Fused to the N-terminus of LIN-41

MNLIKEDMRVKVHMEGNVNGHAFVIEGEGKGKPYEGTQTANLTVKEGAPLPFSYDILTTAVHYGNRVFTKYPEDIPDYFKQSFPEGYSWERTMTFEDKGICTIRSDISLEGDCFFQNVRFKGTNFPPNGPVMQKKTLKWEPSTEKLHVRDGLLVGNINMALLLEGGGHYLCDFKTTYKAKKVVQLPDAHFVDHRIEILGNDSDYNKVKLYEHAVARYSPLPSQAWENLYFQSGKGAGSDYKDDDDKRDYKDDDDKRDYKDDDDKR

**(3) Nucleotide sequence of the *etc-1(tn2077)* deletion allele.**

The two nucleotides that flank the *etc-1* deletion are underlined.

CAGTATTTATTTTTGTTTTGCAATTGAATATTTCAGATTTCCACTAATTAGTTTCCAATTGAAAACGTCACAAGAAAATCACTTCTCTGTAAACTATGTAAAATTTTTGTTATTGCTATCCGGATG

**(4) Nucleotide sequences of the tagged alleles illustrated in Figures S1 and S5.**

At least 70 bp of wild type flanking sequence is included on each side to indicate the position of each insertion in the *C. elegans* genome. Nucleotide sequences inserted or altered relative to the wild type are highlighted.

(4a) *sel-10(tn1816)* and *sel-10(tn1817)*

CTGGAGGCAATGGTGGCTGTATTTGGAGACTTTGTTCTACTTCTACGATGCTAGCGTGTGCAGTCGGATCTCGTAACAACACCGAGGAGACCAAAGTTATTCTCCTTGATTTCGACGCAGTTTATCCGGGAGCATCGGGAGCCTCAGGAGCATCGATGAGTAAAGGAGAAGAATTGTTCACTGGAGTTGTCCCAATCCTCGTCGAGCTCGACGGAGACGTCAACGGACACAAGTTCTCCGTCTCCGGAGAGGGAGAGGGAGACGCCACCTACGGAAAGCTCACCCTCAAGTTCATCTGCACCACCGGAAAGCTCCCAGTCCCATGGCCAACCCTCGTCACCACCTTCTGCTACGGAGTCCAATGCTTCTCCCGTTACCCAGACCACATGAAGCGTCACGACTTCTTCAAGTCCGCCATGCCAGAGGGATACGTCCAAGAGCGTACCATCTTCTTCAAGgtaagtttaaacatatatatactaactactgattatttaaattttcagGACGACGGAAACTACAAGACCCGTGCCGAGGTCAAGTTCGAGGGAGACACCCTCGTCAACCGTATCGAGCTCAAGgtaagtttaaacagttcggtactaactaaccatacatatttaaattttcagGGAATCGACTTCAAGGAGGACGGAAACATCCTCGGACACAAGCTCGAGTACAACTACAACTCCCACAACGTCTACATCATGGCCGACAAGCAAAAGAACGGAATCAAGGTCAACTTCAAGgtaagtttaaacatgattttactaactaactaatctgatttaaattttcagATCCGTCACAACATCGAGGACGGATCCGTCCAACTCGCCGACCACTACCAACAAAACACCCCAATCGGAGACGGACCAGTCCTCCTCCCAGACAACCACTACCTCTCCACCCAATCCGCCCTCTCCAAGGACCCAAACGAGAAGCGTGACCACATGGTCCTCCTCGAGTTCGTCACCGCCGCCGGAATCACCCACGGAATGGACGAGCTCTACAAGGAGAATCTGTACTTTCAATCCGGAAAGgtaagtttaaaataacttcgtatagcatacattatacgaagttattttcagGGAGCCGGATCTGATTATAAAGACGATGACGATAAGCGTGACTACAAGGACGACGACGACAAGCGTGATTACAAGGATGACGATGACAAGAGATAAcgaattctcgaatctctgcccctgtacatagaatgttcttgcttaggaactaatattgtacacgatgccctcatttttaaat

(4b) *sel-10(tn1875)* and *sel-10(tn1876)*

tcttctcttctcattttctcacgaaaatcatagttttatcgactttccttttgtgttcaaattcttcattcccagtagtttttgtcctttattcatttccaattctttttcagccatATGAGTAAAGGAGAAGAATTGTTCACTGGAGTTGTCCCAATCCTCGTCGAGCTCGACGGAGACGTCAACGGACACAAGTTCTCCGTCTCCGGAGAGGGAGAGGGAGACGCCACCTACGGAAAGCTCACCCTCAAGTTCATCTGCACCACCGGAAAGCTCCCAGTCCCATGGCCAACCCTCGTCACCACCTTCTGCTACGGAGTCCAATGCTTCTCCCGTTACCCAGACCACATGAAGCGTCACGACTTCTTCAAGTCCGCCATGCCAGAGGGATACGTCCAAGAGCGTACCATCTTCTTCAAGgtaagtttaaacatatatatactaactactgattatttaaattttcagGACGACGGAAACTACAAGACCCGTGCCGAGGTCAAGTTCGAGGGAGACACCCTCGTCAACCGTATCGAGCTCAAGgtaagtttaaacagttcggtactaactaaccatacatatttaaattttcagGGAATCGACTTCAAGGAGGACGGAAACATCCTCGGACACAAGCTCGAGTACAACTACAACTCCCACAACGTCTACATCATGGCCGACAAGCAAAAGAACGGAATCAAGGTCAACTTCAAGgtaagtttaaacatgattttactaactaactaatctgatttaaattttcagATCCGTCACAACATCGAGGACGGATCCGTCCAACTCGCCGACCACTACCAACAAAACACCCCAATCGGAGACGGACCAGTCCTCCTCCCAGACAACCACTACCTCTCCACCCAATCCGCCCTCTCCAAGGACCCAAACGAGAAGCGTGACCACATGGTCCTCCTCGAGTTCGTCACCGCCGCCGGAATCACCCACGGAATGGACGAGCTCTACAAGGAGAATCTGTACTTTCAATCCGGAAAGgtaagtttaaaataacttcgtatagcatacattatacgaagttattttcagGGAGCCGGATCTGATTATAAAGACGATGACGATAAGCGTGACTACAAGGACGACGACGACAAGCGTGATTACAAGGATGACGATGACAAGAGAATGTGGCCACGTAACGACGTGCATATGGATGATGGATCGATGACACCGGAGGACCAGGAGCCTGTTACCGATAATGATATGGAATATAATgtatgttaacatttacaacccttttcagaatataatttaattctaattgaaatgtttgtcttcagGACAA

(4c) *lin-41(tn1541)*

attgttccattcgttctgaaaagtcaaaaaattcataagtattctaattgtagagtcatcgtttgccctttccactgataattaatcaaccttttcagacttggaaaaagtgaaATGGGATCCCCCGGGCTGCAGGAATTCGATATCAAGCTTATCGATACCGTCGACCTCGAGGGGGCGGTACCGGTAGAAAAAATGAGTAAAGGAGAAGAACTTTTCACTGGAGTTGTCCCAATTCTTGTTGAATTAGATGGTGATGTTAATGGGCACAAATTTTCTGTCAGTGGAGAGGGTGAAGGTGATGCAACATACGGAAAACTTACCCTTAAATTTATTTGCACTACTGGAAAACTACCTGTTCCATGGgtaagtttaaacatatatatactaactaaccctgattatttaaattttcagCCAACACTTGTCACTACTTTCTGTTATGGTGTTCAATGCTTCTCGAGATACCCAGATCATATGAAACGGCATGACTTTTTCAAGAGTGCCATGCCCGAAGGTTATGTACAGGAAAGAACTATATTTTTCAAAGATGACGGGAACTACAAGACACgtaagtttaaacagttcggtactaactaaccatacatatttaaattttcagGTGCTGAAGTCAAGTTTGAAGGTGATACCCTTGTTAATAGAATCGAGTTAAAAGGTATTGATTTTAAAGAAGATGGAAACATTCTTGGACACAAATTGGAATACAACTATAACTCACACAATGTATACATCATGGCAGACAAACAAAAGAATGGAATCAAAGTTgtaagtttaaacatgattttactaactaactaatctaatttaaattttcagAACTTCAAAATTAGACACAACATTGAAGATGGAAGCGTTCAACTAGCAGACCATTATCAACAAAATACTCCAATTGGCGATGGCCCTGTCCTTTTACCAGACAACCATTACCTGTCCACACAATCTGCCCTTTCGAAAGATCCCAACGAAAAGAGAGACCACATGGTCCTTCTTGAGTTTGTAACAGCTGCTGGGATTACACATGGCATGGATGAACTATACAAATGCCCGGGGGATCGGTGGAGCTCCACCGGTGGCGGCCGCTCTAGAGAGAATCTTTATTTTCAGGGCGCCGCCAAATTCAAAGAAACCGCTGCTGCTAAATTCGAACGCCAGCACATGGACAGCGGAGGTGGAGGTATGGCGACCATCGTGCCATGCTCACTTGAGAAAGAAGAAGGAGCACCATCAGGACCTCGTCGGCTTCAAACTGAGATCGACGTGGACGCCAACGACAGCGGAAACGAGCTGTCGATGGGCGGAAGCAGCAGTG

(4d) *lin-41(tn1892)*

attgttccattcgttctgaaaagtcaaaaaattcataagtattctaattgtagagtcatcgtttgccctttccactgataattaatcaaccttttcagacttggaaaaagtgaaATGGTCTCCAAGGGAGAGGCCGTCATCAAGGAGTTCATGCGTTTCAAGGTCCACATGGAGGGATCCATGAACGGACACGAGTTCGAGATCGAGGGAGAGGGAGAGGGACGTCCATACGAGGGAACCCAAACCGCCAAGCTCAAGGTCACCAAGgtaagtttaaacatatatatactaactaaccctgattatttaaattttcagGGAGGACCACTCCCATTCTCCTGGGACATCCTCTCCCCACAATTCATGTACGGATCCCGTGCCTTCATCAAGCACCCAGCCGACATCCCAGACTACTACAAGCAATCCTTCCCAGAGGGATTCAAGTGGGAGCGTGTCATGAACTTCGAGGACGGAGGAGCCGTCACCGTCACCCAAGACACCTCCCTCGAGGACGGAACCCTCATCTACAAGgtaagtttaaacagttcggtactaactaaccatacatatttaaattttcagGTCAAGCTCCGTGGAACCAACTTCCCACCAGACGGACCAGTCATGCAAAAGAAGACCATGGGATGGGAGGCCTCCACCGAGCGTCTCTACCCAGAGGACGGAGTCCTCAAGGGAGACATCAAGATGGCCCTCCGTCTCAAGGACGGAGGACGTTACCTCGCCGACTTCAAGgtaagtttaaacatgattttactaactaactaatctgatttaaattttcagACCACCTACAAGGCCAAGAAGCCAGTCCAAATGCCAGGAGCCTACAACGTCGACCGTAAGCTCGACATCACCTCCCACAACGAGGACTACACCGTCGTCGAGCAATACGAGCGTTCCGAGGGACGTCACTCCACCGGAGGAATGGACGAGCTCTACAAGGAGAATCTGTACTTTCAATCCGGAAAGgtaagtttaaaataacttcgtatagcatacattatacgaagttattttcagGGAGCCGGATCTGATTATAAAGACGATGACGATAAGCGTGACTACAAGGACGACGACGACAAGCGTGATTACAAGGATGACGATGACAAGAGAATGGCGACCATCGTGCCATGCTCATTAGAGAAAGAAGAAGGAGCACCATCAGGACCTCGTCGGCTTCAAACTGAGATCGACGTGGACGCCAACGACAGCGGAAACGAGCTGTCGATGGGCGGAAGC

(4e) *lin-41(tn1894)*

attgttccattcgttctgaaaagtcaaaaaattcataagtattctaattgtagagtcatcgtttgccctttccactgataattaatcaaccttttcagacttggaaaaagtgaaATGAACCTTATTAAGGAAGATATGAGAGTCAAAGTTCATATGGAAGGAAACGTCAACGGTCATGCATTTGTTATTGAAGGAGAAGGAAAAGGAAAGCCATACGAAGGAACTCAAACTGCAAACTTGACTGTCAAAGAAGGAGCACCACTACCATTTAGTTACgtaagtttaaacatatatatactaactaaccctgattatttaaattttcagGATATTCTAACTACTGCCGTCCATTACGGAAACAGAGTTTTTACTAAATACCCAGAAGATATTCCTGATTACTTCAAGCAATCGTTTCCAGAAGGATACTCGTGGGAAAGAACTATGACTTTCGAAGATAAAGGTATTTGCACTATTgtaagtttaaacagttcggtactaactaaccatacatatttaaattttcagAGAAGTGATATTAGTCTAGAAGGTGATTGCTTCTTCCAAAATGTCAGATTTAAAGGAACTAACTTTCCTCCTAACGGACCAGTTATGCAAAAGAAGACTCTTAAGTGGGAACCATCGACTGAAAAACTACATGTTAGAGATGGACTACTTGTTGGAgtaagtttaaacttggacttactaactaacggattatatttaaattttcagAACATTAACATGGCACTACTACTAGAAGGTGGAGGTCACTACCTTTGCGATTTTAAAACTACTTACAAAGCAAAGAAGGTCGTCCAACTTCCAGATGCACACTTTGTTGATCACAGAATTGAAATACTAGGAAACGATTCGGATTACAACAAAGTTAAGCTATACGAACACGCAGTTGCAAGATACAGTCCTCTACCAAGTCAAGCATGGGAGAATCTGTACTTTCAATCCGGAAAGgtaagtttaaaataacttcgtatagcatacattatacgaagttattttcagGGAGCCGGATCTGATTATAAAGACGATGACGATAAGCGTGACTACAAGGACGACGACGACAAGCGTGATTACAAGGATGACGATGACAAGAGAATGGCGACCATCGTGCCATGCTCATTAGAGAAAGAAGAAGGAGCACCATCAGGACCTCGTCGGCTTCAAACTGAGATCGACGTGGACGCCAACGACAGCGGAAACGAGCTGTCGATGGGCGGAAGCAGCAGTGAAGgtaacacttga

(4f) *lin-41(tn2054)* and *lin-41(tn2055)*

attgttccattcgttctgaaaagtcaaaaaattcataagtattctaattgtagagtcatcgtttgccctttccactgataattaatcaaccttttcagacttggaaaaagtgaaATGAGTAAAGGAGAAGAATTGTTCACTGGAGTTGTCCCAATCCTCGTCGAGCTCGACGGAGACGTCAACGGACACAAGTTCTCCGTCTCCGGAGAGGGAGAGGGAGACGCCACCTACGGAAAGCTCACCCTCAAGTTCATCTGCACCACCGGAAAGCTCCCAGTCCCATGGCCAACCCTCGTCACCACCTTCTGCTACGGAGTCCAATGCTTCTCCCGTTACCCAGACCACATGAAGCGTCACGACTTCTTCAAGTCCGCCATGCCAGAGGGATACGTCCAAGAGCGTACCATCTTCTTCAAGgtaagtttaaacatatatatactaactactgattatttaaattttcagGACGACGGAAACTACAAGACCCGTGCCGAGGTCAAGTTCGAGGGAGACACCCTCGTCAACCGTATCGAGCTCAAGgtaagtttaaacagttcggtactaactaaccatacatatttaaattttcagGGAATCGACTTCAAGGAGGACGGAAACATCCTCGGACACAAGCTCGAGTACAACTACAACTCCCACAACGTCTACATCATGGCCGACAAGCAAAAGAACGGAATCAAGGTCAACTTCAAGgtaagtttaaacatgattttactaactaactaatctgatttaaattttcagATCCGTCACAACATCGAGGACGGATCCGTCCAACTCGCCGACCACTACCAACAAAACACCCCAATCGGAGACGGACCAGTCCTCCTCCCAGACAACCACTACCTCTCCACCCAATCCGCCCTCTCCAAGGACCCAAACGAGAAGCGTGACCACATGGTCCTCCTCGAGTTCGTCACCGCCGCCGGAATCACCCACGGAATGGACGAGCTCTACAAGGAGAATCTGTACTTTCAATCCGGAAAGgtaagtttaaaataacttcgtatagcatacattatacgaagttattttcagGGAGCCGGATCTGATTATAAAGACGATGACGATAAGCGTGACTACAAGGACGACGACGACAAGCGTGATTACAAGGATGACGATGACAAGAGAATGGCGACCATCGTGCCATGCTCATTAGAGAAAGAAGAAGGAGCACCATCAGGACCTCGTCGGCTTCAAACTGAGATCGACGTGGACGCCAACGACAGCGGAAACGAGCTG

(4g) *puf-3(tn1820)*

TCGGAGATGTACGGCATGTGGCTCGAGAAGATTCACGGACGAGTGATGCGAAACGCCCACCGCCTCGAAAGATTTTCGTCGGGCAAGAAGATTATCGAAGCGCTCCAATCAATGTCATTGTACGGAGCATCGGGAGCCTCAGGAGCATCGATGAGTAAAGGAGAAGAATTGTTCACTGGAGTTGTCCCAATCCTCGTCGAGCTCGACGGAGACGTCAACGGACACAAGTTCTCCGTCTCCGGAGAGGGAGAGGGAGACGCCACCTACGGAAAGCTCACCCTCAAGTTCATCTGCACCACCGGAAAGCTCCCAGTCCCATGGCCAACCCTCGTCACCACCTTCTGCTACGGAGTCCAATGCTTCTCCCGTTACCCAGACCACATGAAGCGTCACGACTTCTTCAAGTCCGCCATGCCAGAGGGATACGTCCAAGAGCGTACCATCTTCTTCAAGgtaagtttaaacatatatatactaactactgattatttaaattttcagGACGACGGAAACTACAAGACCCGTGCCGAGGTCAAGTTCGAGGGAGACACCCTCGTCAACCGTATCGAGCTCAAGgtaagtttaaacagttcggtactaactaaccatacatatttaaattttcagGGAATCGACTTCAAGGAGGACGGAAACATCCTCGGACACAAGCTCGAGTACAACTACAACTCCCACAACGTCTACATCATGGCCGACAAGCAAAAGAACGGAATCAAGGTCAACTTCAAGgtaagtttaaacatgattttactaactaactaatctgatttaaattttcagATCCGTCACAACATCGAGGACGGATCCGTCCAACTCGCCGACCACTACCAACAAAACACCCCAATCGGAGACGGACCAGTCCTCCTCCCAGACAACCACTACCTCTCCACCCAATCCGCCCTCTCCAAGGACCCAAACGAGAAGCGTGACCACATGGTCCTCCTCGAGTTCGTCACCGCCGCCGGAATCACCCACGGAATGGACGAGCTCTACAAGGAGAATCTGTACTTTCAATCCGGAAAGgtaagtttaaaataacttcgtatagcatacattatacgaagttattttcagGGAGCCGGATCTGATTATAAAGACGATGACGATAAGCGTGACTACAAGGACGACGACGACAAGCGTGATTACAAGGATGACGATGACAAGAGATAGatacacataggcaatttttattcatttccatttgaatcctaacccccaccatcacccaactcacgtgtatactattattatttttattattatattgctagaaaattgtagaaagagtattgcaacctcttctatattatat

(4h) *puf-11(tn1824)*

ATGTACGGCATGTGGCTCGAGAAGATTCGCGAACGAGTCATGCGAAACGCCAATCGCCTCGAAAGATTCTCGTCGGGCAAGAAGATAATCGAAGCGCTCCAATCAATGTCATTTTACGGAGCATCGGGAGCCTCAGGAGCATCGATGAGTAAAGGAGAAGAATTGTTCACTGGAGTTGTCCCAATCCTCGTCGAGCTCGACGGAGACGTCAACGGACACAAGTTCTCCGTCTCCGGAGAGGGAGAGGGAGACGCCACCTACGGAAAGCTCACCCTCAAGTTCATCTGCACCACCGGAAAGCTCCCAGTCCCATGGCCAACCCTCGTCACCACCTTCTGCTACGGAGTCCAATGCTTCTCCCGTTACCCAGACCACATGAAGCGTCACGACTTCTTCAAGTCCGCCATGCCAGAGGGATACGTCCAAGAGCGTACCATCTTCTTCAAGgtaagtttaaacatatatatactaactactgattatttaaattttcagGACGACGGAAACTACAAGACCCGTGCCGAGGTCAAGTTCGAGGGAGACACCCTCGTCAACCGTATCGAGCTCAAGgtaagtttaaacagttcggtactaactaaccatacatatttaaattttcagGGAATCGACTTCAAGGAGGACGGAAACATCCTCGGACACAAGCTCGAGTACAACTACAACTCCCACAACGTCTACATCATGGCCGACAAGCAAAAGAACGGAATCAAGGTCAACTTCAAGgtaagtttaaacatgattttactaactaactaatctgatttaaattttcagATCCGTCACAACATCGAGGACGGATCCGTCCAACTCGCCGACCACTACCAACAAAACACCCCAATCGGAGACGGACCAGTCCTCCTCCCAGACAACCACTACCTCTCCACCCAATCCGCCCTCTCCAAGGACCCAAACGAGAAGCGTGACCACATGGTCCTCCTCGAGTTCGTCACCGCCGCCGGAATCACCCACGGAATGGACGAGCTCTACAAGGAGAATCTGTACTTTCAATCCGGAAAGgtaagtttaaaataacttcgtatagcatacattatacgaagttattttcagGGAGCCGGATCTGATTATAAAGACGATGACGATAAGCGTGACTACAAGGACGACGACGACAAGCGTGATTACAAGGATGACGATGACAAGAGATAGatataggcaattcatttgcattattatcctaacccccactcacgtgtattattattattattattattattatattgctagaaaacattagaaagagtattgcaacctcttccatgttatatgaagaccccggcccctttttgcacaa

(4i) *cyb-1(tn1806)*

cataattatttgcagAATAAGTATCAATCAAGCAAGCTTGCACAAGTTTCCAACTTGATGACCGACGACGTACTCGAGAAAATCAATCGGATGGGCCAGAATGCAAAAGTAGACGCATCAGAAATGGAAGGAGCATCGGGAGCCTCAGGAGCATCGATGAGTAAAGGAGAAGAATTGTTCACTGGAGTTGTCCCAATCCTCGTCGAGCTCGACGGAGACGTCAACGGACACAAGTTCTCCGTCTCCGGAGAGGGAGAGGGAGACGCCACCTACGGAAAGCTCACCCTCAAGTTCATCTGCACCACCGGAAAGCTCCCAGTCCCATGGCCAACCCTCGTCACCACCTTCTGCTACGGAGTCCAATGCTTCTCCCGTTACCCAGACCACATGAAGCGTCACGACTTCTTCAAGTCCGCCATGCCAGAGGGATACGTCCAAGAGCGTACCATCTTCTTCAAGgtaagtttaaacatatatatactaactactgattatttaaattttcagGACGACGGAAACTACAAGACCCGTGCCGAGGTCAAGTTCGAGGGAGACACCCTCGTCAACCGTATCGAGCTCAAGgtaagtttaaacagttcggtactaactaaccatacatatttaaattttcagGGAATCGACTTCAAGGAGGACGGAAACATCCTCGGACACAAGCTCGAGTACAACTACAACTCCCACAACGTCTACATCATGGCCGACAAGCAAAAGAACGGAATCAAGGTCAACTTCAAGgtaagtttaaacatgattttactaactaactaatctgatttaaattttcagATCCGTCACAACATCGAGGACGGATCCGTCCAACTCGCCGACCACTACCAACAAAACACCCCAATCGGAGACGGACCAGTCCTCCTCCCAGACAACCACTACCTCTCCACCCAATCCGCCCTCTCCAAGGACCCAAACGAGAAGCGTGACCACATGGTCCTCCTCGAGTTCGTCACCGCCGCCGGAATCACCCACGGAATGGACGAGCTCTACAAGGAGAATCTGTACTTTCAATCCGGAAAGgtaagtttaaaataacttcgtatagcatacattatacgaagttattttcagGGAGCCGGATCTGATTATAAAGACGATGACGATAAGCGTGACTACAAGGACGACGACGACAAGCGTGATTACAAGGATGACGATGACAAGAGATGAgcacttcgatccgaaataaacattttgtaccattcagtctttcttgattcttatgtttcttccttcattgcttttcttttcggctcctttccattttttaactcctccagtcgtttccacgtcaatctggtcacac

(4j) *rnp-8(tn1860)*

GACGAGAAGATGCTGCTCGACGAGGCGTCTTCAATCATCGAAAATACAACTCCAGCAGTGTCTACTGCTCCGGCTGCTGCTCCAGGAGCTACAATGCTCCAAATAGGAGCATCGGGAGCCTCAGGAGCATCGATGAGTAAAGGAGAAGAATTGTTCACTGGAGTTGTCCCAATCCTCGTCGAGCTCGACGGAGACGTCAACGGACACAAGTTCTCCGTCTCCGGAGAGGGAGAGGGAGACGCCACCTACGGAAAGCTCACCCTCAAGTTCATCTGCACCACCGGAAAGCTCCCAGTCCCATGGCCAACCCTCGTCACCACCTTCTGCTACGGAGTCCAATGCTTCTCCCGTTACCCAGACCACATGAAGCGTCACGACTTCTTCAAGTCCGCCATGCCAGAGGGATACGTCCAAGAGCGTACCATCTTCTTCAAGgtaagtttaaacatatatatactaactactgattatttaaattttcagGACGACGGAAACTACAAGACCCGTGCCGAGGTCAAGTTCGAGGGAGACACCCTCGTCAACCGTATCGAGCTCAAGgtaagtttaaacagttcggtactaactaaccatacatatttaaattttcagGGAATCGACTTCAAGGAGGACGGAAACATCCTCGGACACAAGCTCGAGTACAACTACAACTCCCACAACGTCTACATCATGGCCGACAAGCAAAAGAACGGAATCAAGGTCAACTTCAAGgtaagtttaaacatgattttactaactaactaatctgatttaaattttcagATCCGTCACAACATCGAGGACGGATCCGTCCAACTCGCCGACCACTACCAACAAAACACCCCAATCGGAGACGGACCAGTCCTCCTCCCAGACAACCACTACCTCTCCACCCAATCCGCCCTCTCCAAGGACCCAAACGAGAAGCGTGACCACATGGTCCTCCTCGAGTTCGTCACCGCCGCCGGAATCACCCACGGAATGGACGAGCTCTACAAGGAGAATCTGTACTTTCAATCCGGAAAGgtaagtttaaaataacttcgtatagcatacattatacgaagttattttcagGGAGCCGGATCTGATTATAAAGACGATGACGATAAGCGTGACTACAAGGACGACGACGACAAGCGTGATTACAAGGATGACGATGACAAGAGATAGgagaagatcacatatacaataatataatcttattgcattttcgcaattctcgttctctccacacacatacacacatcatcccaagtattcctgtgctgaatctcagtttgaatgatgtttcataccgtttttatcccactattgccttatcgtttcctgt

(4k) *etc-1(tn1919)*

tgaataagtgttaaatgtaaacactcagtatttatttttgttttgcaattgaatatttcagatttccactaattagtttccaattttagATGAGTAAAGGAGAAGAATTGTTCACTGGAGTTGTCCCAATCCTCGTCGAGCTCGACGGAGACGTCAACGGACACAAGTTCTCCGTCTCCGGAGAGGGAGAGGGAGACGCCACCTACGGAAAGCTCACCCTCAAGTTCATCTGCACCACCGGAAAGCTCCCAGTCCCATGGCCAACCCTCGTCACCACCTTCTGCTACGGAGTCCAATGCTTCTCCCGTTACCCAGACCACATGAAGCGTCACGACTTCTTCAAGTCCGCCATGCCAGAGGGATACGTCCAAGAGCGTACCATCTTCTTCAAGgtaagtttaaacatatatatactaactactgattatttaaattttcagGACGACGGAAACTACAAGACCCGTGCCGAGGTCAAGTTCGAGGGAGACACCCTCGTCAACCGTATCGAGCTCAAGgtaagtttaaacagttcggtactaactaaccatacatatttaaattttcagGGAATCGACTTCAAGGAGGACGGAAACATCCTCGGACACAAGCTCGAGTACAACTACAACTCCCACAACGTCTACATCATGGCCGACAAGCAAAAGAACGGAATCAAGGTCAACTTCAAGgtaagtttaaacatgattttactaactaactaatctgatttaaattttcagATCCGTCACAACATCGAGGACGGATCCGTCCAACTCGCCGACCACTACCAACAAAACACCCCAATCGGAGACGGACCAGTCCTCCTCCCAGACAACCACTACCTCTCCACCCAATCCGCCCTCTCCAAGGACCCAAACGAGAAGCGTGACCACATGGTCCTCCTCGAGTTCGTCACCGCCGCCGGAATCACCCACGGAATGGACGAGCTCTACAAGGAGAATCTGTACTTTCAATCCGGAAAGgtaagtttaaaataacttcgtatagcatacattatacgaagttattttcagGGAGCCGGATCTGATTATAAAGACGATGACGATAAGCGTGACTACAAGGACGACGACGACAAGCGTGATTACAAGGATGACGATGACAAGAGAATGTCATTCAACCTGGTATTTTTTAACGGTGATGTTCGGAGGGACAAAAAACAGGAAGCAATGATCAAAAGCCTAAACGAGCGGGACAACTTTCTTAACAGACTCCGCGAAATGGAACATAAACGGGAAAAGGATG

(4l) *etc-1(tn1920)*

tcagatttccactaattagtttccaattttagATGTCATTCAACCTGGTATTTTTTAACGGTGATGTTCGGAGGGACAAAAAACAGGAAGCAATGAGTAAAGGAGAAGAATTGTTCACTGGAGTTGTCCCAATCCTCGTCGAGCTCGACGGAGACGTCAACGGACACAAGTTCTCCGTCTCCGGAGAGGGAGAGGGAGACGCCACCTACGGAAAGCTCACCCTCAAGTTCATCTGCACCACCGGAAAGCTCCCAGTCCCATGGCCAACCCTCGTCACCACCTTCTGCTACGGAGTCCAATGCTTCTCCCGTTACCCAGACCACATGAAGCGTCACGACTTCTTCAAGTCCGCCATGCCAGAGGGATACGTCCAAGAGCGTACCATCTTCTTCAAGgtaagtttaaacatatatatactaactactgattatttaaattttcagGACGACGGAAACTACAAGACCCGTGCCGAGGTCAAGTTCGAGGGAGACACCCTCGTCAACCGTATCGAGCTCAAGgtaagtttaaacagttcggtactaactaaccatacatatttaaattttcagGGAATCGACTTCAAGGAGGACGGAAACATCCTCGGACACAAGCTCGAGTACAACTACAACTCCCACAACGTCTACATCATGGCCGACAAGCAAAAGAACGGAATCAAGGTCAACTTCAAGgtaagtttaaacatgattttactaactaactaatctgatttaaattttcagATCCGTCACAACATCGAGGACGGATCCGTCCAACTCGCCGACCACTACCAACAAAACACCCCAATCGGAGACGGACCAGTCCTCCTCCCAGACAACCACTACCTCTCCACCCAATCCGCCCTCTCCAAGGACCCAAACGAGAAGCGTGACCACATGGTCCTCCTCGAGTTCGTCACCGCCGCCGGAATCACCCACGGAATGGACGAGCTCTACAAGGAGAATCTGTACTTTCAATCCGGAAAGgtaagtttaaaataacttcgtatagcatacattatacgaagttattttcagGGAGCCGGATCTGATTATAAAGACGATGACGATAAGCGTGACTACAAGGACGACGACGACAAGCGTGATTACAAGGATGACGATGACAAGAGAATGATCAAAAGCCTAAATGAACGAGACAACTTTCTTAACAGACTCCGCGAAATGGAACATAAACGGGAAAAGGATGAGAAGCAAGAAAAGGCTGCGAGAAAAGTTCAAAAGTTTTGGAGAGGACATCGAGTAC
